# Somatic repetitive element insertions Define New Biomarkers in Pan-cancer genome

**DOI:** 10.1101/2025.02.19.638966

**Authors:** Lin Xia, Kailing Tu, Huan Wang, Xuyan Liu, Yahui Liu, Qilin Zhang, Ziting Feng, Chunyan Yu, Maolin Li, Yan Huang, Jing Wang, Ranlei Wei, Tianfu Zeng, Xuenan Pi, Chen Ling, Qianwen Zhang, Yixin Qiao, Shengzhong Hou, Zhuoyuan Zhang, Ailin Wei, Haotian Liao, Wenhao Guo, Jifeng Liu, Zhong Wu, Zhoufeng Wang, Ang Li, Weiming Li, Dan Xie

## Abstract

Repetitive elements insertions (REIs), including tandem repeat insertions (TRIs) and transposable element insertions (TEIs), constitute >90% of large-scale (>50bp) somatic insertions and are linked to diseases and cancer biomarkers. However, the genomic landscape of large-scale REIs in pan-cancer has not been well-characterized due to short-read sequencing limitations. Here, we detected somatic REIs in 325 cancer genomes across 12 cancer types using long-read whole-genome sequencing and customized algorithms. The diverse sequence patterns of somatic TRIs and the insertion mechanisms of somatic TEIs were characterized across different cancer types. We identified 152 high-frequency somatic TRIs, with 10 showing pan-cancer characteristics, including TRIs in *SHROOM2, PALMD*, and the enhancer of *PTPRZ1*. The MHC class II gene cluster exhibited the highest somatic TEI abundance, affecting ∼40% tumors. Adenocarcinomas and squamous cell carcinomas exhibited differing TEI distributions within this region. Our study highlights REIs as important but under-explored markers for personalized cancer diagnosis and immunotherapy.

## Introduction

Cancer is formidable in its vast diversity, although all cancers share common hallmarks^1,2^. Understanding both the commonalities and variations among cancer types is crucial for uncovering the fundamental mechanisms of oncogenesis and advancing precision medicine. Pan-cancer studies, which aim at identifying shared genomic patterns and alterations driving cancer progression across cancers ^3–7^, not only enhances our understanding of cancer biology, but also holds considerable promise for advancing early diagnosis and developing personalized treatment strategies. These efforts have significantly expanded the knowledge of cancer driver genes, with *TP53*, *PIK3CA*, *KRAS*, *PTEN*, and *ARID1A* among the most frequently mutated genes across various cancers ^8^. Furthermore, pan-cancer studies have uncovered new non-coding driver genes ^9,10^, mutational signatures ^11^, and evolutionary patterns ^12–14^, as well as their downstream effects on gene expression ^15^ and cancer-specific characteristics ^16,17^. Despite these advances, many findings have yet to translate into clinical benefits, highlighting the pressing need for the identification of other critical genetic variants in cancer progression.

Previous pan-cancer research has predominantly focused on small-scale mutations, such as single nucleotide variations (SNVs) and small insertions/deletions (INDELs, < 50bp). Structural variations (SVs), which impact a broader range of nucleotide change ^18^, have increasingly garnered attention due to their significant role in cancer progression ^19^. With short-read sequencing (SRS) data, researchers have identified various genomic SVs, including whole-genome duplications ^20^, chromothripsis ^21^, chromosomal instability ^17^, and complex structural variants ^22^, all associated with poorer prognoses. Nevertheless, large-scale insertions (>50 bp) in tumors remain less explored, particularly for repetitive element insertions (REIs), including tandem repeat insertions (TRIs) and transposable element insertions (TEIs) ^23^. Previous population studies and tumor analyses indicate that over 80% of large-scale insertions are repetitive element insertions (REIs) ^24–26^. Both TRIs ^27,28^ and TEIs ^29–32^ are drawing increasing attention as potential diagnostic and therapeutic target for various types of cancer. Despite extensive mapping efforts to characterize TRI and TEI events in cancer genomes ^27,28,33–35^, only a limited number of recurrent somatic REIs have been identified. The short read lengths of SRS technologies have hindered the in-depth exploration of REI variation patterns in cancer genomes ^23^. These challenges underscore the urgent need for advancements in sequencing technologies and bioinformatics approaches to enhance the detection and analysis of somatic REIs in cancer.

The continuous advancement of third-generation sequencing (TGS) technology, particularly Oxford Nanopore Technologies (ONT) and Pacific Biosciences (PacBio), has significantly enhanced long-read sequencing (LRS). Technology of ONT produces reads exceeding 10 kb, with some extending to millions of bases, enabling the complete capture of large-scale SVs and their flanking regions in a single read. This improvement enhanced the sensitivity and accuracy of SV detection ^26,36–38^. LRS has enabled researchers to accurately map TR alternations across populations, revealing complex TR alternations that were previously undetectable by SRS ^39,40^. LRS were also used in population studies to uncover the widespread presence of short interspersed nuclear element (SINE) insertions, which are more difficult to capture than long interspersed nuclear element (LINE) insertions typically identified by SRS ^36,37^. However, most population studies have focused on germline genomic alterations, leaving the study of somatic REIs in tumor genome largely unexplored ^41–43^. While some studies have applied long-read sequencing to examine structural variations in tumors, the lack of large tumor sample cohorts and suitable bioinformatics tools has limited the comprehensive analyses.

In this study, we conducted long-read nanopore whole genome sequencing (LRS WGS) on 325 paired tumor and blood samples from 12 cancer types. With specialized algorithms developed, we identified 39,661 somatic REIs. The diversity of somatic TRIs sequence patterns and variable TEI insertion mechanisms across cancer types were elucidated. In total, we identified 152 high-frequency somatic TRIs, where 10/152 are shared in diverse cancer types, including somatic TRI events were located within the intron of *SHROOM2* and *PALMD*, and in the distal enhancer region upstream of *PTPRZ1*.

Regarding TEIs, approximately 40% of tumors harbored somatic TEIs in the MHC class II gene cluster, with adenocarcinomas showing greater TEI enrichment than squamous cell carcinomas. Notably, adenocarcinoma cells with a higher TEI burden within this region showed significantly enhanced T cell activation compared to those with a lower burden, indicating that TEIs may contribute to the regulation of tumor immune responses. Overall, this study provides a valuable resource to the scientific and clinical community and a paradigm for character the somatic genomic alternations in pan-cancer genomes using LRS.

## Results

### Section 1: Specimens, Tumor Types and Dataset

This pan-cancer study encompassed 325 tumor samples and 300 paired blood samples from 12 distinct cancer types (Figure 1a). Solid tumor types were from gynecologic (ovarian [OV], uterine corpus endometrial carcinoma [UCEC], cervical squamous cell carcinoma [CCA]), core gastrointestinal (stomach adenocarcinoma [GC], colon adenocarcinoma [IC]), developmental gastrointestinal (liver hepatocellular carcinoma [HCC], pancreatic adenocarcinoma [PAAD]), head and neck (Oral squamous cell carcinoma [OSCC], Laryngocarcinom [LC]), thoracic (lung adenocarcinoma [LUAD], lung squamous cell carcinoma [LUSC]), and central nervous system cancers (glioblastoma multiforme [GBM]). 4, 2, 27 and 2 multiple synchronous tumors were obtained from 9 patients with LUAD (2), GBM (1), HCC (5) and PAAD (1), separately (Table S1). All patients provided written informed consent for genomic studies approved by local institutional review boards.

**Figure 1.**
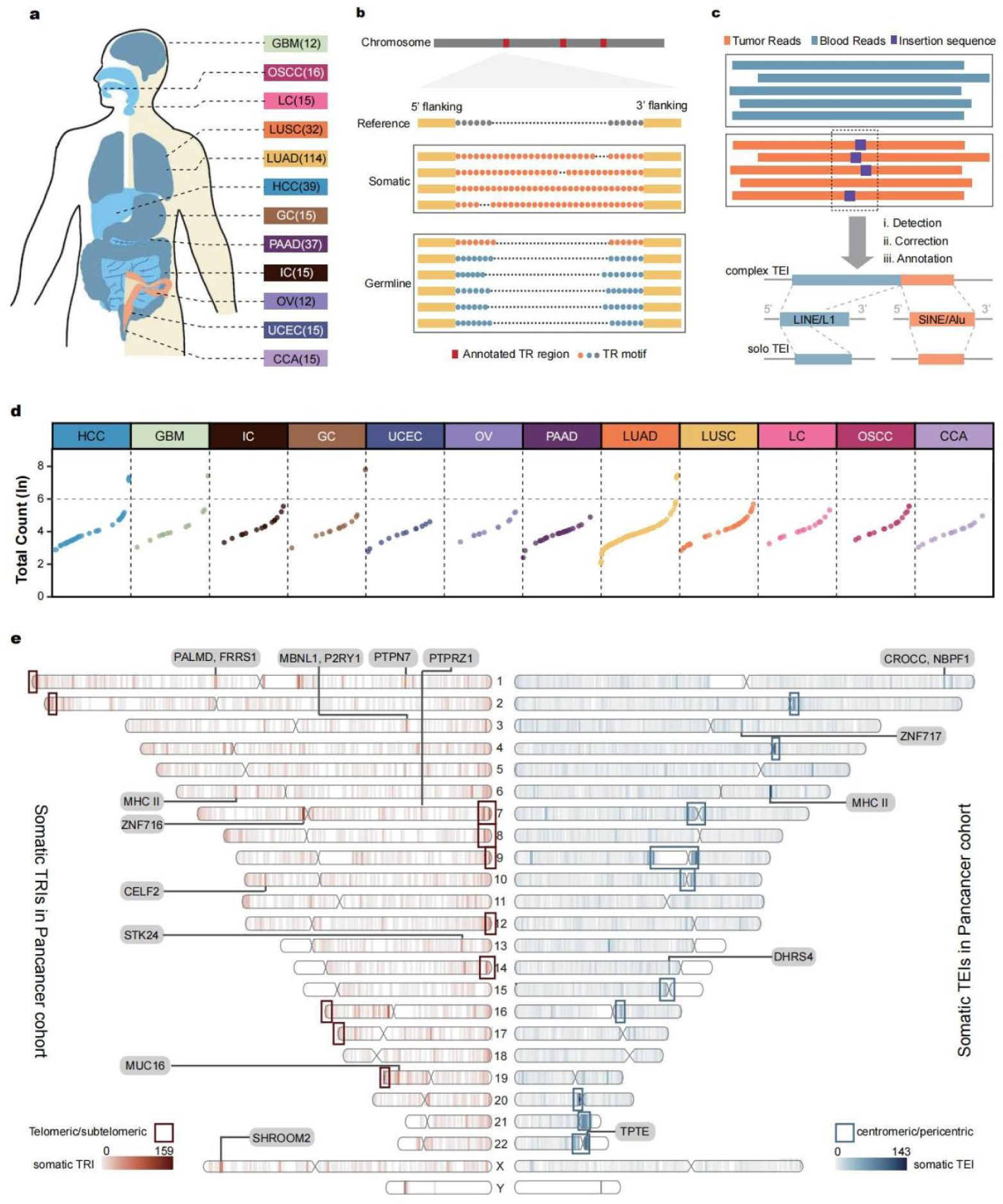
Genome-wide detection of somatic insertions in pan-cancer genomes. a. Overview of a pan-cancer cohort comprising 325 paired tumor and blood samples from 299 patients across 12 cancer types: 1) glioblastoma (GBM); 2) oral squamous cell carcinoma (OSCC); 3) laryngeal cancer (LC); 4) lung adenocarcinoma (LUAD); 5) lung squamous cell carcinoma (LUSC); 6) hepatocellular carcinoma (HCC); 7) gastric carcinoma (GC); 8) pancreatic adenocarcinoma (PAAD); 9) intestinal carcinoma (IC); 10) ovarian cancer (OV); 11) uterine corpus endometrial carcinoma (UCEC); 12) cervical squamous cell carcinoma (CCA). b. Scheme of the bioinformatics pipeline used to identify somatic TRIs. The top track (‘reference’) shows the hg38 genome. the second track (‘somatic’) shows reads alignments supporting somatic SVs mapped to the reference genome, and the third track (‘germline’) shows reads alignments supporting germline SVs mapped to the reference genome. Circles represent repeat motifs, with different colors indicating different repeat motifs. c. Scheme of the bioinformatics pipeline for identifying somatic TEI. The first blue track represents alignments from the tumor sample mapped to the reference genome, while the second orange track shows alignments from the paired blood sample. After initial detection (Method), inserted sequences are corrected. Following annotation (Method), somatic TEIs are classified into two categories: complex TEIs (upper track) or solo TEIs (lower track). d. Somatic insertion frequency (ln) across cancer types. Somatic insertion frequencies were calculated based on the total number of somatic TRIs and TEIs per sample. e. Ideogram depicting the distribution of somatic TEI (blue, left) and somatic TRI (red, right). The ideogram shows the genomic distribution of somatic TEIs (blue) and somatic TRIs (red), with darker colors indicating higher frequencies of TEIs or TRIs within each 1Mb window. The top 30 high-frequency 1Mb windows are highlighted according to the following features: regions with clustered TEs near characteristic genes are marked with a gray background; high-frequency windows located in telomeric, sub-telomeric, centromeric regions are enclosed in a box.

Whole-genome sequencing (WGS) was performed on all tumor and paired blood samples using PromethION nanopore sequencer, yielding long sequencing reads (Table S2). A total of 45.73 terabases (Tb) of long clean reads were generated, averaging 93.03 gigabases (Gb) per tumor sample and 51.63 Gb per paired blood sample. The mean read depth reached 30 X (13 X∼46 X) for tumor samples and 17 X (3 X∼43 X) for paired blood samples (Figure S1a), with mean read N50 length across all samples was 24.9 kbp (4.5 kbp∼43.1 kbp) for clean long reads (Figure S1b). Due to an update in ONT technology during data acquisition, two versions of sequencing chips were employed in this study. Specifically, 83.06% (510/614) of the LRS WGS data were generated using the R9 chips, while the remaining data were obtained using the R10 chips (Figure S1c). Notably, the data produced with the R10 chips exhibited significantly higher quality compared to that generated with the R9 chips (Wilcoxon test, p < 2.2e-16) (Figure S1d). Additionally, RNA-seq datasets were generated for 89 NSCLC samples on an NGS platform ^25^ (Table S3).

### Section 2: Accurate identification of somatic REIs in pan-cancer genomes

In our previous research, nearly 90% somatic insertions in NSCLC genomes were identified as somatic TRIs and TEIs, where somatic TRIs were predominantly located within annotated simple repeat regions (77.8%) ^25^. The high repetitiveness of these sequences complicates alignment accuracy, often resulting in a high false-positive rate. To address this issue, we designed a more precise algorithm for detecting somatic TRIs across tumor genomes (STAR Methods) (Figure 1b and S2a). First, cleaned LRS reads were mapped to the hg38 reference genome using Minimap2 software (version 2.17) ^44^], achieving an average mapping rate of 99.91% (93.07%-100%). Next, widely used SV detection softwares were applied to identify loci harboring somatic TRIs. Specifically, Sniffles2 (version 2.0.6) ^45^ was employed to detect INSs within tandem repeat regions annotated by RepeatMasker in the hg38 reference genome. Repeat expansions (REs) in tumor genomes were then identified using Straglr (version 1.4.1) ^46^. Regions identified by either tool were combined to form a final list of candidate somatic tandem repeat insertion windows for further analysis. For each paired tumor-blood samples, somatic TRIs within these candidate windows were detected based on the clustering of reads according to editing distance and the TDScope algorithm ^47^ (Figure S2a, STAR Methods).

Somatic transposable element insertions (TEIs) were identified by optimizing an in-house pipeline for somatic structural variant (SV) detection ^25^ (Figure 1c and S2b). First, tumor sample alignments were merged with those of the paired blood sample. Next, large-scale INSs were detected using Sniffles2 (version 2.0.6) ^45^. A somatic INS was considered valid if more than 3 supporting reads from the tumor sample, but no read from the paired blood sample (STAR Methods). Next, Iris software (version 1.0.5)^48^ was used to correct the sequences and breakpoints of the candidate somatic INSs. To false positives due to reference-individual genome differences, a custom pipeline was developed. The reference genome at each INS loci was polished using Racon (version 1.4.16)^49^, based on alignments from the paired blood samples. Tumor read were then realigned to the polished reference, and those with more than 3supporting reads were retained as candidate somatic INSs. Finally, consensus sequences of somatic INSs were categorized using RepeatMasker (version 4.1.2)^50^, Tandem Repeat Finder (version 4.10.0) ^51^, and Sdust (version 0.1)^52^ (STAR Methods). Somatic INSs with consensus sequences containing TE segments were classified as somatic TEIs (STAR Methods).

A total of 16,382 somatic TRIs (Table S4) were identified across 325 tumor genomes, with counts ranging from 1 to 849, and an average of 50. Similarly, 23,279 somatic TEIs (Table S5) were identified across 325 tumor genomes, with counts ranging from 0 to 1,776, and an average of 72 (Figure 1d and S2c). No correlation was identified between tumor purity (mean 46.33%, Table S1) and the count of somatic TRIs or somatic TEIs (Figure S3a-S3d, STAR Methods). Squamous carcinomas (SCC) exhibited a significantly higher burden of somatic TEIs compared to other cancer types (Welch two sample t-test, p value=0.0014 with Other, p value=1.6e-05 with Adenocarcinoma, p value=2.9e-07 with high burden) (Figure S3e and S3f). Notably, 75.88% (3351/4416) of the somatic TEIs were specific to SCC. However, only 43of the 3351 of the SCC-specific somatic TEIs were shared between two SCC samples, suggesting that while TEs are more active in SCC genomes, but SCC-specific TEIs were mostly low-frequency mutations. The distribution of somatic TRI and TEI counts across various cancer types did not show significant differences (Figure 1c, S3g).

Interestingly, a subset of 11 non-SCC tumor samples exhibited markedly higher numbers of both somatic TRIs (ranging from 592 to 849) and somatic TEIs (ranging from 621 to 1776) compared to the remaining tumors (TRIs: ranging from 1 to 139, TEIs: ranging from 0 to 279) (Figure 1d and S2c). This group, termed high-burden samples, included 2 HCC samples, 1 GBM sample, 2 GC samples, and 6 LUAD samples. Using the Genomic Regions Enrichment of Annotations Tool (GREAT)^53^, the high-burden-specific somatic TRIs (Table S6) were linked to distal cis-regulatory effects on genes involved in cell cycle regulation, telomere length, and chromosome localization (Figure S3h). Additionally, transcriptomic analysis of available NSCLC NGS RNA-seq data revealed that high-burden LUAD samples exhibited significantly lower expression of DNA repair-related genes, including *ERCC3*, *RAD51*, and *XRCC4* (Wilcoxon rank test, p < 0.05, |log2(FoldChange)| > 1), compared to other LUAD samples (Figure S3i). Reduced expression of these DNA repair genes may lead to the accumulation of unrepaired DNA damage, contributing to genomic instability ^54–56^.

Somatic TRIs were significantly enriched in subtelomeric regions (5.308-fold compared to the rest region of chromosome arm in average, Student t-test, p-value = 9.81e-14), whereas no enrichment of somatic TEIs was detected in subtelomeric regions (Student t-test, p-value = 0.6275) (Figure S4a, Table S7). Similar to the previous LRS study ^57^, the long arms of human chromosomes were more likely exhibited a broader zone of somatic TRIs enrichment (5–10 Mbp) compared to the short arms (Figure S4b). Similar to a large healthy Chinese cohort genomic study ^58^ and a recent TE recombination study ^59^, somatic TEIs were frequently enriched in centromeric and pericentromeric regions, particularly on chromosomes 20, 4, and 9 (Figure S4c). Notably, solo LINE insertions, solo SINE insertions, and complex TEIs containing LTR segments exhibited the most pronounced enrichment in these regions. (Figure S4c). In contrast, a comparable analysis of TEs in the reference genome and all somatic TEIs revealed no such patterns for solo LINE insertions.

To identify hotspot regions of somatic TRIs and TEIs, the genome was segmented into non-overlapping 1 Mb windows, and their distribution was investigated (Figure 1e). Hotspot regions containing genes that overlapped or were adjacent to somatic TRIs and TEIs were identified. These included *MUC16*, associated with various cancers ^60–62^ and widely used as a biomarker for ovarian cancer (CA125) ^63^ *STK24*, a prognostic marker for HCC, LUAD, PAAD, and KIRC (p < 0.001, log-rank test), involved in apoptosis regulation and cell migration^64,65^; *NBPF1*, part of the neuroblastoma breakpoint family, linked to genomic instability in neuroblastoma and gliomas^66,67^; and *CELF2*, a prognostic marker for HNSC and KIRC (p < 0.001, log-rank test), implicated in apoptosis and cancer progression ^68^. Genes such as *ZNF716* and *ZNF717*, which encoding zinc finger proteins involved in transcriptional regulation and signaling ^69^; *PTPN7* and *PTPRZ1*, which were members of the protein tyrosine phosphatase family ^70,71^, were also identified as highly recurrent but whose role in cancer remain to be fully characterized. In addition to genes with established roles in tumor progression, we also identified a number of genes with high-frequency somatic REI events across pan-cancer samples, including those involved in cytoskeleton regulation (e.g., *SHROOM2*, *PALMD*, and *CROCC*), suggesting their potential as biomarkers.

We next examined the enrichment of somatic TRIs and somatic TEIs in gene and regulatory regions by comparing the real dataset (‘O’, observed) to datasets representing the genomic coordinates of gene and regulatory regions (‘E’, expected) (Figure S4d). 52.5% somatic TRIs were located within gene regions, while 47.9% somatic TEIs were detected in these regions. Notably, somatic TRIs were significantly enriched in candidate cis-regulatory elements predicted by ENCODE (cCRE)) (log2(O/E) ranging from 0.27 to 2.07, binomial test, FDR < 0.05), suggesting their potential roles in chromosomal structure and genome regulation. In contrast, somatic TEIs were enriched in regions proximal to genebody (±2.5 kb, log2(O/E) ranging from 0.1 to 0.2, binomial test, FDR < 0.05), implicating them in noncanonical splicing events, as previously reported ^31^.

### Section 3: The Sequence Features and Mechanisms of Somatic REIs

To explore the mutational patterns of somatic TRI, we applied pairwise alignment algorithms to compare the consensus sequences of germline and somatic events within the somatic TRI windows across various samples. Sequences identified exclusively in somatic TRI events, relative to their paired germline events, were classified as somatic TRI mutational sequences. These sequences were classified into three categories based on the repeat motifs involved. The first category, ‘Expansion’, referred to somatic TRI events where a pre-existing germline repeat motif is expanded in the somatic sample (Figure 2a). For example, in the somatic TRI window (chr18:80,195,548-80,195,600) of sample UCEC12, the AT motif was notably expanded (Figure S5a), reflecting an increase in repeat length. The second category, ‘Motif-Change’, involved somatic TRI events where a new repeat motif emerges, absent in the germline sequence (Figure 2a). In sample Lung136, the somatic sequence showed the appearance of the AGATATATAT motif, which was not present in the germline TR window (chrX:145,918,007-145,918,062) (Figure S5a). The third category includes somatic TRI events where the inserted sequence did not match any recognizable repeat motif, classified as ‘Undefined’ (Figure 2a and S5a).

**Figure 2.**
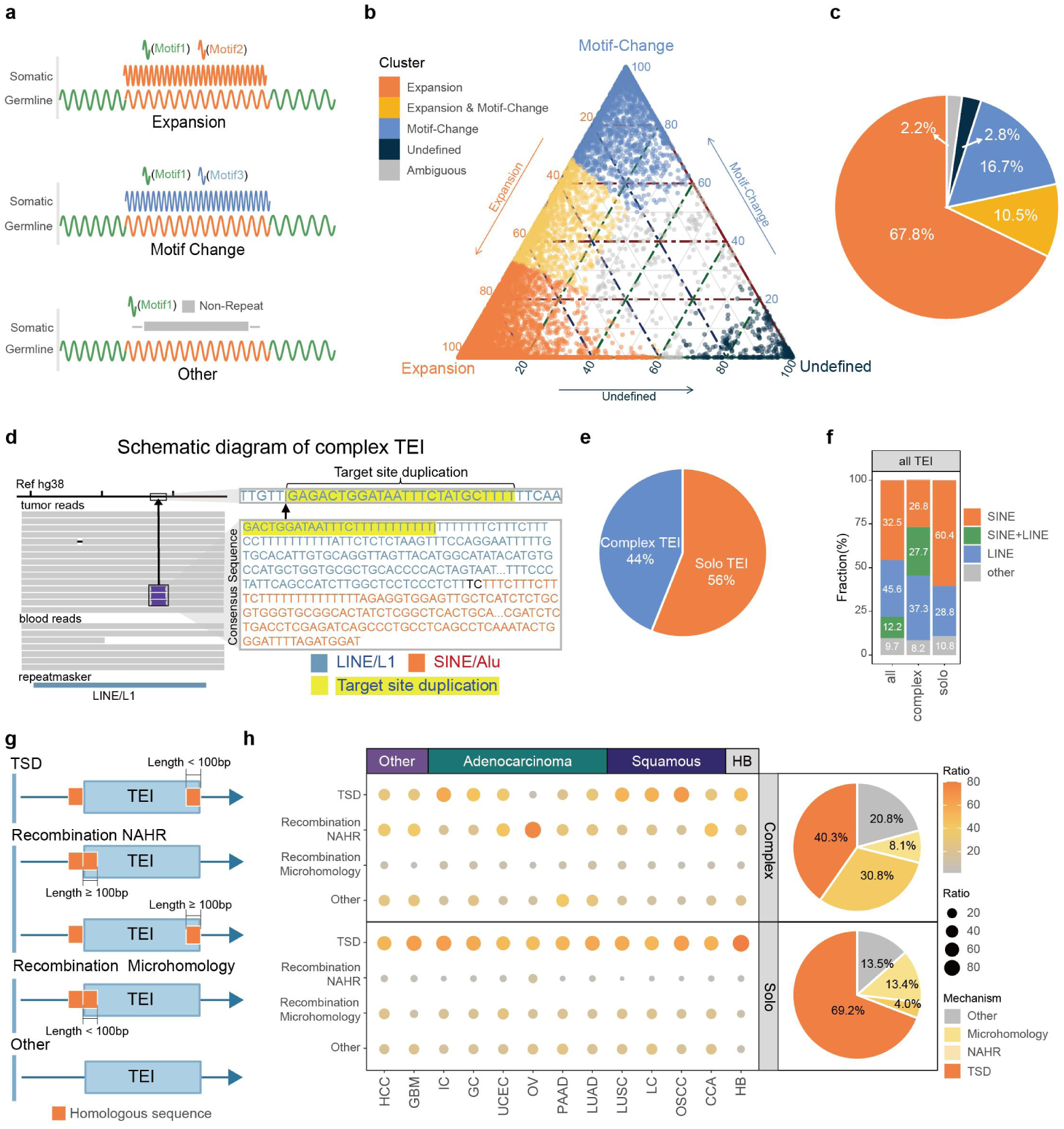
Sequence features and mechanisms of somatic TRIs and somatic TEIs. a. Schematic representation of three somatic TRI motif mutation patterns, namely expansion, motif change and other, arranged from top to bottom. In each subpanel, the first row illustrates the somatic repeat patterns, whereas the second row presents the corresponding germline repeat patterns. Different repeat motifs are color-coded for clarity. b. Ternary phase diagram illustrating the relative proportions of expansion, motif-change, and other events in somatic TRIs. Categories include: Expansion (orange), Expansion & Motif-Change (yellow), Motif-Change (blue), Undefined (dark blue), and Ambiguous (gray). c. Proportion of the five categories of somatic TRI: Expansion, Expansion & Motif-Change, Motif-Change, Undefined, and Ambiguous. d. Schematic representation of a somatic complex TEI sequence. The color coding of the bases indicates their origin, with yellow-highlighted sequences representing target site duplication (TSD) sequences. For LINE/L1 sequences, 5755 base pairs were omitted, and for SINE/Alu sequences, 141 base pairs were omitted. e. Pie charts shows the proportion of somatic complex (blue) and solo (orange) TEI. f. Proportion of different TE families within all somatic TEIs. ‘SINE’ indicates TEIs containing SINE elements but lacking LINE elements; ‘LINE’ indicates TEIs containing LINE elements but lacking SINE elements; ‘SINE+LINE’ denotes TEIs containing both SINE and LINE elements. g. Schematic illustration of somatic TEI insertion mechanisms. h. Summary of somatic TEI insertion mechanisms. The bubble chart shows the relative proportions of insertion mechanisms for complex and solo TEIs across various cancer types, with bubble size and color intensity corresponding to the magnitude of these proportions. A pie chart summarizes the overall distribution of insertion mechanisms for complex and solo somatic TEIs. TSD: target site duplication; NAHR: non-allelic homologous recombination; microhomology: microhomology recombination.

As expected, 95% of somatic TRI events were primarily characterized by the expansion of repeat motifs (Figure 2b, 2c and S5b). Among these, 67.8% involved the expansion of pre-existing germline motifs, with the proportion of mutational sequences ranging from 40.28% to 100% (average 87.89%, median 91.48%). 16.7% were driven by the expansion of newly emerged motifs, with the proportion of mutational sequences ranging from 50.15% to 100% (average 81.90%, median 82.12%). Furthermore, 10.5% of somatic TRI events exhibited a combination of both germline motif expansion (average 47.85%, median 49.03%) and the expansion of new motifs (average 45.96%, median 44.98%). In addition to these, 2.8% of somatic TRI events were classified as ‘Undefined’, where the repeat motifs in the somatic INSs could not be identified. These events had an undetermined insertion frequency ranging from 63.25% to 99.84% (average 86.64%, median 86.84%). Furthermore, 2.2% of somatic TRI events displayed a combination of all three mutational types, preventing classification into a predominant pattern. These events were categorized as “Ambiguous”. Given the challenges associated with nanopore sequencing, particularly with long tandem repeat sequences ^72^, we propose that the “Undefined” and “Ambiguous” categories may be affected by sequencing errors.

In terms of the sequence pattern of TEIs, we identified two types of somatic TEIs: ‘complex TEIs’, whose inserted sequences comprise multiple TE fragments (Figure 2d), and ‘solo TEIs’, which consist of a single TE fragment. Among the somatic TEIs, 56% were classified as solo TEIs, while 44% were complex TEIs (Figure 2e). Complex TEIs were significantly longer than solo TEIs (average length: 1,823.8 bp vs. 570.5 bp, Student’s t-test, p-value < 2.2e-16) (Figure S5c). Approximately 80% of complex TEIs exceeded 300 bp, beyond the typical read length of short-read sequencing (SRS), making them difficult to resolve with SRS. Notably, 3.4% of complex TEIs matched the known somatic recombination pattern of Alu and L1 ^33,59^. The remaining complex TEIs exhibited more intricate compositions of TE families and inverted TE segments (Table S5) — observations not systematically identified in SRS studies.

Somatic TEIs containing LINE elements were most prevalent (57.8%), followed by those containing SINE elements (44.7%) (Figure 2c, S5d and S5e). The distribution of somatic TEIs varied across different tumor types. In SCC, gastrointestinal cancers (including IC and GC), and UCEC, somatic TEIs predominantly contained LINE segments (45.4%-76.0%), consistent with previous SRS studies showing a higher frequency of somatic solo L1 insertions in SCC ^73,74^. These tumors exhibited a typical ∼300 bp somatic TEI profile with no distinct peaks (Figure S5f). In contrast, other tumors a higher proportion of somatic SINE-containing somatic TEIs (54.6%-70.4%). Peaks were identified around 300 bp and 6 kb (Figure S5f), corresponding to SINE- and LINE-containing somatic TEIs, respectively. High-burden samples exhibited a noticeable peak at approximately 300 bp, with 48.2% (5422/11238) being solo SINEs. These findings suggested differential TE activity across cancer types. Notably, younger TE subfamilies, such as AluY and L1HS, significantly contributed to the tumor genome (Figure S5g), highlighting their activity in tumor development and their key role in shaping the somatic TEI composition of the cancer genome.

To explore the insertion mechanisms, we analyzed the sequences features of somatic TEIs (Supplementary Note). By aligning somatic TEI sequences with the flanking regions of insertion sites (STAR Methods), we identified 82.1% of somatic TEIs contained homologous sequences. Based on the positions and lengths of homologous sequences, we classified somatic TEIs into two mechanisms: retrotransposition and double-strand break (DSB) repair via homologous recombination^59^. 58.2% of somatic TEIs were associated with target site duplications (TSDs), a hallmark of retrotransposition (Figure 2g and Figure S6a). The median TSD length was 14.6 bp (Figure S6h), consistent with previous findings ^74^. 31.3% of somatic TEIs exhibited homologous sequences (average: 760.8 bp) on the one side of the breakpoint, suggesting the homologous recombination. Among these, 12.3% had homologous sequences longer than 100 bp (average: 262 bp, ranging from 101 bp to 19,659 bp), similar to the non-allelic homologous recombination (NAHR) mechanism described previously (^59,75^ (Figure 2g and Figure S6b). 12.9% of somatic TEIs carried shorter homologous sequences (<100 bp, average: 7 bp), indicating microhomology-mediated recombination (Figure 2g and Figure S6c). The remaining somatic TEIs were classified as ‘other’ indicating unknown mechanisms (Figure 2g and Figure S6d).

We further uncovered the mechanisms underlying the formation of solo and complex TEIs. Somatic solo TEIs were predominantly generated via retrotransposition (69.2%), with only 17.4% arising from recombination (Figure 2e). In contrast, complex TEIs exhibited a more balanced distribution, with 38.9% arising from recombination and 40.3% by retrotransposition (Figure 2e). Notably, the mechanisms of somatic complex TEIs varied across cancer types. GC, GBM, and HCC had a larger proportion of somatic complex TEIs generated by NAHR recombination (Chi-square test, p-value < 2.2e-16, Fold Change = 2.56). Digestive system cancers (GC, IC, PAAD, and HCC) displayed a larger proportion of somatic complex TEIs with unidentified mechanisms (Chi-square test, p-value < 2.2e-16, Fold Change = 1.96). These findings suggest that the formation of somatic complex TEIs is characterized by significant stochastic variability rather than a strictly ordered process.

### Section 4: High-Frequency Somatic TRI Events among samples are potential Pan-Cancer Biomarkers

To identify non-random somatic TRI events, we compared the observed frequency distribution of somatic TRI events with the expected distribution under random conditions. This analysis revealed that 152 somatic TRI events, each detected in ≥10 patients, are exceptionally rare under the random distribution model (permutation test, p-value ≤ 0.002; STAR Methods, Figure S7a). Notably, three of these events have also been reported as recurrent repeat expansion events (rREs) with subtype-specific characteristics in previous pan-cancer studies of somatic simple repeat expansions^34^ (Table S4). We therefore defined these somatic TRI events as high-frequency somatic TRI events. Similar to the genome-wide distribution pattern of somatic TRI events, we identified that high-frequency TRI events are significantly underrepresented in coding sequence (CDS) regions compared to the genome-wide background (binomial test, FDR = 0.0425; Figure S7b). This finding suggested that the high-frequency somatic TRI events are more likely to exert their downstream effects through alterations in non-coding regions, potentially impacting regulatory processes. Previous studies have demonstrated that tandem repeat (TR) sequence variations contribute to genomic regulation primarily by modulating transcription factor (TF) binding sites ^76^ or by introducing CpG dinucleotide sequences that influence methylation patterns, thereby affecting downstream gene transcription ^77^.

High-frequency somatic TRI events are significantly enriched in 10 out of 490 ENCODE TF binding regions (Table S9) compared to low-frequency TRI events (Chi-square test, p-value ≤ 0.05, absolute ratio difference > 5%; Figure S7c). Among these, the most notable enrichment was observed in CTCF binding sites, where high-frequency somatic TRIs displayed an odds ratio of 2.032 (p-value = 3.3e-4). This suggested that high-frequency somatic TRIs may be involved in 3D chromatin structure alterations and distal regulatory processes ^78^. Additionally, when comparing somatic sequences of high-frequency TRIs to their germline counterparts, we identified significant differences in CpG dinucleotide counts in 62 out of 152 high-frequency somatic TRIs (Wilcoxon rank test, FDR < 0.05, |log2(Fold Change)|>1, Figure S7d, Table S10). Leveraging the unique ability of nanopore sequencing to simultaneously detect sequence variations and methylation states ^79^, we found that 18 out of these 62 high-frequency somatic TRIs introduced methylation marks at somatic extended CpG dinucleotides (Wilcoxon rank test, p-value ≤ 0.05, Figure S7e, Table S11). For instance, in the sample Lung120, we investigated that a somatic TRI region (chr6:64,295,338–64,295,575) within an intron of the *EYS* gene introduced 55 methylated CpG sites (Figures S7f-S7i, Supplementary Note). As mentioned in the previous study ^77^, these somatic TRIs may act as potential regulatory elements.

To investigate the association between high-frequency somatic TRIs and cancer-related biological processes, we performed functional enrichment analysis on genes neighboring high-frequency somatic TRIs. Compared to low-frequency somatic TRIs, genes associated with high-frequency somatic TRIs were significantly enriched in functional categories related to focal adhesion, apical junctions, and cytokine-cytokine receptor interactions (Figure 3a, Chi-square test, p-value < 2e-4). Among these, the somatic TRI event located at chrX:9,791,969–9,792,160, within the intron of *SHROOM2,* which is associated with apical junction functions, exhibited the highest shared frequency across patients (57 out of 299, Figure 3b, Table S12). Using targeted PCR and Sanger sequencing, we validated this somatic TRI event in 25 paired tumor and blood samples (Figure 3c, Figure S8a, Table S13 and Table S14). Additionally, this *SHROOM2*-related TRI was detected in liver cancer and intestinal cancer cell lines (Figure S8b). Overall, this somatic TRI event was present in nearly 20% of pan-cancer samples (57 out of 299) and occurred across 11 of the 12 cancer types analyzed. Notably, this somatic TRI exhibited particularly high recurrence rates in HCC (58.82%; 10 out of 17) and IC (40%; 6 out of 15). Intriguingly, all multi-site biopsies from three HCC patients with this somatic TRI consistently contained it, while biopsies from two HCC patients without this somatic TRI consistently lacked it. This result indicated a high degree of intra-tumor consistency for this somatic TRI. Based on length distribution, somatic TRI sequences within this window were categorized into two groups: long-fragment expansions and short-fragment expansions (Figure 3d). The long-fragment expansions ranged from 3,896 to 8,284 bp, whereas the short-fragment expansions ranged from 639 to 3,259 bp (Figure 3e). Interestingly, the two cancer types with high recurrence rates (HCC and IC) predominantly carried somatic TRIs with long-fragment expansions (HCC: 7 out of 10; IC: 5 out of 6; Figure 3f, Fisher’s exact test, p-value = 3.29e-6). These long-fragment expansions were primarily characterized by the amplification of the AGAAT repeat motif (Figure S9, Table S15).

**Figure 3.**
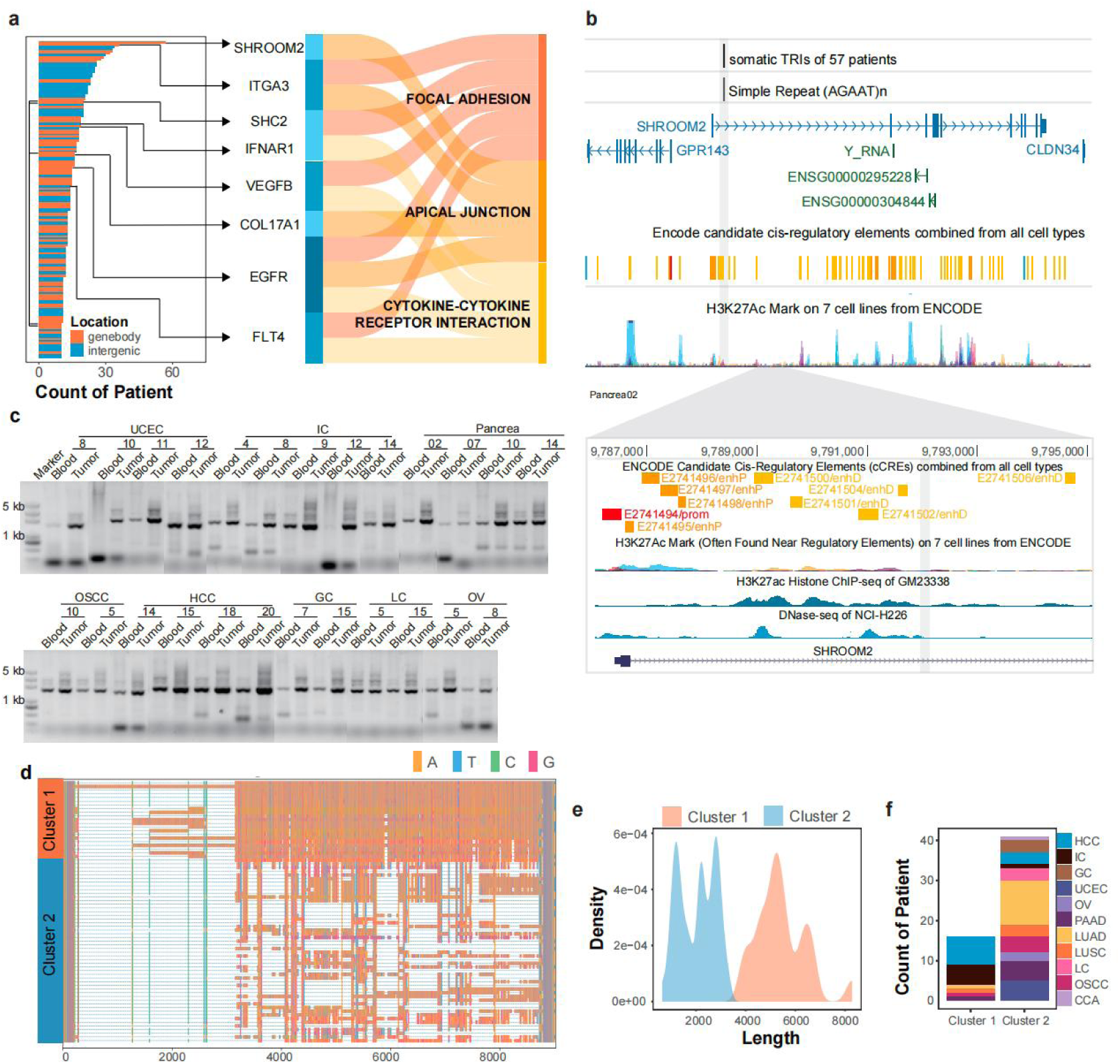
High-frequency somatic TRIs located in the intron of *SHROOM2*. a. Bar plot showing the number of patient<u>s</u> harbored somatic TRIs in each high-frequency TRI window (>= 10 patients), along with the pathways enrichment analysis of genes located within or near these high-frequency somatic TRI windows. b. Genome browser tracks displaying the ENCODE-annotated genes, and ENCODE-predicted cis-regulatory elements (cCREs) surrounding somatic TRIs in the SHROOM2 gene locus. Lower panel: Epigenomic signals around somatic TRIs in SHROOM2, as visualized by chromatin marks. c. Detecting somatic TRI events harbored by *SHROOM2* through PCR in clinical samples, blood DNA as a control. d. The detailed sequence overview of somatic TRIs harbored by *SHROOM2* in different clusters. The ATCG bases are colored differently, with gaps represented as short dashes. e. Distribution of somatic TRI lengths in *SHROOM2* across different clusters. f. Bar chart showing the distribution of cancer types in patients harboring somatic TRIs within SHROOM2 across the different clusters.

Subsequently, using our NSCLC NGS RNA-seq dataset, we performed differential gene expression analysis within a 300 kb window surrounding all high-frequency somatic TRI events detected in NSCLC. This analysis identified 51 differentially expressed genes associated with 33 high-frequency somatic TRI events (Figure S10a, Table S16). Among these, we identified the somatic TRI event located at chr7:121,603,262–121,603,506, which was previously reported in our study to be present in nearly 40% of LUSC samples (Figure S10b and S9c). This somatic TRI was shown to regulate the expression of its downstream *PTPRZ1* gene by altering the 3D genomic structure ^25^. Additionally, we identified two other somatic TRI events associated with the expression of genes in the PTP family: the somatic TRI at chr1:202,204,213–202,204,518, which was correlated with the upregulation of *PTPN7* expression (Figure S10b and S9d), and the somatic TRI at chr7:158,595,097–158,595,342, which was associated with the downregulation of *PTPRN2* expression (Figure S10b and S9e). Notably, while the somatic TRI at chr7:121,603,262–121,603,506 was significantly enriched in LUSC and LC samples, the other two PTP-related somatic TRI events did not exhibit enrichment in specific cancer types (Figure S10f). Furthermore, we noticed the presence of these three TRI events across the pan-cancer cohort, encompassing 66 samples.

By performing clustering analysis based on the distribution spectrum of somatic TRI events, we identified that samples from HCC, GBM, GC, and LUAD formed four distinct clusters (Figure 4a). Notably, GC, GBM, and LUAD samples did not harbor somatic TRI events with an intra-cancer frequency exceeding 30% (Figure 4b), indicating a high degree of inter-patient heterogeneity for TRI events within these cancer types. In total, we identified 12 somatic TRI events that were detected within single cancer types with high frequency (Figure 4c, STAR Methods). HCC harbored the highest number of intra-cancer high-frequency somatic TRI events (4 out of 12; Table S16, Figures 4d). These included a previously described somatic TRI event located within the SHROOM2 intron, as well as three additional events: one within the PALMD intron, predominantly characterized by expansions of the AAAG motif (chr1:99,683,035–99,683,173, Figure S11a), which is consistent with previous reports in other pan-cancer somatic tandem repeat study ^34^ (Table S4); another within the *LIPG* intron, mainly involving expansions of the GGGAGA motif (chr18:49,579,001–49,579,348, Figure S11b); and an intergenic event featuring expansions of the GGTTT motif (chr1:34,802,227–34,802,447, Figure S11c). Notably, these HCC-specific high-frequency somatic TRI events were also recurrently detected in IC, UCEC, OV, OSCC, and LC. Genes such as *SHROOM2* and *PALMD* are implicated in the regulation of the cytoskeleton ^80–83^, while *LIPG* is involved in lipid metabolism ^84^. High-frequency somatic TRIs shared across different cancer types may suggest common malignant biological behaviors, making them potential diagnostic and therapeutic biomarkers for these cancers.

**Figure 4.**
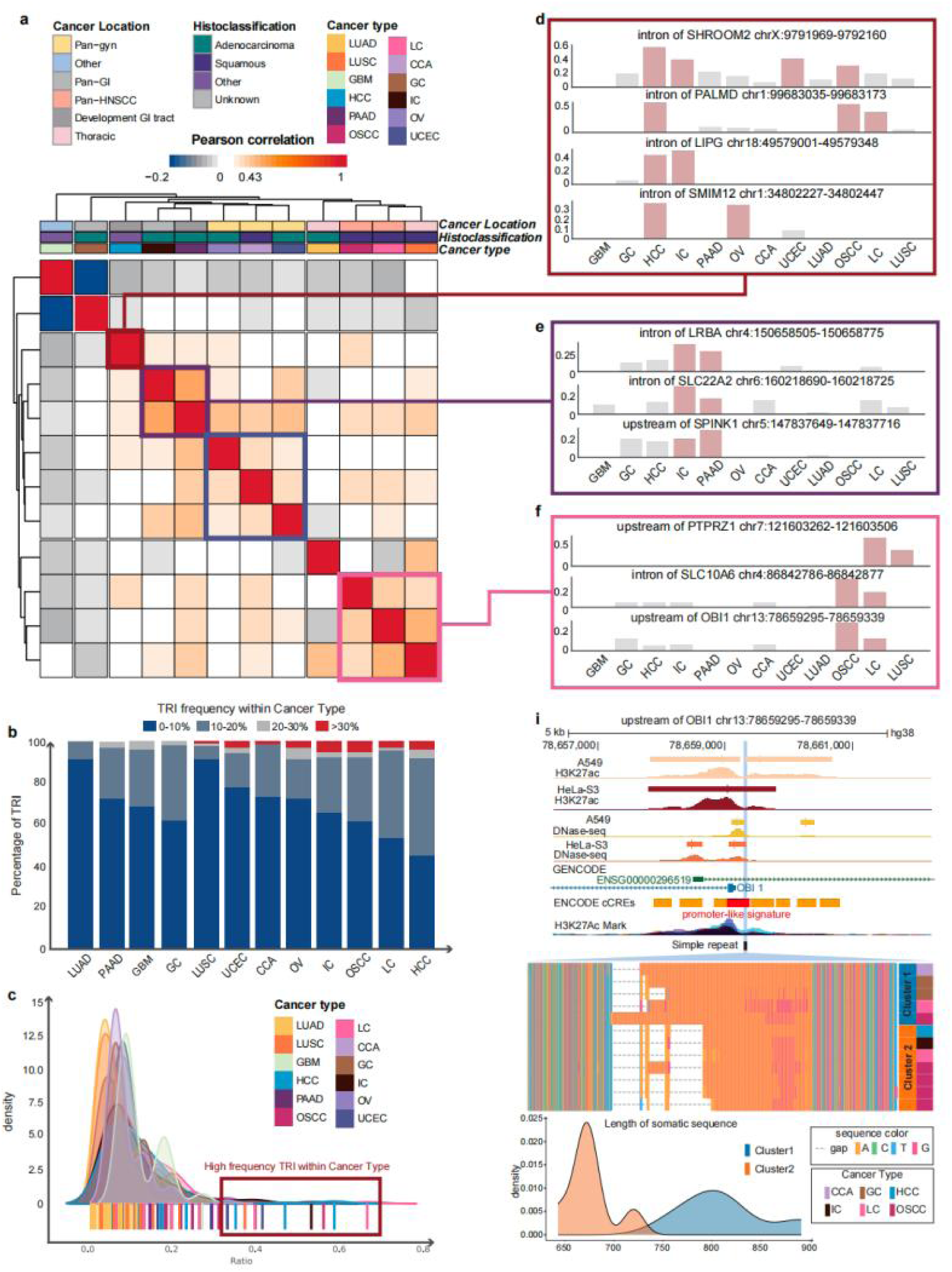
The characteristic of somatic TRI clusters. a. Clustering heatmap of somatic TRI frequencies across cancer types. b. Proportion of tumor samples with somatic TRIs. The number represents the proportion of mutations within the cancer type. Different colors represent various percentage categories. c. Frequency distribution of somatic TRIs across cancer types. The bars at the bottom represent the mutation frequency of the mutation within individual cancer types, with colors indicating different cancer types. d. Bar plots showing the frequency of HCC associated high-frequency somatic TRI across cancer types. The high frequency cancer types (over 30%) are filled in red. e-f. Bar plots showing the frequency distribution of high-frequency somatic TRI in IC&PAAD (e), as well as OSCC, LC and LUSC (f) clusters compared to other cancer types. Bars of significant cancer types are filled in red. g. Browser plot displaying the epigenomic signal, ENCODE-annotated gene transcript, and cCREs (Candidate Cis-Regulatory Elements) predicted by ENCODE around the upstream somatic TRIs of the OBI1 gene. The detailed sequence overview of different clusters surrounding the upstream somatic TRIs of OBI1 is presented in the lower enlarged panel, with the length distribution of these 2 clusters color-coded.

Additionally, digestive system adenocarcinomas (IC and PAAD), gynecological tumors (CCA, OV, and UCEC), and three SCCs (OSCC, LC, and LUSC) formed three distinct clusters based on the frequency of somatic TRI events (Figure 4a). Each cluster exhibited specific somatic TRI events. Within the IC-PAAD group, we identified three somatic TRI events with significantly higher frequencies compared to other cancer types (Student’s t-test, p-value < 0.05): a somatic TRI event located in the *LRBA* intron (chr4:150,658,505–150,658,775; Figure S11d), another in the *SLC22A2* intron (chr6:160,218,690–160,218,725; Figure S11e), and the third upstream of the *SPINK1* gene (chr5:147,837,649–147,837,716, 6073 bp upstream; Figure S12a). These genes are associated with tumor cell growth (*LRBA*) ^85^, chemotherapeutic resistance (*SLC22A2*) ^86^, and tumor dedifferentiation (*SPINK1*) ^87^, respectively. In contrast to the IC-PAAD group, gynecological tumors (OV, CCA, and UCEC) formed a distinct cluster in the somatic TRI frequency spectrum, but exhibited low recurrence rates for their specific somatic TRI events. Two cluster-specific somatic TRI events (Student’s t-test, p-value < 0.05) included one upstream of the ENSG00000257323 gene (chr12:74,615,532–74,615,609; Figure S12b) and another within the *MED16* intron (chr19:878,845–879,150; Figure S12c). These events appeared at frequencies below 30% within any individual gynecological tumor type. The SCC cluster, consisting of OSCC, LC, and LUSC, exhibited three cluster-specific high-frequency somatic TRI events (Figure 4f). These included the previously described *PTPRZ1*-related somatic TRI event (Figure S10c), as well as two somatic TRI events located in the promoter of the OBI1 gene (chr13:78,659,295–78,659,339; Figure 4g) and the *SLC10A6* intron (chr4:86,842,786–86,842,877; Figure S12d). In the available samples, we validated the cluster-specific high-frequency TRI events through PCR experiments and Sanger sequencing, highlighting the existence of similar somatic TRI distribution patterns among cancer types within the same cluster (Figure S12, Table S14).

### Section 5: somatic TEIs were enriched in the MHC II gene cluster

To investigate the somatic TEI distribution across cancer types, we clustered samples based on somatic TEI breakpoints, identifying five distinct clusters (Table S1) with unique histological compositions (Figure 5a). Cluster 1 predominantly comprised SCC samples (67%) (Figure 5b and 5c), including 87.5% of LUSC and 68.8% of OSCC cases (Figure 5c and S14a). Notably, approximately 20% of ADC samples were also assigned to Cluster1, with IC (73.3%) and OV (50%) being the most represented (Figure 5c and S14a). This suggested that IC and OV share TEI characteristics with SCC samples in Cluster 1. Moreover, ADC samples in Cluster 1 exhibited lower somatic TEIs burden (mean: 36.7, median: 19) compared to those in other clusters (mean: 156, median: 23) (Welch two-sample test, p = 0.005), indicating the distinct somatic TEI profiles of ADC samples within Cluster 1. 57.7% of Cluster 1 SCC samples further subdivided into two subclusters. In Subcluster 1, 75% of the samples were SCC, while in Subcluster 2, nearly 60% of the samples were SCC (Figure 5d and S14b). Subcluster 1 exhibited a significantly higher somatic TEI burden than Subcluster 2 (Figure S14c), highlighting the substantial molecular heterogeneity within SCC samples.

**Figure 5.**
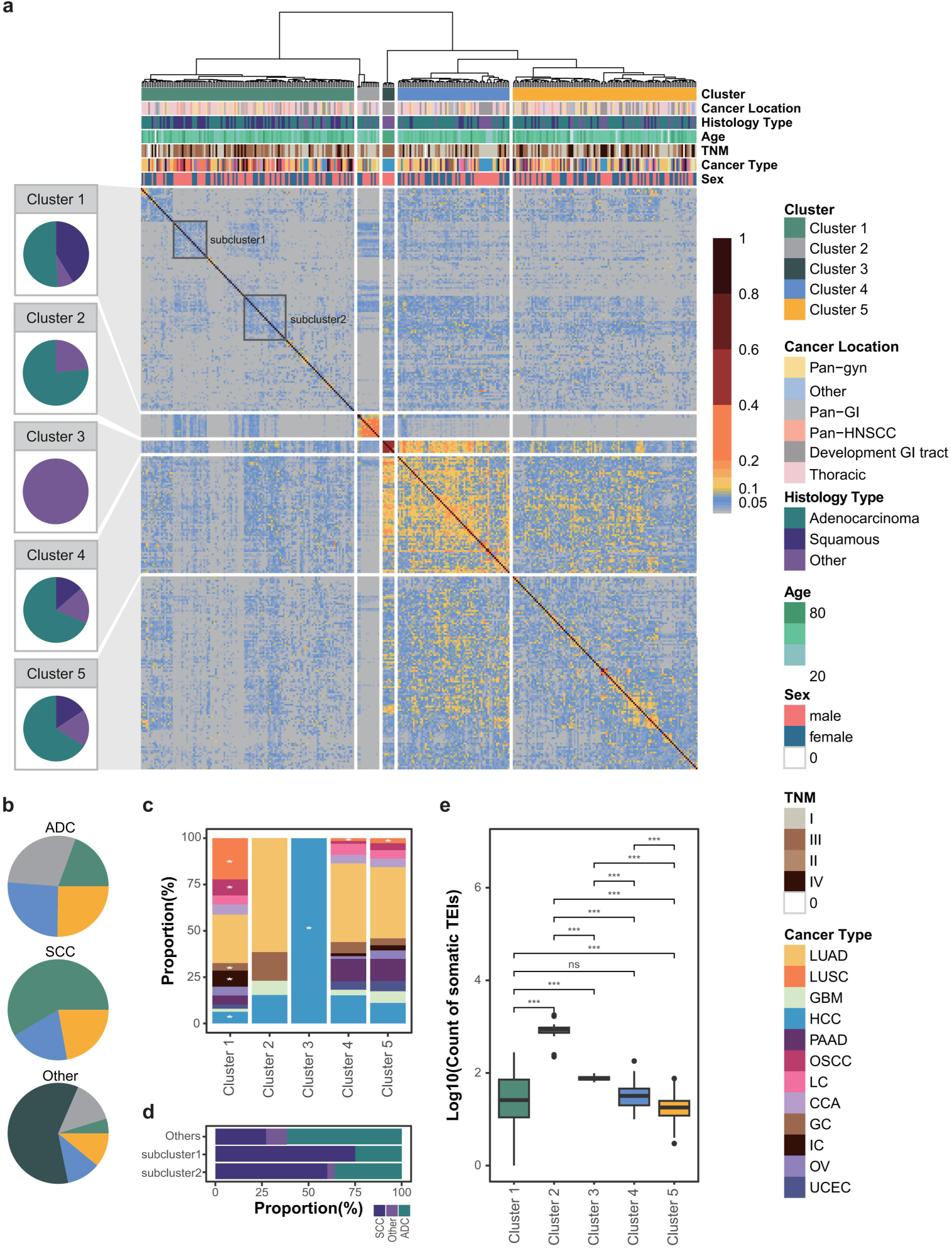
Clusters of somatic TEIs. a. Heatmap showing the clustering results of somatic TEIs. Each column and each row represent Jaccard similarities of a sample compared to others. Hierarchical clustering was performed according to Jaccard similarity with Ward.D linkage. Each cluster has a pie chart to indicate the proportions of adenocarcinoma, squamous cell carcinoma, and others. Two SCC-subclusters of cluster 1 are distinguished. b. Pies showing the proportional of 5 clusters within the three major histology categories: SCC (squamous cell carcinoma), ADC (adenocarcinoma), and Others. c. A stacked bar chart displays the distribution percentages of different cancer types across five clusters. d. Proportional distribution of ADC (adenocarcinoma), SCC (squamous cell carcinoma), and others within subclusters of Cluster 1. e. A boxplot depicting the number of somatic TEIs (log10) within 5 clusters, with the p-value calculated using t-test. ns p>0.05, *p < 0.05, **p < 0.005, ***p < 0.0005

Cluster 2 encompassed tumor samples with the highest levels of TEI burden, including all high-burden samples and two multi-site biopsies from a LUAD patient, which ranked just below the high-burden samples. Although multi-site biopsies were collected from 9 patients, with samples from the same patient clustering together, only the complete set from patient HCC10 was assigned to Cluster 3, suggesting a potential association between he multi-lesional nature of the patient and their distinct somatic TEI profile.

Clusters 4 and 5 were predominantly composed of ADC samples (68% and 66%, respectively) but differed in somatic TEI burden and distribution. Cluster 4 exhibited a higher somatic TEI burden and a greater degree of overlap between samples (Figure 5e), indicating a more homogeneous TEI profile. This cluster also included poorly differentiated, non-metastatic CCA samples and early-stage (I/II) HCC biopsies, which clustered with multi-site samples from a stage I LUAD patient. In contrast, cancers commonly associated with high heterogeneity, such as PAAD, UCEC, and GBM, were largely detected in Cluster 5, with PAAD samples showing duodenal or peripancreatic invasion (Figure S14a, Table S1).

We analyzed high- and low-frequency somatic TEI regions across the clusters, covering 1,464 1Mb genomic windows. In Cluster 1 and Cluster 5, we identified 15 and 6 regions, respectively, with significantly low-frequency somatic TEIs (Fisher’s exact test, FDR < 0.01, Fold change < 0.5, and >10% of non-cluster samples showing somatic TEIs in the same window). In contrast, Cluster 2, Cluster 3, and Cluster 4 contained 1384, 45, and 14 regions, respectively, with significantly high-frequency somatic TEIs (Fisher’s exact test, FDR < 0.01, Fold change > 2, and >10% of cluster samples exhibiting somatic TEIs in the same window) (Figure S14d, Table S17).

To identify non-randomly enriched TEI windows, we applied a permutation test (STAR Methods). In windows with ≥14 patients, somatic TEI frequency was significantly higher than in randomly selected background regions. This suggested that these windows are non-randomly enriched with somatic TEIs (permutation test, p-value=0.0002, Figure 6a). Given the high sequence variability and repetitive nature of centromeric and pericentromeric regions, as well as the limited functional data and reference genome quality in these areas, we excluded them from further analysis. After exclusion, the chr6:32,000,000-33,000,000 window, containing the MHC II gene cluster, exhibited the highest somatic TEI count and the greatest proportion of TEI-positive samples (40.6%) (Figure 6b). Particularly in Cluster 1 (67% SCC samples), the proportion of samples harboring TEI (27.8%) is significantly lower than that in clusters predominantly composed of non-SCC samples (49.7%) (Fisher’s exact test, FDR=0.0007).

**Figure 6.**
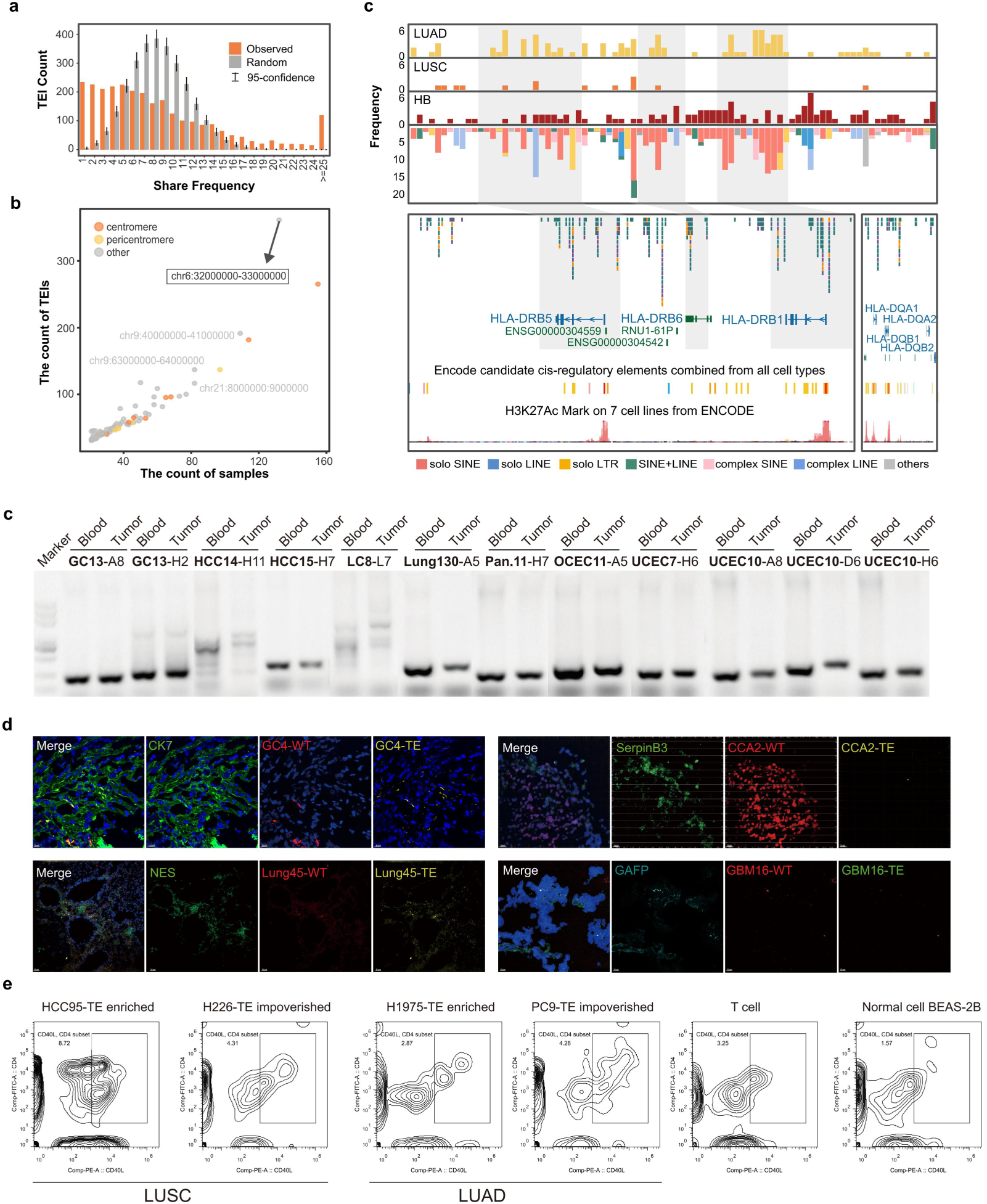
The MHC II gene family harbored high frequency somatic TRIs. a. Distribution of the number of shared TEI samples. Gray: random distribution (Methods); Orange: observed distribution. The x-axis represents the number of samples sharing a somatic TEI, and the y-axis represents the count of somatic TEIs. b. Scatter plot illustrating the number of samples sharing somatic TEIs and the count of somatic TEIs. Each dot represents a 1Mb genomic window, with the color indicating the source of its chromosomal coordinates. The genomic window containing the highest number of TEIs, chr6:32,000,000-33,000,000, is highlighted. c. Bar plots showing the number of somatic transposable element insertions (TEIs) within 500 bp windows in LUAD (first track), LUSC (second track), and high-burden samples (third track). The fourth track displays the cumulative number of somatic TEIs within 500 bp windows, with bars color-coded according to the type of TEI. Each bar represents a 500 bp window. The browser plot includes epigenomic signals, ENCODE-annotated gene transcripts, and candidate cis-regulatory elements (cCREs) predicted by ENCODE around the somatic TEIs within the MHC II gene family. d. Detecting somatic TEIs events through PCR in some clinical samples, blood DNA as a control (A: ALL; H: HCC; L: LC; D: DRB5). e. Detecting somatic TEIs events through FISH in in some sample tissue slices, CK7/serpinB3/NES/GAFP is the marker for GC/CCA/LUNG/GBM respectively. WT refers to the sequence without somatic TEIs events, TE contains the sequence of somatic TEIs events. Flow cytometric analyses of CD40L mobilization in CD4+ T cells after 24 hours of in vitro co-culture with tumor cells.

Somatic TEIs in this region were primarily located in the intergenic and coding regions of the *HLA-DRB1*, *HLA-DRB5*, and *HLA-DRB6* genes (Figure 6c). The distribution of somatic TEIs varied across cancer types. High-burden samples showed widespread somatic TEIs around the MHC class II gene cluster, while LUAD samples showed higher frequencies of somatic TEIs than LUSC (Figure 6c). Notably, in the *HLA-DRB1* genebody and its 5 kb flanking regions, somatic TEIs were absent in LUSC, but frequent in LUAD (Fisher’s exact test, FDR = 0.00066). Most somatic TEIs in this region were solo SINEs (Figure S15a), consistent with previous findings of LINE enrichment in SCC samples.

Next, we examined the impact of somatic TEIs in the chr6:32,000,000-33,000,000 region on gene expression using transcriptomic data from LUAD tumor samples. Comparison of 32 LUAD samples with somatic TEIs in this region to 39 samples without revealed 1,798 differentially expressed genes (p value < 0.05, |log2(FC)| > 1, Table S18, Figure S15b). These genes were enriched in cytokine signaling and immune response pathways (Figure S16a and S16b). 6 immunoglobulin-related and 4 T-cell receptor-related genes were significantly downregulated in LUAD samples with somatic TEIs (Figure S15c). These results suggested that TEIs in this region may contribute to immune evasion and tumor progression.

Next, we further validated our bioinformatics findings through experiments. First, in a subset of samples that underwent LRS, the authenticity of the TEI event was gauged by Polymerase Chain Reaction (PCR). With blood DNA serving as a reference, when a TEI event was present in the tumor tissue sample, the target band obtained from the tumor tissue sample exceeded that of the blood tissue under identical primer amplification conditions (Figure S15 d and e, Table S19). Moreover, through the utilization of specific fluorescence in situ hybridization (FISH) probes (Table S20), TEI events were also discerned in certain sample tissue sections (Figure 6e). It is widely recognized that tumor tissues encompass a rich and diverse spectrum of cell types, inclusive of infiltrating immune cells. Considering that the TEI insertion site is located in the MHC II gene cluster region, we gathered additional fresh clinical samples (Table S21). By sorting CD45 positive cells, we further amplified the TEI events identified in this region within the clinical sample LRS sequencing outcomes. The findings confirmed that these insertions occurred in both CD45 positive and negative cells (considered as tumor cells) (Figures S17). Subsequently, we detected TEI events within the MHC II gene cluster in 33 tumor cell lines stored in the laboratory. In alignment with the sequencing results in clinical samples, non-squamous carcinomas harbored a greater number of TEI events in contrast to squamous carcinomas (a total of 63 TEIs were detected, with an average detection of 11.07 in squamous carcinomas and 19.35 in non-squamous carcinomas) (Figures S18, S19, and S20a).

MHC class II molecules play a crucial role in cellular immunity. In antigen-presenting cells (such as macrophages, dendritic cells, and B cells), MHC class II molecules are involved in the processing and presentation of ingested exogenous antigens into peptide fragments, and presenting them to CD4+ T cells, thereby initiating the adaptive immune response^88^. Here, we compared the in vitro phenotypes of CD40L activation by TEI-enrich and TEI-impoverished tumor cells through in vitro co-culture of tumor cells and mouse CD4-positive t cells. The results showed that in squamous cell carcinoma cell lines, the surface expression of CD40L on TEI-rich tumor cells was twice that of TEI-impoverished cells (Figure 6e and Figure S20b); while the activation situation is reversed in adenocarcinoma cells (Figure 6e and Figure S20b), this all suggested that the frequency of TEI events and the sequence type of TEI event insertions may be important factors in regulating the success of tumor adaptive immune initiation.

## Discussion

This study was designed to elucidate high-frequency somatic large-scale REIs (>50bp) in the pan-cancer genome. Previous research has been constrained by the limitations of SRS technology, which hindered the precise identification of these somatic SVs and their distribution across tumor population. LRS technology offers a promising solution to these challenges ^89^. In this work, we applied nanopore LRS to sequence 325 tumor samples and their matched blood samples from 12 cancer types, generating nearly 46 Tb of data. This extensive LRS dataset, one of the largest to date, provides a robust resource for exploring somatic large-scale REI events across the pan-cancer landscape.

Current LRS-based somatic SV detection algorithms are hindered by alignment biases, particularly in repetitive genomic regions. These biases, stemming from the seed-and-chain alignment strategy, undermine the accuracy of insertion detection ^89,90^. Although assembly-based approaches hold promise, they depend on high-quality individual genome assemblies, which are challenging to produce in the context of heterogeneous tumor genomes ^91,92^. To overcome these limitations in detecting somatic REIs, we developed specialized bioinformatics methods tailored for somatic TRI and TEI detection, respectively. By harnessing the technical strengths of long reads that span somatic insertions, and combining unsupervised learning with partial order alignment graphs and statistical analysis of local sequence features, we minimized errors introduced by reference genomes and alignment algorithms, thus enabling more accurate detection of somatic REIs. Recent advancements in long-read sequencing (LRS) technology, alongside continuous improvements to the human genome reference—particularly the completion of the Telomere-to-Telomere (T2T) project ^93^—have significantly enhanced the study of large-scale repetitive sequence variations, such as centromeric and segmental duplications ^94^. This study also benefits from these developments, uncovering numerous related somatic REIs. However, the structural complexity and dynamic nature of these regions, marked by variation in repeat copy number and sequence divergence, present substantial challenges to accurate detection. Consequently, refined bioinformatics methods and ultra-long sequencing technologies are expected to provide critical solutions to these issues ^95^.

Using our developed methodology, we identified a significant number of high-frequency somatic REIs across pan-cancer genomes. In addition to the somatic TRI events located within the introns of the *SHROOM2* and *PALMD* genes, which exhibited remarkably high recurrence in HCC, IC, CCA, UCEC, OV, LC, OSCC, and LUSC. Interestingly, cancers from anatomically neighboring tissues tend to exhibit similar somatic TRI profiles. For example, the gynecological cancers CCA, UCEC, and OV shared a similar somatic TRI spectrum, with a somatic TRI event in the intron of *MED16* being specific to these tumor types. The *MED16* gene is associated with estrogen receptor sensitivity in the previous research ^96^, the gynecological cancers specific somatic TRI event may play a role as a regulatory element of this gene. Similarly, squamous cell carcinomas, including LUSC, LC, and OSCC, displayed similar somatic TRI profiles. Notably, a somatic TRI event identified in approximately 40% of LUSC samples was found to occur at an even higher frequency in LC samples (nearly 70%). This event has been shown to influence the three-dimensional structure of the genome, thereby modulating the expression of the downstream gene *PTPRZ1* through enhancer activity ^25^. In addition to *PTPRZ1*, further analysis of paired RNA-seq data from non-small cell lung cancer (NSCLC) revealed two additional somatic TRI events (chr1:202,204,213-202,204,518 and chr7:158,595,097-158,595,342, Figure S10a and S10b), which were significantly associated with the expression of their downstream genes, *PTPN7* and *PTPRN2*. Notably, 20.31% of pan-cancer patients carried at least one somatic TRI event potentially associated with the PTP gene family. As an important receptor kinase family, the PTP genes play a crucial role in immune evasion mechanisms ^71^. We propose that somatic TRI events may function as key distal regulatory elements for the expression of this gene family. Further investigation into the transcriptional regulatory mechanisms underlying these somatic TRI events will be essential to understanding the role of the PTP gene family in immune regulation. Understanding the impact of the high-frequency somatic TRI events we identified on the tumor genome network is a key focus for future research.

The success of immune checkpoint inhibitors in certain cancers has demonstrated that the immune system can target and reject tumors through T cell responses. Tumor cell recognition relies on MHC-mediated presentation of tumor-associated antigens. Our research reveals that somatic TEIs are enriched in MHC II gene regions, with over 40% of pan-cancer samples harboring somatic TEI events in these regions. Notably, LUAD exhibited a significantly higher somatic TEI burden in MHC II regions compared to LUSC (Figure 6c). Although the immunological effects of these somatic transposable element insertions (TEIs) have not been fully elucidated, co-culture experiments indicated that TEI-enriched adenocarcinoma cell lines are more potent in suppressing CD4+ T cell activation compared to TEI-impoverished counterparts. Furthermore, LUSC patients have been shown to derive greater clinical benefit from PD-1 inhibitors than LUAD patients ^97^. Given that TEI-enriched LUAD cell lines suppress CD40L expression on CD4+ T cells, we hypothesize that immune activation within these tumors may be constrained. Since PD-1 inhibitors function to counteract PD-L1-mediated immune suppression of CD8+ T cells, their efficacy in TEI-enriched LUAD may be diminished due to the absence of CD4+ T cell activation. Nevertheless, this hypothesis warrants further in vivo validation. Moreover, we propose that TEI enrichment in MHC II regions could serve as a potential target for tumor immune evasion. Yet, the precise mechanisms by which TEIs modulate immune responses remain poorly understood, highlighting a critical area for future investigation in tumor immunology.

Here, we leverage LRS technology to investigate somatic REIs and their potential as diagnostic and therapeutic biomarkers in cancer. By integrating optimized bioinformatics algorithms and advancing long - read sequencing technologies, future studies will be able to systematically characterize diverse somatic genomic alterations, such as translocations and inversions, from the dataset we provided. Additionally, integrating multimodal nanopore sequencing data will enable single-cell resolution analysis of large genomic regions and their associated epigenetic modifications, as demonstrated in recent studies ^98^. These efforts will provide critical insights into the regulatory mechanisms underlying large-scale somatic structural variations (SVs) in cancer genomes, shedding light on their functional impact. While our current dataset offers valuable insights into pan-cancer diagnostics, expanding the cohort to include diverse cancer types and enhancing clinical follow-up data—such as treatment responses and patient survival—will enable a comprehensive exploration of the relationship between somatic SVs and cancer outcomes. To achieve this, we are actively expanding our research cohort to encompass a broader spectrum of tumor types and patient demographics, ensuring robust characterization in future studies.

In conclusion, our study demonstrates the transformative potential of long-read sequencing in cancer genomics. By expanding the understanding of the genetic variants of cancer, this work sets the stage for future studies that will explore the role of SVs in human disease. As LRS technology and analysis methods evolve, we anticipate these advancements will enable more accurate SVs detection and a deeper insight into their functional consequences, ultimately guiding the development of targeted therapies and personalized treatment approaches in oncology.

## STAR Methods

### Patient Cohort and Ethics

The matched tumor-blood samples across 8 cancer types (HCC, GBM, PAAD, LUAD, IC, GC, LUSC, and LC) were collected from patients who undergone surgery at our hospital. The matched tumor-blood samples across 3 gynecological cancer types (CCA, OV and UCEC) were collected from patients who undergone surgery at West China Second University Hospital, Sichuan University. And the matched tumor-blood samples of OSCC were collected from patients who undergone surgery at West China Hospital of Stomatology, Sichuan University. None of the patients were received preoperative chemotherapy or radiotherapy. All diagnoses confirmed through histological review by a board-certified pathologist. Cancer clinical stage was determined based on the 8th edition of the American Joint Committee on Cancer (AJCC) TNM staging system. This study protocol was approved by the Institutional Review Board of our hospital (Ethics: Project identification code: 20242416). Written informed consent was obtained from all patients. Demographic and clinical data were collected for analysis(Table S1).

### PromethION whole genome Sequencing

High-molecular-weight (HMW) genomic DNA (gDNA) was extracted from tissue or blood samples using the MagAttract HMW DNA Kit (QIAGEN). DNA quality was assessed by Novogene Co. LTRI. using three standard methods: (1) DNA purity was determined by measuring OD260/280 and OD260/230 ratios with a NanoDrop spectrophotometer, (2) agarose gel electrophoresis was used to evaluate DNA degradation and detect RNA contamination, and (3) DNA concentration was quantified with the Qubit® DNA Assay Kit and Qubit® 3.0 Fluorometer (Life Technologies, CA, USA).

For library preparation, 273 samples (84%) were processed using the SQK-LSK109 kit, while 52 samples (16%) were prepared with the SQK-LSK114 kit (Oxford Nanopore Technologies, UK), following the manufacturer’s protocols (Table S1). For each sample, 48 μL of gDNA (2 μg) and nuclease-free water (NFW) were combined with 3.5 μL of NEBNext FFPE DNA Repair Buffer, 2 μL of NEBNext FFPE DNA Repair Mix, 3.5 μL of NEBNext Ultra II End Prep Reaction Buffer, and 3 μL of NEBNext Ultra II End Prep Enzyme Mix. The mixture was gently combined in a PCR tube, spun down, and incubated sequentially at 20 °C for 5 minutes and 65 °C for 5 minutes. The prepared sample was transferred to a 1.5-mL tube, and 60 μL of AMPure XP beads was added. After gentle mixing and two washes with 70% ethanol, the purified DNA was combined with 25 μL of Ligation Buffer (LNB), 10 μL of NEBNext Quick T4 DNA Ligase, and 5 μL of Adapter Mix (AMX). This mixture was incubated at 25 °C for 20 minutes. Subsequently, 40 μL of AMPure XP beads was added, and the pellet was resuspended in 25 μL of Elution Buffer (EB).

For sequencing preparation, 30 μL of flush tether (FLT) was added to PromethION Flush Buffer (PFB), and 500 μL of the Priming Mix was loaded onto a PromethION flow cell and equilibrated for 5 minutes at room temperature. The library preparation involved mixing 75 μL of sequencing buffer (SQB), 51 μL of loading buffer (LB), and 24 μL of DNA library. This was followed by loading 500 μL of the Priming Mix and 150 μL of the library onto the flow cells. DNA sequencing was performed on flow cells as per the manufacturer’s instructions. Samples processed with SQK-LSK109 were sequenced on R9 flow cells (84%), while those prepared with SQK-LSK114 were sequenced on R10 flow cells (16%). Base calling was performed using Guppy 3.2.8 with default parameters in real time during sequencing.

### Short-read RNA-seq Data Analysis

For some NSCLC samples, paired-end reads were aligned to the hg38 reference genome using HISAT2 ^99^ (version 2.1.0). SAMtools ^100^ was used to sort and index the bam files. StringTie ^101^(version 1.3.6) was used to generate the GTF files containing the read count information and calculate the TPM matrix of gene expression for all genes across all samples. The raw count matrix of gene expression for all genes across all samples is calculated by htseq-count ^102^ (version 2.0.2) based on the gencode annotation file version 42.

### Detection of differential expression gene

In order to defined potential gene associated to somatic REIs, we classified NSCLC samples into two groups based on whether harboring target somatic REI event. Then the gene expression normalized and compared between two groups through DESeq2 ^103^. Genes with p-value <= 0.05 and |log2fold-change| > 1 were defined as differential expression genes.

### Oxford Nanopore Sequencing Data Processing

Initially, raw nanopore sequencing data were processed using NanoFilt (v2.8.0, https://github.com/wdecoster/nanofilt.git, parameters: -q 7 -l 1000) to filter the fastq files and generate clean reads. The quality of these filtered reads was then assessed using NanoPlot ^104^(v1.39.0, parameter: --maxlength 40000). Next, the ONT long reads were aligned to the hg38 reference genome using Minimap2 ^44^ (v2.2.17) with the following parameters: --MD, -ax map-ont, -L, and -t 30.

### Tumor purity calculation

Somatic variants (SNVs) and copy number alterations (CNVs) were identified in tumor samples using multiple approaches. For LRS WGS sequencing data, SNVs were called using ClairS ^105^ (version 0.1.2) with parameters set to --platform ont_r10 --min_coverage 3. CNVs were detected using Control-FREEC ^106^ (version 11.6) with default settings. For the NGS WGS dataset, SNVs were identified in both tumor and matched blood samples using Mutect2 from GATK ^107^ (version 4.1.2.0). The resulting variants were filtered using GATK’s FilterMutectCalls tool. CNVs in the WGS dataset were also detected using Control-FREEC with default parameters. Tumor purity was then estimated using both SNVs and CNVs with ABSOLUTE ^108^ (version 1.0.6), applying the following parameters: min.ploidy = 0.5, max.ploidy = 8, max.sigma.h = 0.2, max.neg.genome=0.005, sigma.p=0, max.as.seg.count=1500, and max.non.clonal=100.

### Somatic TRI Identification

Considering the high false-positive rate associated with detecting somatic SVs of the tandem repeat type using current algorithms, we have developed a more accurate method for identifying somatic tandem repeat insertions across pan-cancer samples (Extended Figure 2a). Our preliminary screening for potential somatic tandem repeat insertion sites involved the integration of two algorithms: Sniffles2 and Straglr, to pinpoint candidate regions. We initially employed Sniffles2 (version 2.0.7) with parameters --minsvlen 50 --mapq 5 --minsupport 3 to analyze tumor LRS WGS data, detecting INS SV intervals. These intervals were then intersected with tandem repeat regions annotated in the hg38 genome by RepeatMasker, utilizing the bedtools intersect command, with RepeatMasker-annotated tandem repeats as the initial candidate windows. Subsequently, we performed a secondary screen with straglr (version 1.4.1) using parameters --min_str_len 2 --max_str_len 100 --min_ins_size 100 --genotype_in_size --min_support 2 --max_num_clusters 9 on the tumor LRS WGS data, uncovering additional potential tandem repeat regions not identified by the sniffles2 workflow. These regions were added to the pool of candidate somatic tandem repeat intervals using the bedtools ‘intersect’ command. All candidate windows from each sample were further refined using the TDScope-simple command to efficiently filter for windows likely to contain somatic tandem repeat variations. We then consolidated all candidate somatic tandem repeat windows across samples using bedtools merge -d 200, yielding a comprehensive set of candidate windows for the pan-cancer cohort. Utilizing TDScope (version 1.0) with default parameters, we assessed these windows in both cancer samples and their corresponding normal blood samples. Candidate somatic tandem repeat intervals were identified by filtering for intervals with ≥3 supporting reads in normal blood samples, ≥3 supporting reads in tumor samples, an SV length (SVLEN) of ≥50, and a somatic genotype number = 1.

### Classification of Somatic TRI by Mutation Type

To annotate the consensus sequences of tumor and blood in TRI windows, we employed RepeatMasker, pyTRF (https://github.com/lmdu/pytrf), and mTR^109^. Due to the superior performance of pyTRF in annotating simple repeat sequences, we prioritized its annotation results. The remaining unannotated sequences were subsequently annotated using RepeatMasker and mTR. To eliminate inconsistencies between the different software tools, we normalized motifs of simple repeats, accounting for strand orientation and repeat unit size. This allowed us to obtain a consistent annotation of tumor and blood consensus sequences.

Pairwise alignment of the tumor and blood consensus sequences was performed to identify regions in blood characterized by gaps corresponding to insertions in tumor.

Annotation of the tumor insertion sequence was compared with all annotations of the blood sequence in the corresponding TRI window. If the tumor insertion motif matched the blood motif, the mutation type was classified as "Expansion." If the motifs differed, it was categorized as "Motif-Change." If no motif was annotated for the tumor insertion, the mutation type was classified as "Other." Using this approach, we determined the length of each mutation type in the tumor insertion sequence.

### Detection of Differential Somatic TRI Methylation Levels

To identify somatic TRI windows with significantly altered CpG patterns, we compared the CpG counts in all high-frequency TRI window pairs between somatic and germline sequences. A paired Wilcoxon rank-sum test was employed to assess differences in CG counts between somatic and germline samples. P-values were adjusted for multiple comparisons using the Benjamini-Hochberg method to calculate the false discovery rate (FDR) for each window. The relative change in CpG counts between somatic and germline sequences for each window was expressed as the log2 fold change (log2(FoldChange)), calculated as: log2FolgChange = log2(mean(somaticCG)+1 / mean(germlineCG+1)). High-frequency TRI windows with FDR ≤ 0.05 and |log2(FoldChange)| ≥ 1 were defined as significant CpG alteration windows, and these were used to compare methylation level changes in somatic sequences relative to germline sequences.

Specifically, reads from tumor samples classified by the TRIscope algorithm as ‘somatic support’ or ‘unsupport’ reads were aligned to reference genomes where somatic and germline sequences had been replaced, using Minimap2 (version 2.2.17) with the following parameters: --MD -ax map-ont -L. The methylation levels of somatic and germline TRI events were then calculated using the f5c pipeline (version 1.3, parameters: call-methylation -t 20 --iop 40).

To identify CpG insertion sites in somatic TRIs relative to germline TRIs, we used a pairwise alignment comparison algorithm (Python package: pairwise2 from the Biopython library). Based on these comparisons, we calculated methylation level changes for the inserted CpG sequences in somatic TRI events.

### Refinement Of High-confidence Somatic TEI Candidates

Samtools (version 2.8.0) was employed to tag BAM format files with the following command: samtools view -H “$bam_sort”; samtools view “$bam_sort” | awk ‘BEGIN{OFS=“\t”}{$1=“tag”$1; print $0}’) | samtools view -S -b - -o “$bam_sort_tag”. Following tagging, Samtools was again used to merge the tagged BAM files from tumor and paired blood samples (samtools merge -h “$blood_bam” “$merge_bam” “$blood_bam” “$tumor_bam”). Subsequently, SVs were identified using Sniffles2 (v2.0.6) based on the merged BAM files, which reported all supporting reads of each SV in the variant call format (VCF) file. Reads with a mapping quality below 10 were excluded from SV detection. To improve the accuracy of SV detection, a tandem repeat file (https://github.com/fritzsedlazeck/Sniffles/blob/master/annotations/human_GRCh38_no_alt_analysis_set.trf.bed) was provided using the --tandem-repeats parameter, enabling Sniffles to better resolve SVs involving repetitive sequences. SVs supported only by reads shorter than 1,000 base pairs (bp) were also discarded. Large-scale insertions were considered up to a maximum length of 10,000 bp. All other parameters were set to default values. Then, an initial set of candidate somatic SVs was filtered using a custom Python script with the following parameters: tumor reads supporting SV ≥ 3, blood reads supporting SV = 0, and SV length ≥ 50 bp.

To improve accuracy, an additional screening of candidate somatic SVs was performed. Specifically, SV positions were restricted to a standard deviation of less than 150 bp. For insertions (INS) and duplications (DUP), comparisons were made between the alignment-supported lengths and those reported by Sniffles, with outliers excluded based on:

A.mean(Lengtha)-2*STRI(Lengtha)<=Lengths <=mean(Lengtha)+2*STRI(Lengtha) B.Q1-1.5lQR<=Length:<=Q3+1.5IQR.

Additionally, SVs located in repeat regions were assessed for significant length differences between tumor and blood alignments, retaining only those exhibiting clear distinctions.

To further ensure high-confidence somatic SV identification, an error-correction process was conducted. Candidate regions were first identified as those within a ±50 bp range centered around insertion sites reported by Sniffles2. Alignments in these regions were extracted, and the reference genomes were polished using Racon (version 1.4.16, default parameters) over six consecutive iterations, utilizing paired blood reads from each region. Tumor and blood reads were then realigned to these polished reference genomes, and SVs were detected using Sniffles2. SVs supported by more than three tumor reads, with a length exceeding 50 bp, were retained as high-confidence somatic SVs.

### Somatic TEI Annotation

Iris is a software developed to improve the accuracy of insertion calling. We used Iris software (version 1.0.4, parameters: genome_buffer=1000 --keep_long_variants) to correct the positions and sequences of somatic SVs identified as INSs and DUPs. To improve the accuracy of Iris, we pre-processed the data by supplementing missing sequences for variants that Sniffles did not report. Specifically, we extracted the highest-quality reads from those supporting the Sniffles call that also spanned the insertion site. The insertion sequence from these reads was then identified using pysam and used as the input for Iris correction.

After error correction, we used RepeatMasker (version 4.1.2) to annotate the corrected insertion sequences with the following parameters: -species human -engine RMBlast -q, along with 50 bp of reference genome sequences upstream and downstream of the insertion site. If RepeatMasker failed to identify transposon regions, we performed re-annotation using Tandem Repeat Finder (TRF, version 4.10.0, parameters: 2 5 7 80 10 50 2000 -l 10 -ngs -h) and Sdust (version 0.1). Insertion sequences marked as “Unmask” by RepeatMasker were also re-annotated with TRF and Sdust if the annotated length exceeded 5 bp.

### Classification of Somatic TEI

If the 50 bp flanking sequences and the insertion sequence are not annotated as the same type of repeat and the insertion carries a transposable element (TE) sequence, the somatic INS is classified as a somatic TEI. Insertions containing TE sequences are further classified based on their structure and recombination mechanism. A single, complete insertion in the same orientation is classified as Solo, while insertions with multiple or fragmented TEs, or those with different orientations, are classified as Complex.

### Classification of SINE and LINE Truncations in Somatic TEIs

Each SINE and LINE insertion was annotated using RepeatMasker to compare the insertion sequences against template sequences in the RepeatMasker database. Using the positive strand as an example, we classified the insertion sequences based on their alignment characteristics. If the somatic insertion aligned fully with the template, with no gaps at either end, it was classified as Full Length. If the sequence matched entirely to the 3’ end (i.e., left = 0), it was categorized as a 5’ Truncation. Conversely, if it matched fully to the 5’ end (i.e., begin = 1), it was classified as a 3’ Truncation. If the sequence partially aligned with both the 3’ and 5’ ends but was not fully aligned at either end, it was classified as a TE Insertion. Additionally, the alignment coordinates of each SINE and LINE insertion against the template sequence were recorded for subsequent statistical analysis.

### Detection of SINE and LINE PolyA tails in Somatic TEI

For an insertion sequence on the negative strand, if there were 10 or more consecutive ‘A’ nucleotides within the first 30 bp of the sequence, it was classified as containing a polyA tail. Conversely, for an insertion sequence on the positive strand, the presence of 10 or more consecutive ‘T’ nucleotides within the last 30 bp was used as the criterion for classifying the sequence as containing a polyA tail.

### Detection of Somatic TEI Homologous Sequences

To identify homologous sequences between the ends of an insertion and the flanking regions of the reference genome, we performed BLAST analysis (version 2.14.1, parameters: -outfmt 6 -word_size 4), This analysis compared each insertion sequence with reference genomic sequences of varying lengths on both sides of the insertion site.

For gradient length determination, the maximum reference length was set equal to the length of the insertion, and the minimum length was defined by the allowable breakpoint deviation. For sequences shorter than 100 bp, we used length increments of 5 bp, while for sequences longer than 100 bp, increments were set to one-twentieth of the insertion length to optimize computational efficiency. Homologous sequences were defined as the longest segments within the breakpoint deviation that closely aligned with the ends of the insertion sequence.

Based on the relative position and length of these homologous sequences, insertions were classified into four categories:

1. Non-allelic homologous recombination (NAHR): Homologous sequences greater than 100 bp in length.
2. Tandem duplications (TSD): Homologous sequences less than 100 bp in length with a distal relative position.
3. Recombination (Else): Homologous sequences that did not meet the criteria for NAHR or TSD.
4. No homologous sequence detected: Insertions were classified as "N" when no homologous sequence was identified.

### Somatic TEIs Motif Clustering

To examine the distribution patterns of somatic TEIs across the genome, we divided the genome into 1 Mb fixed-size windows to create a pan-cancer atlas of somatic TEI distribution. The genome was segmented into consecutive 1 Mb windows, and the presence of somatic TEIs within each window was recorded for each tumor sample. This resulted in a binary matrix representing the presence or absence of somatic TEIs across the genome for each sample.

Windows containing TEIs in only a single tumor sample were excluded from further analysis. For the remaining data, we calculated the Jaccard similarity coefficient for each pair of tumor samples to assess their genomic TEI similarity. Hierarchical clustering was then performed using the pheatmap package in R (ward.D method) to group tumor samples based on their somatic TEI distribution.

### Analysis of the Association Between L1 Insertion Rate and Genomic Features

The L1 insertion rate is determined by calculating the total number of somatic L1 insertions (10,140) identified in 317 out of 322 samples within each 1Mb window. To investigate the association between L1 insertion rates and multiple predictor variables, we employed a statistical framework based on negative binomial regression1. Specifically, we categorized the genome into four bins (bin0 - bin3) based on the degree of match between the classic L1 endonuclease motif (defined as TTTT|R, where R is A or G or Y|AAAA, where Y is C or T) and the DNA sequence. Bin0 contains DNA motifs that have four or more mismatches (out of five possible), which are highly dissimilar. Bin1, bin2, and bin3 include genomic segments of the hg38 genome assembly with three, two, and at most one mismatch, respectively. The most tightly matching DNA strands at each site are considered.

Regional analysis of genomic features utilized histone mark data (H3K9me3, H3K4me3, H3K36me3, H3K27ac ChIP-Seq data) and DNase hypersensitivity (DHS) data from the Roadmap Epigenomics Consortium. These data were divided into four bins based on the average enrichment scores across eight cell types (E017, E114, E117, E118, E119, E122, E125, E127). Bin0 includes regions with enrichment scores below baseline (Roadmap enrichment scores relative to input <1), while bins 1 to 3 correspond to regions with higher-than-average enrichment scores across the genome. Specifically, the DHS bins 1-3 span 132.1-136.1 Mb, the H3K9me3 bins 1-3 span 192.8-198.7 Mb, the H3K36me3 bins 1-3 span 142.3-146.5 Mb, the H3K4me3 bins 1-3 span 44.3-45.9 Mb, and the H3K27ac bins 1-3 span 77.8-80.1 Mb.

Replication timing (RT) data from ENCODE were processed by averaging smoothed waveform signals from eight cell types (IMR-90, HeLa S3, HUVEC, BJ, HEP G2, NHEK, SK-N-SH, and MCF-7). The data were divided into four equal-sized bins (quartiles), where bin0 represents the latest replication and bin3 represents the earliest replication.

RNA-Seq data, also from the Roadmap, were processed similarly. By averaging data from eight cell types (E071, E096, E114, E117, E118, E119, E122, E127), the data were divided into four genomic bins: bin0 contains non-expressed genes (logRPKM=0) and non-expressed intergenic DNA (totaling 1173.9 Mb), while bin1 (maximum logRPKM=0.02), bin2 (maximum logRPKM=0.15), and bin3 (logRPKM > 0.15) span 353.7 Mb, 302.2 Mb, and 382.9 Mb, respectively.

In all figures (Supplementary Fig. 3b), the enrichment scores compare differences in specific features (replication timing, histone marks, gene expression, and L1 motifs) between bins 1-3 and bin0. The logarithmic enrichment for bin0 is defined as 0, and thus it is not displayed in the enrichment plots. Regional analysis is limited to genomic regions with perfect alignment scores, based on the hg38 reference genome.

### Fresh tissue preparation and dissociation into single cells

All the freshly resected surgical specimens were immediately washed with phosphate-buffered saline (PBS) and processed to generate single cell suspensions. Tissue digestion was performed in 15 ml tube containing 10 ml pre-warmed RPMI 1640 (ThermoFisher Scientific), 2 mg/ml dispase (Roche), 1 mg/ml type IV collagenase (Sigma) and 10 U/µl DNase I (Roche) for 60 minutes at 37°C. The reaction was deactivated by 10% FBS. Cell suspensions were filtered using a 70 μm filter and then centrifuged at 500 rpm for 6 min at 4°C to pellet dead cells and red blood cells. The cells were washed twice and suspended in PBS with 0.5% bovine serum albumin (BSA, Sigma), and, used CD45 MicroBeads (Miltenyibiotec, 130-045-801) to distinguish CD45+ cells from the rest of the cells.

### Validation of the Somatic TEI&TRI in Tissues and Cell Lines using PCR and Sanger sequencing

DNA extraction and PCR: DNA was extracted from tumor tissues, blood, LUSC cell lines (H226, H520, HCC95), LUAD cell lines (PC-9, A549, H1299 and H1975), IC cell lines (LOVO, SW620, RKO, HCT15, HCT116,T84), Breast cancer cell line (HCC1395), OSCC cell lines (TU212, TU686, SNU46, SNU899), CCA cell lines (H8, C33A, CASK, SiHA), PAAD cell lines (HPAC, PANC-1,BXPC-1,ASPC-1),HCC cell lines (sun387, Huh7, hcclm9, snu499,sk-hep1), the H1581 cell line, Normal cell line BEAS-2B, HCC1395BL using the DNeasy Blood & Tissue Kit (QIAGEN) according to the manufacturer’s instructions. The forward and reverse primers were designed according to the 100 bp upstream and downstream sequences of the insertion position and synthesized by Sangon Biotech (Table TEI,Table TRI). DNA was extracted from clinical samples or cell lines and subjected to PCR using primers. Each 50-μL reaction consisted of 2× PremSTAR Max Premix 25 μL, 0.8 μL 10 μM anchor primer, 0.8 μL 10 μM test primer, and 50-150 ng DNA. PCR was performed as follows: an initial step of 2 minutes at 98 °C, 30 cycles of 10 seconds at 98 °C, 15 seconds at 60 °C and 60-300 seconds at 72 °C (size of PCR products between 200 bp and 6 kb). Finally, the samples were incubated at 72 °C for 5 minutes. The PCR products were sent to Sangon for Sanger sequencing.

### Cell Culture

Human cell lines provided by the American Type Culture Collection (Manassas, VA, USA) were used in the present study. The cells were incubated in Dulbecco’s modified Eagle’s medium (DMEM) or RPMI 1640 medium (Gibco; Thermo Fisher Scientific, Inc.). The medium was supplemented with 10% fetal bovine serum (FBS) (Gibco; Thermo Fisher Scientific, Inc.), and the cells were all maintained at 37 °C in 5% CO2. Upon reaching 70-80% confluence, the cells were washed with PBS and detached with 0.25% trypsin/0.2% ethylenediaminetetraacetic acid (EDTA). Cell morphology was viewed under a light microscope, and the cells were suspended to a concentration of 1 x 106 cells/ml.

### Immunofluorescence and FISH

Briefly, 10 μm thick frozen sections were treated with 0.2% Triton X-100 in PBS for 15 minutes and blocked with 10% serum in PBST for 40 minutes at room temperature. They were then incubated with primary antibodies, Anti CK7/serpinB3/NES/GFAP (1:500, Huabio, ET1609-62, ET7111-21, ET1610-96, ET1601-23) overnight at 4°C and washed with PBS for 15 min, Alexa-conjugated secondary antibodies (1:1000, iFluor 488 Goat anti-rabbit IgG antibody, Huabio, HA720120F; iFluor 594 Conjugated Goat anti-rabbit IgG polyclonal Antibody, Huabio, HA1122) were incubated for 1 hour at room temperature. For FISH, probes specific to the target sequences (Table S20) were hybridized to the cells overnight at 37°C. After washing, the cells were counterstained with DAPI and analyzed under a fluorescence microscope.

### Cell co-culture

Splenic cells were collected from 6-8 weeks old C57/BL6 mice and resuspended in RoboSep™ Buffer (STEMCELL Technologies, 20104). Total splenocytes were enriched for CD4+ T cells with EasySep™ Mouse CD4+ T Cell Isolation Kit (STEMCELL Technologies, 19852). The selected CD4+ T cells and tumor cells were directly co-cultured with RPMI 1640 medium (containing 10% FBS) at a ratio of 10:1 for 24 hours, and suspended T cells were collected at 24 hours for subsequent flow experiments.

### Flow cytometry

For cell surface staining, cells were stained with the corresponding antibodies, APC anti-mouse CD45 (1:100, Biolegend, 103112); FlTC anti-mouse CD4 (1:100, Biolegend, 100405); PE anti-mouse CD154 (CD40L) (1:50, Biolegend, 157003); eBioscience™ Fixable Viability Dye eFluor™ 506 (1:100, Thermo Fisher Scientific, 65-0866-18), in the presence of Fc receptor blocker on ice for 30 min, and then fixed with 4% paraformaldehyde for 10 minutes, washed with 1x PBS solution, and resuspended after centrifugation. Samples were assessed by a spectral flow cytometer (Cytoflex, BECKMAN Biosciences,) equipped with SpectroFlo software (Cytek Biosciences). Acquired flow cytometry results were analyzed by FlowJo software.

## Data availability

All sequencing data are available at the Genome Sequence Archive for Human: https://ngdc.cncb.ac.cn/gsa-human/s/QfWwWORT, https://ngdc.cncb.ac.cn/gsa-human/s/d97GuJ2L and https://ngdc.cncb.ac.cn/gsa-human/s/MnWe8KbN.

## Code availability

All code used is open source and available at github: https://github.com/LinXialab/TGS_somatic_TEI/ and https://github.com/Goatofmountain/TDScope.

## Acknowledgments

We thank members of the Dan Xie lab for useful discussions. This work was supported by the National Natural Science Foundation of China 82173383 and 82241238 (to D.X.), 32200508 (to L.X.), 32300522 (to K.L.T), 32300003 (to H.W.), 82371883 (to J.F.L.), 82472365 (to H.T.L.), the Sichuan Province Science and Technology Program 2025ZNSFSC1098 (to K.L.T.), and the 1·3·5 project for disciplines of excellence, West China Hospital, Sichuan University ZYYC23024 (to D.X.).

## Author contributions

Conceptualization: D.X.; Data Curation: W.L., Z.F.W., Z.W., C.L., J.F.L., Y.X.Q., Q.W.Z., H.T.L., S.Z.H., Z.Y.Z., A.L., A.L.W.; Software, Formal Analysis and Methodology: L.X., K.L.T., X.Y.L., Q.L.Z., R.L.W., Z.T.F., X.N.P.; Validation: H.W., M.L.L., Y.H., J.W.; Supervision: D.X.; Writing – Original Draft: D.X., L.X., K.L.T., H.W., X.Y.L., Y.H.L., C.Y.Y., Z.T.F.; Writing – Review & Editing: all authors.

---

**Figure S1.**
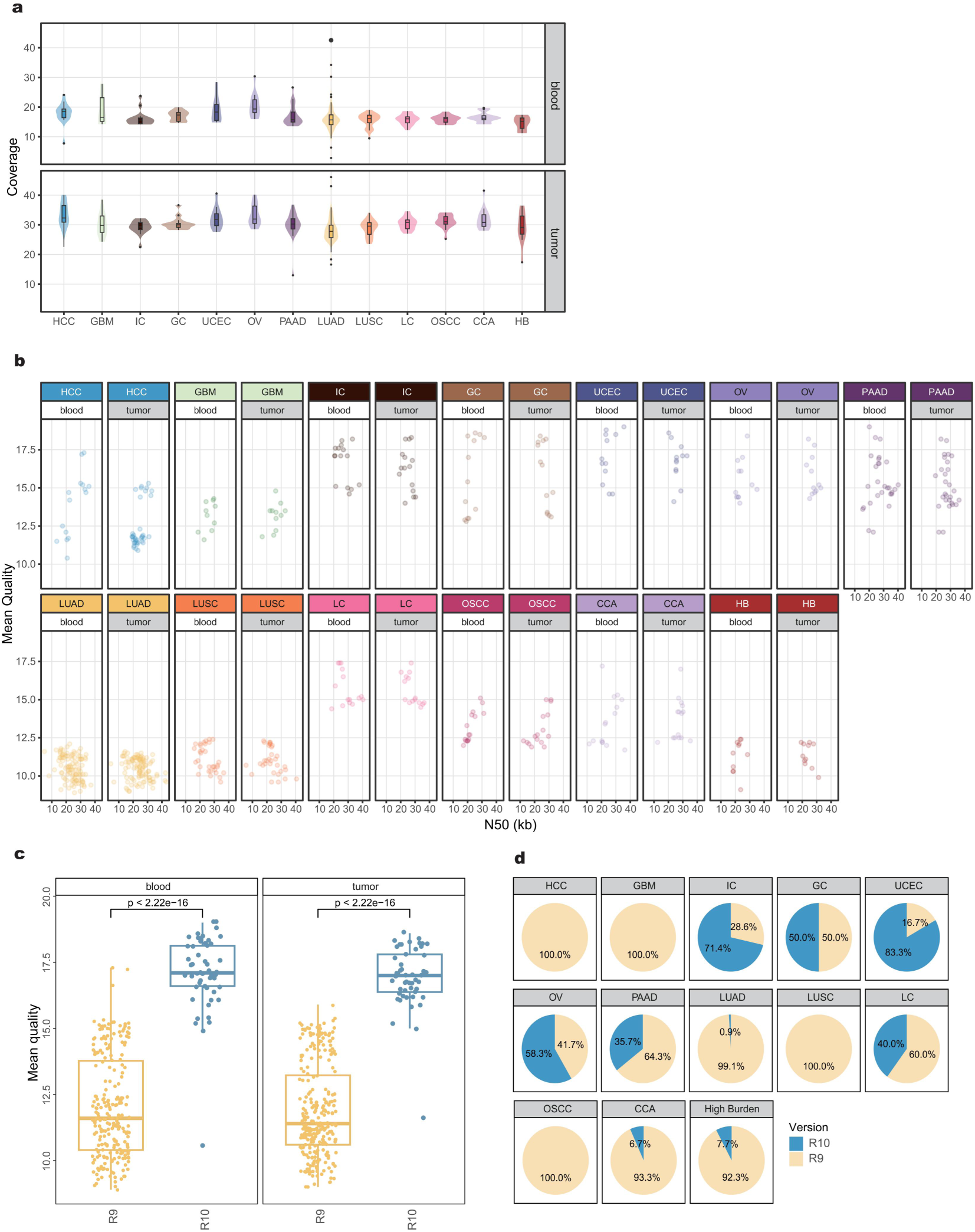
Summary of long-read sequencing data. a. Sequencing depth of long-read data from 325 paired tumor (below) and paired blood (above) samples across 12 cancer types and high-burden samples. b. Scatter plots showing the N50 and mean quality of LRS reads derived from paired tumor (gray) and blood (white) samples, spanning 12 cancer types and high-burden samples. Each dot represents a sample. c. Box plots comparing sequencing quality between R9 (blue) and R10 (yellow) chips, with statistical significance assessed by the Wilcoxon rank test. Each dot represents a sample. d. Pie chart illustrating the distribution of sequencing chip types used across various cancer types and high-burden samples. Each color represents a different chip version.

**Figure S2.**
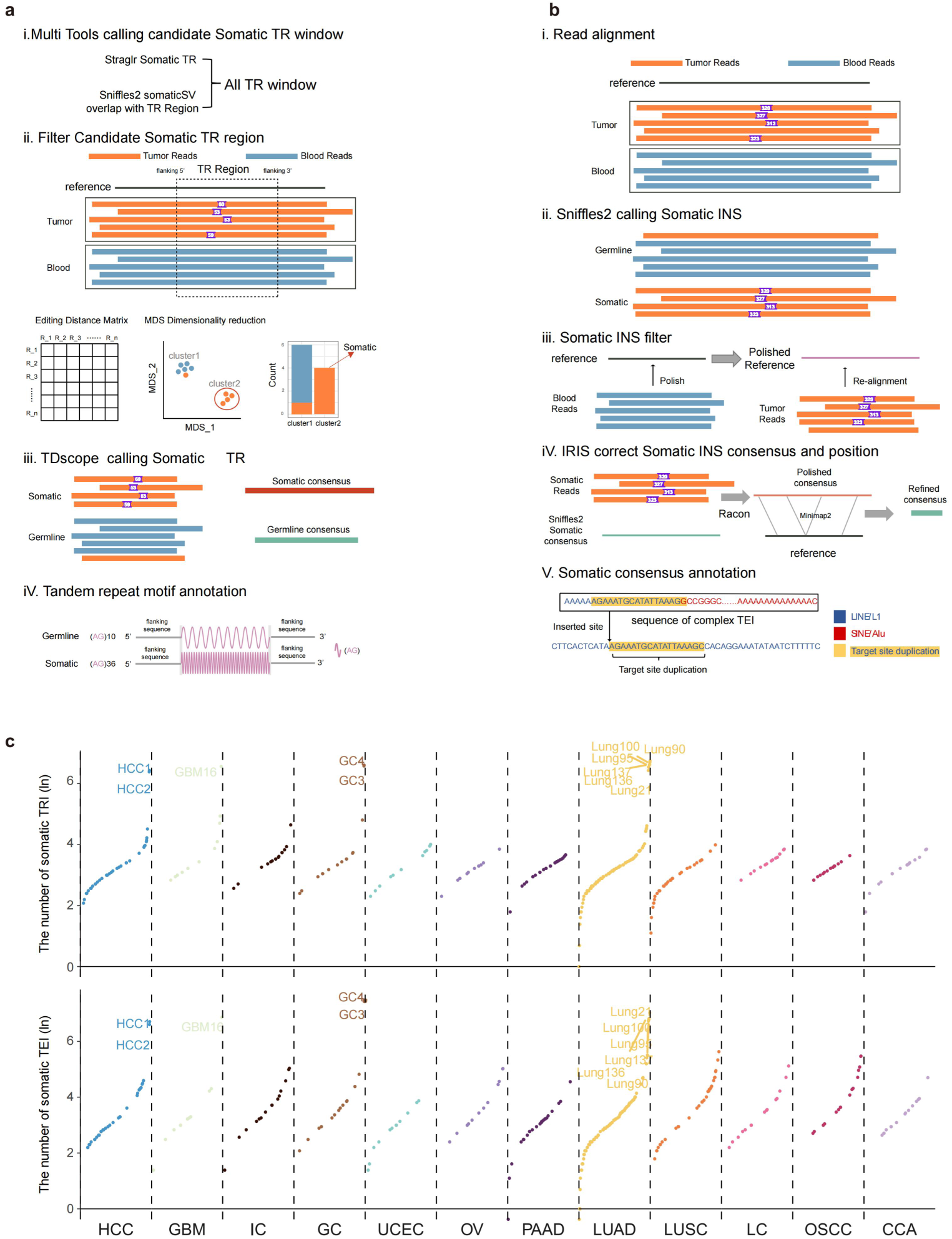
Overview of Methods for Identifying Somatic Genomic Alterations and The Frequency of Somatic TRIs/TEIs. a. Schematic representation of the workflow for identifying somatic TRI. i. Candidate somatic TRI regions are identified using Sniffles2 and Straglr; ii. Reads spanned these regions are extracted, and edit distances are calculated between sequences spanned potential somatic TRI region, followed by clustering to initial screen the presence of somatic TRIs; iii. Somatic TRIs are confirmed using TDscope; iv. Repeat motif of somatic TRIs are annotated (see Methods for details). b. Schematic representation of the workflow for identifying somatic TEI. i. Reads from tumor and blood samples are aligned to the reference genome; ii. Candidate somatic SVs are identified using Sniffles2.; iii. After polishing the reference genome, reads are realigned to the refined reference to validate the presence of candidate SVs; iv. Somatic SVs are further refined using IRIS; v. Somatic SVs are annotated (see Methods for details). c. The upper panel shows the frequency (log10 scale) of somatic TRIs, while the lower panel displays the frequency of somatic TEIs across various cancer types. Each dot represents an individual sample, with distinct colors indicating different cancer types. Samples with high burden of TRIs or TEIs are labeled accordingly.

**Figure S3.**
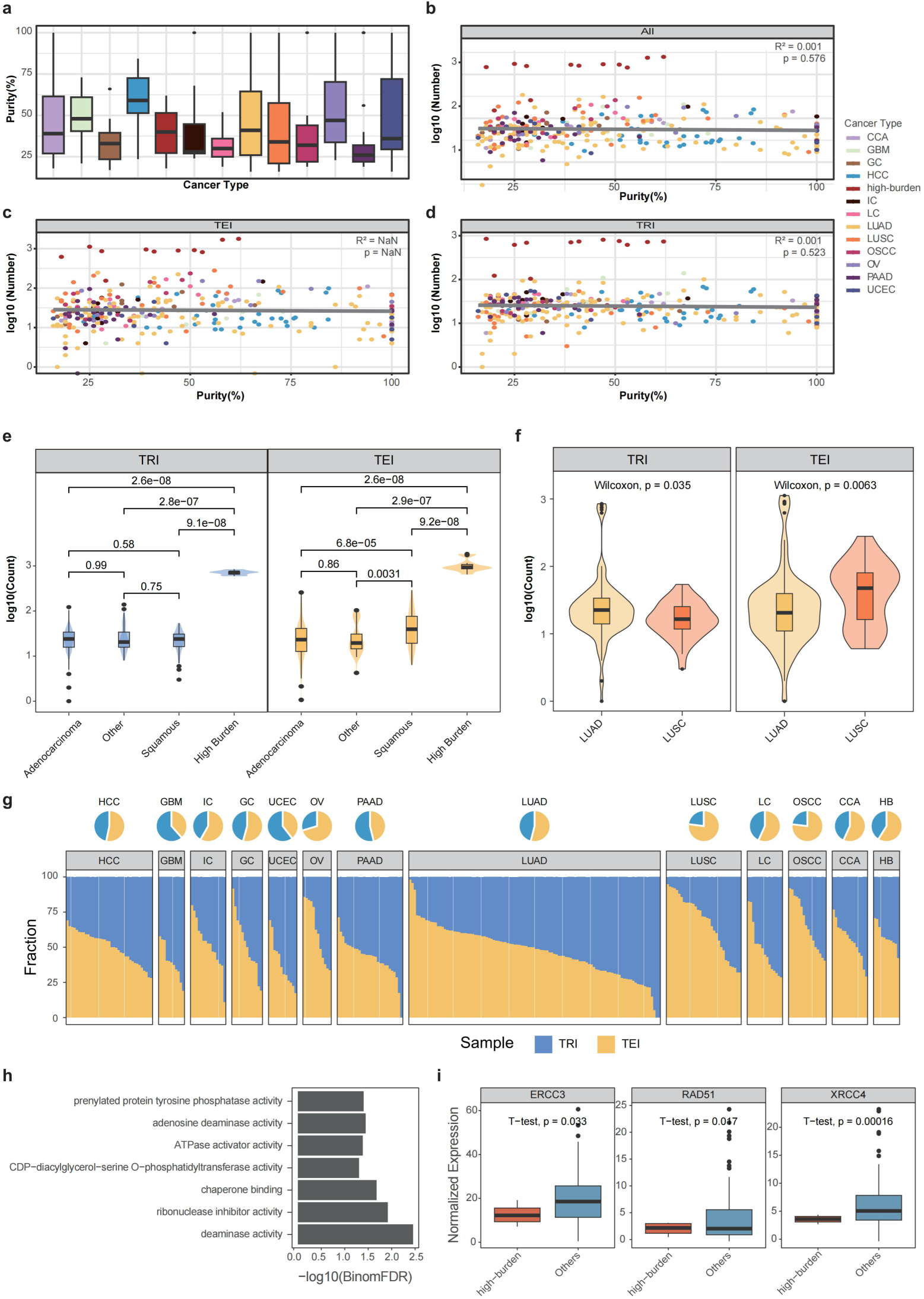
Tumor Purity and Characteristics of Somatic REIs. a. Box plot of tumor purity across different cancer types. Each box represents a specific cancer type, with the y-axis indicating tumor purity percentage. b-d. Scatter plots showing the correlation between tumor purity and the count of somatic TRIs (e), somatic TEIs (f), and the combined count of somatic TRIs and TEIs (d). Data points represent individual samples, colored by cancer type. Pearson correlation coefficients (R²) and p-values are displayed in the upper-right corner of each plot. Boxplot showing the log-transformed number of somatic TRIs (blue) and somatic TEIs (yellow) across different cancer types: "Other" (including HCC, GBM), Squamous (LUSC, CCA, OSCC, LC), and Adenocarcinoma (LUAD, PAAD, GC, IC, OV, UCEC). High-burden samples are indicated separately. Statistical significance is assessed using a two-tailed t-test. e. Boxplot illustrating the log-transformed number of somatic TRIs (blue) and somatic TEIs (yellow) in Adenocarcinoma, Squamous, Other and high-burden samples. p-values were calculated using the Wilcoxon rank-sum test. f. Boxplot illustrating the log-transformed number of somatic TRIs and somatic TEIs in LUSC (yellow) and LUAD (orange) samples. p-values were calculated using the Wilcoxon rank-sum test. g. A bar chart depicting the proportion of somatic TEIs (yellow) and somatic TRIs (blue) across cancer types and high-burden samples. Each bar represents a distinct sample. A pie chart above summarizes the overall percentage of somatic TRIs and TEIs within each cancer type. h. Bar plot showing the GO Molecular Function enrichment analysis of genes associated with high-burden specific TRIs, as predicted by the GREAT. The x-axis represents the −log10 transformed FDR from the binomial test. i. Boxplots comparing the expression levels of *ERCC3*, *RAD51*, and *XRCC4* in high-burden LUAD samples (red) and low-burden LUAD samples (blue). Statistical significance was assessed using a two-tailed t-test.

**Figure S4.**
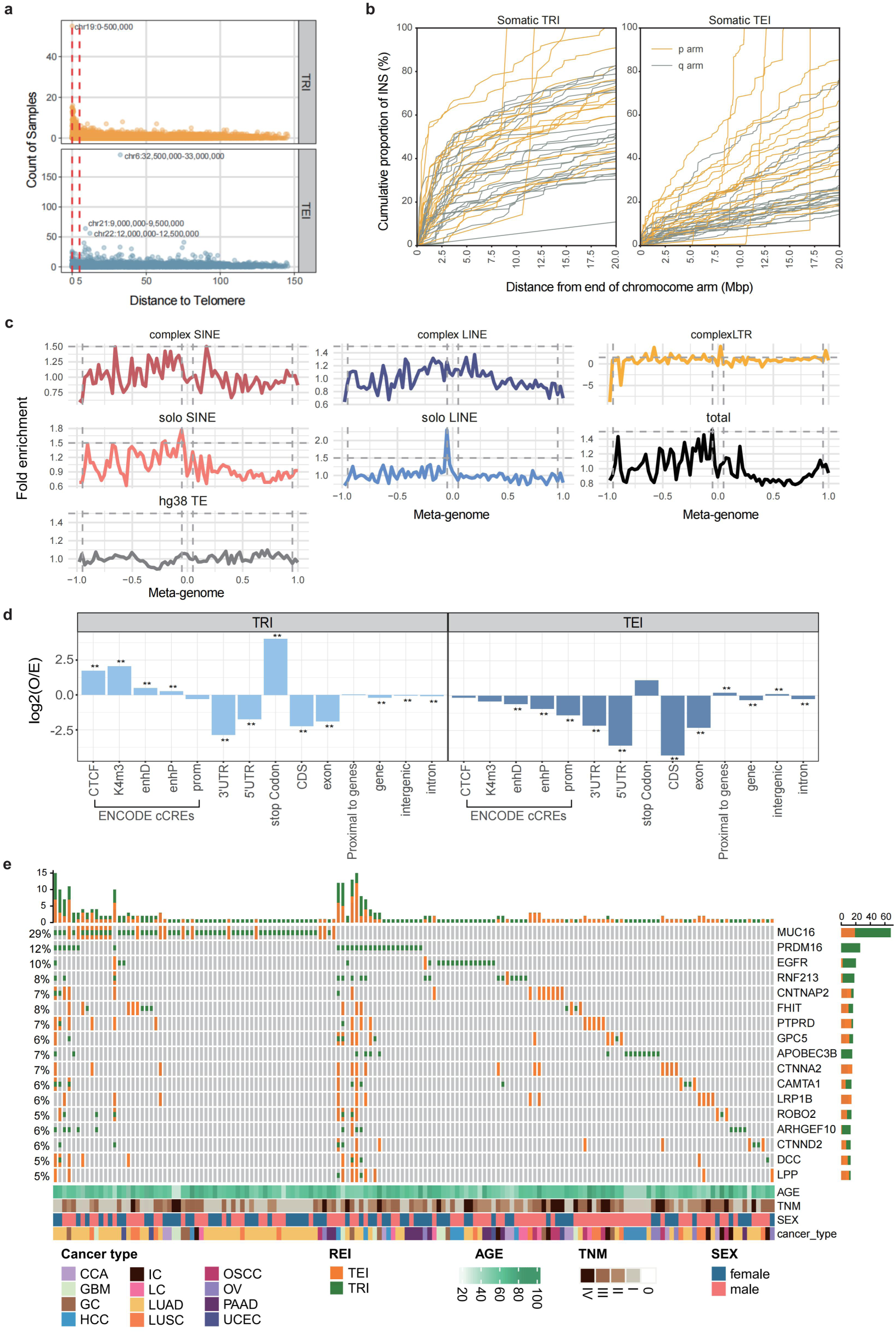
The genome distribution of somatic TRIs and TEIs. a. Distribution of the distance from somatic TRIs (yellow, above) and somatic TEIs (blue, below) to telomeres. The x-axis represents the distance from somatic insertions to the telomeric ends, while the y-axis indicates the number of samples. Two red dashed lines mark the positions of 0 bp (telomere ends) and 5 Mbp from the telomere ends. b. Cumulative distribution of somatic TRI (left) and somatic TEI (right). The plot reveals a more rapid saturation of somatic TRIs at the ends of chromosome arms. A sliding window of 500,000bp, starting from the telomeric regions and extending toward the centromere, was used to cumulatively count somatic TRIs and TEIs. The x-axis is truncated at 20 Mbp. c. Chromosome-level distribution of various somatic TEI types ("observed") compared to TEs in the hg38 reference genome ("expected"). The fold enrichment of observed TEIs relative to random expectations is calculated refer to the previous study ^58^. Each density plot corresponds to a distinct type of TEI, represented by different colors. d. Enrichment of somatic TRIs (left) and somatic TEIs (right) on genomic elements. The ratio of observed insertions to random expectation (O/E) is calculated using a hypergeometric test. *p < 0.05. The y-axis is presented on a log scale for O/E values. Somatic TRIs and TEIs are shown in different colors to distinguish between the two insertion types. e. Oncoplot shows high frequency (Sample frequency > 5%) COSMIC cancer genes with somatic REI events located within gene body and 2kbp flanking regions. The bar plot on the top and right show the number of somatic REI events located within samples and gene flank regions with color coded in the bottom,

**Figure S5.**
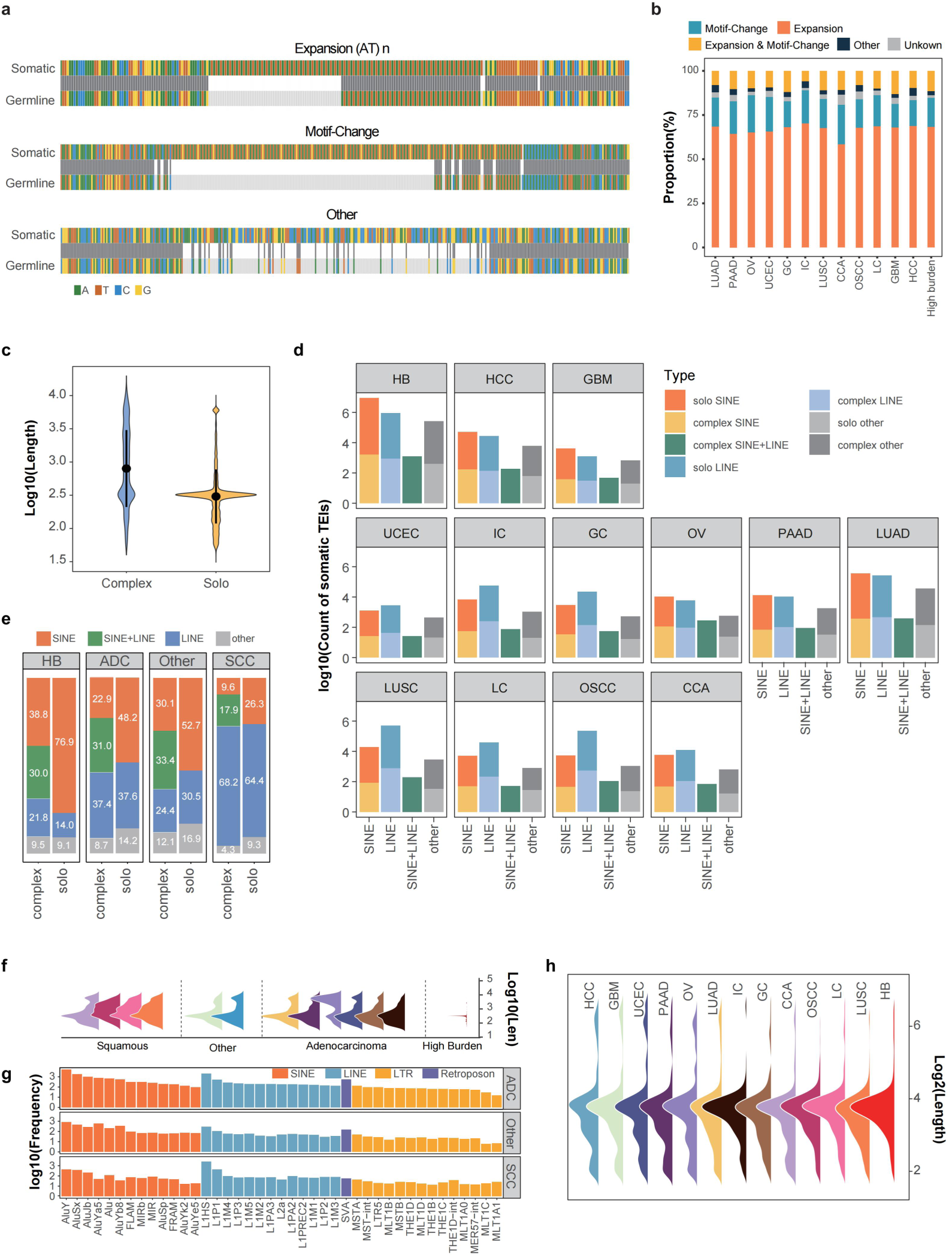
Sequence charateristic of somatic TRIs and somatic TEIs. a. Schematic representation of somatic TRI patterns for Expansion, Motif Change, and Other. The top row represents somatic sequences, the middle row indicates aligned/unaligned, and the bottom row shows germline sequences. b. Proportions of each somatic TRI category in cancer types: Expansion (orange), Expansion & Motif-Change (yellow), Motif-Change (blue), Ambiguous (dark blue) and Unknown (gray). c. Violin plot displaying the log-transformed length distribution for somatic complex (blue) and solo (yellow) TEI. d. Proportions of somatic complex and solo TEI across various cancer types. ‘complex SINE’ indicates complex TEIs containing SINE elements but lacking LINE elements; ‘complex LINE’ indicates complex TEIs containing LINE elements but lacking SINE elements; ‘complex SINE+LINE’ denotes TEIs containing both SINE and LINE elements. e. Frequency (log10) of SINE (orange), LINE (blue), LTR (yellow) and Retroposon (purple) subfamilies in adenocarcinoma, squamous cell carcinoma and other cancer type. Each column represents a subfamily. ‘ADC’: adenocarcinoma, ‘SCC’: squamous cell carcinoma, ‘other’: HCC and GBM. f. Length density distribution (log10 scale) of somatic TEIs in squamous cell carcinoma, other cancer types, adenocarcinoma, and high-burden samples. Different colors represent different cancer types, similar to panel h. g. Log-transformed counts of different subfamilies of somatic TEIs across cancer types. "ADC" represents adenocarcinoma, "SCC" stands for squamous cell carcinoma, and "other" includes HCC and GBM. h. Density plot of log-transformed length distribution of TSD across different cancer types.

**Figure S6.**
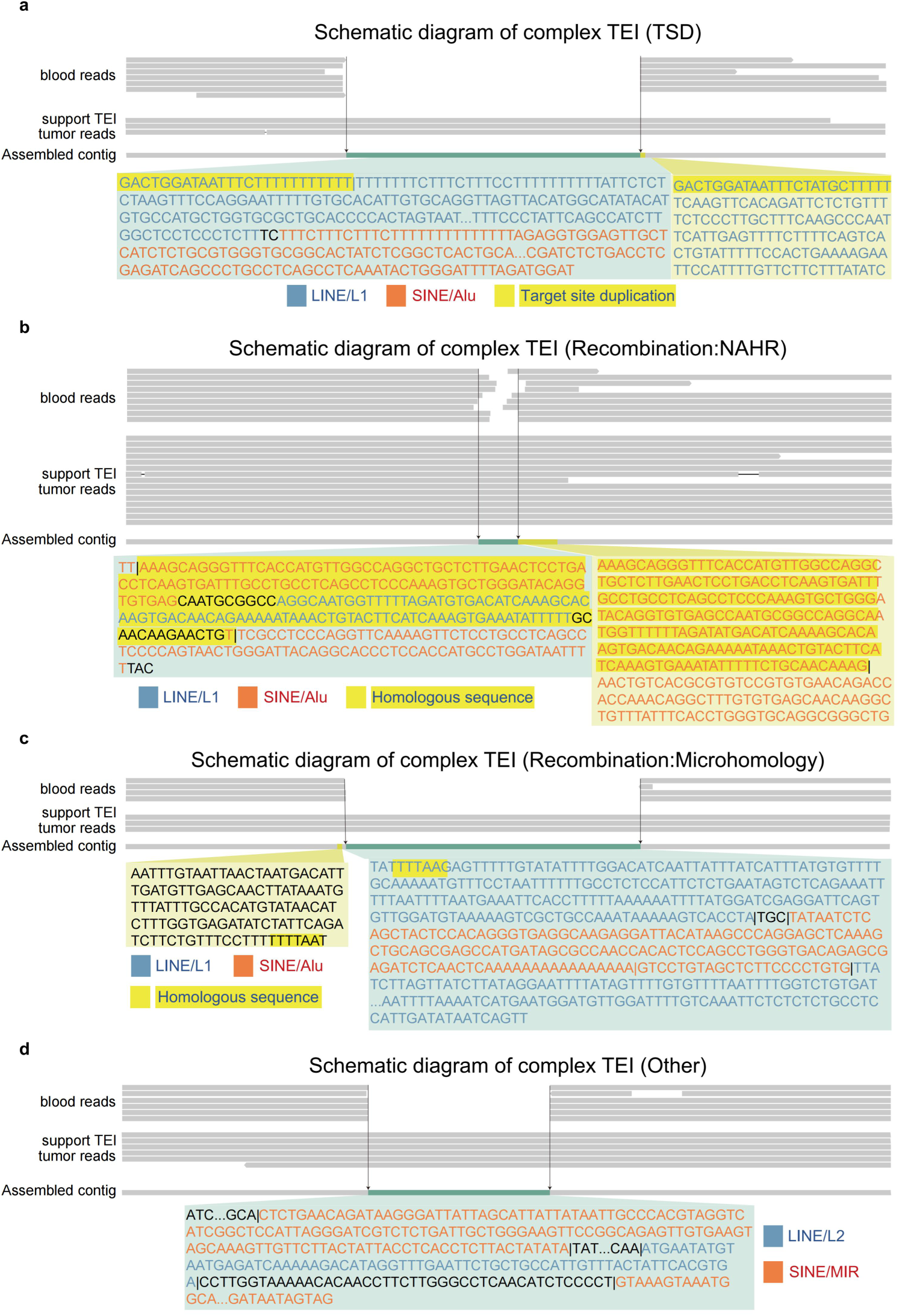
Examples of somatic TEI insertion mechanisms. a. The schematic illustrates example of Target Site Duplication. In IGV, the reference genome comprises assembled sequences containing TEI-supporting reads, with all blood reads and tumor TEI-supporting reads aligned to this assembly. Complex TEI sequences are highlighted in light green, and homologous sequences are marked in yellow-green. LINE elements are colored in blue, while SINE elements are colored in orange-red. Target site duplications are marked in bright yellow. b. The schematic illustrates example of Recombination (NAHR). In IGV, the reference genome comprises assembled sequences containing TEI-supporting reads, with all blood reads and tumor TEI-supporting reads aligned to this assembly. Complex TEI sequences are highlighted in light green, and homologous sequences are marked in yellow-green. LINE elements are colored in blue, while SINE elements are colored in orange-red. Homologous sequences are marked in bright yellow. c. The schematic illustrates example of Recombination (Microhomology). In IGV, the reference genome comprises assembled sequences containing TEI-supporting reads, with all blood reads and tumor TEI-supporting reads aligned to this assembly. Complex TEI sequences are highlighted in light green, and homologous sequences are marked in yellow-green. LINE elements are colored in blue, while SINE elements are colored in orange-red. Homologous sequences are marked in bright yellow. d. The schematic illustrates example of Other. In IGV, the reference genome comprises assembled sequences containing TEI-supporting reads, with all blood reads and tumor TEI-supporting reads aligned to this assembly. Complex TEI sequences are highlighted in light green, and homologous sequences are marked in yellow-green. LINE elements are colored in blue, while SINE elements are colored in orange-red.

**Figure S7.**
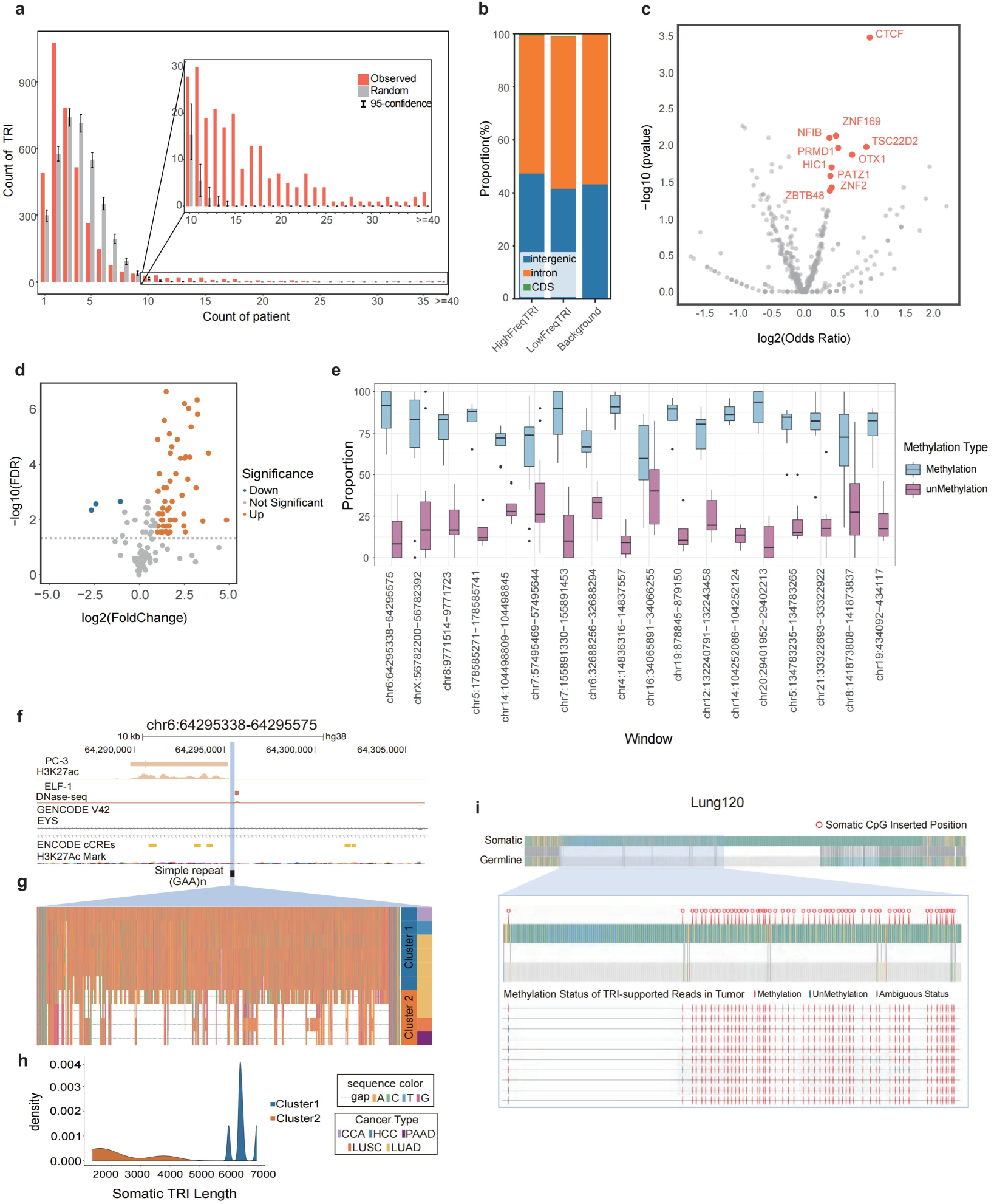
Overview of TRI distribution. a. Bar plot depicting the number of patients sharing the same somatic TRI events. Gray bars indicate the random distribution (Method), while orange bars represent the observed distribution. The x-axis represents the number of patients sharing somatic TRIs, and the y-axis represents the count of TRIs. b. Bar plot illustrating the percentage distribution of low-frequency, high-frequency, and background TRIs located within intergenic, intronic, and coding sequence (CDS) regions. c. Volcano plot depicting the differential enrichment between high and low frequency somatic TRIs on known 490 TF binding regions from ENCODE project. Elements with p-value < 0.05 (Chi-square test) and |log2(Odds Ratio)| > 1 are highlighted in red. d. Volcano plot illustrating high-frequency TRIs with significant changes in CpG counts between somatic and matched germline TRI sequences. Orange: TRIs with FDR < 0.05 and log2(FoldChange) > 1 (Wilcoxon test); Blue:FDR < 0.05 and log2(FoldChange) < 1 (Wilcoxon test). e. Box plot displaying the proportions of methylated and unmethylated samples at 18 high-frequency somatic TRIs introduced methylation marks at generated CpG dinucleotides (Wilcoxon test, p < 0.05). Each dot representing the proportion of methylated (blue) and unmethylated (pink) states for each sample within each TRI window. f. Browser plot displaying the epigenomic signal, ENCODE-annotated gene transcript, and cCREs (Candidate Cis-Regulatory Elements) predicted by ENCODE around the somatic TRI located in the intron of the EYS gene. g. An enlarged panel providing a detailed sequence overview of different clusters surrounding the somatic TRIs in the EYS intron. h. Density plot depicting the length distribution of two clusters, color-coded for clarity. i. Methylation analysis of the TRI sequence from a representative sample: the top panel displays overall sequences of somatic and corresponding germline with pairwise alignment details; the middle panel highlights the specific somatic CpG insertion site; the bottom panel shows all tumor reads supporting the TRI event, along with methylation status at the somatic CpG insertion site. Methylated CpG sites are colored in red and unmethylated CpG sites are colored in blue.

**Figure S8.**
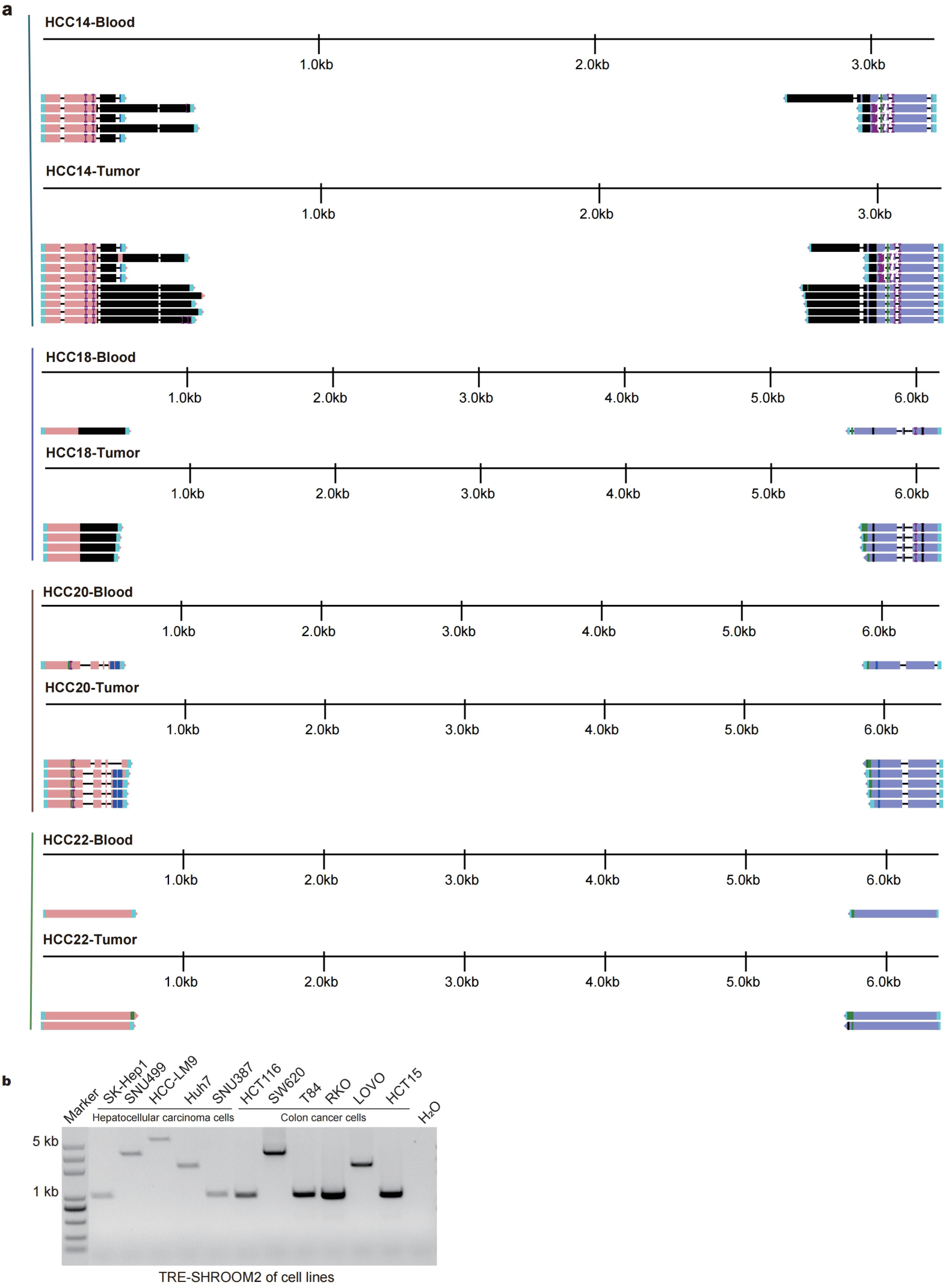
Sanger sequencing of Shroom2-related SRE in clinical samples and detection in cell lines. a. Visual display of sanger sequencing of Shroom2-related SRE in clinical samples. b. Detection of Shroom2-related SRE in liver cancer and intestinal cancer cell lines.

**Figure S9.**
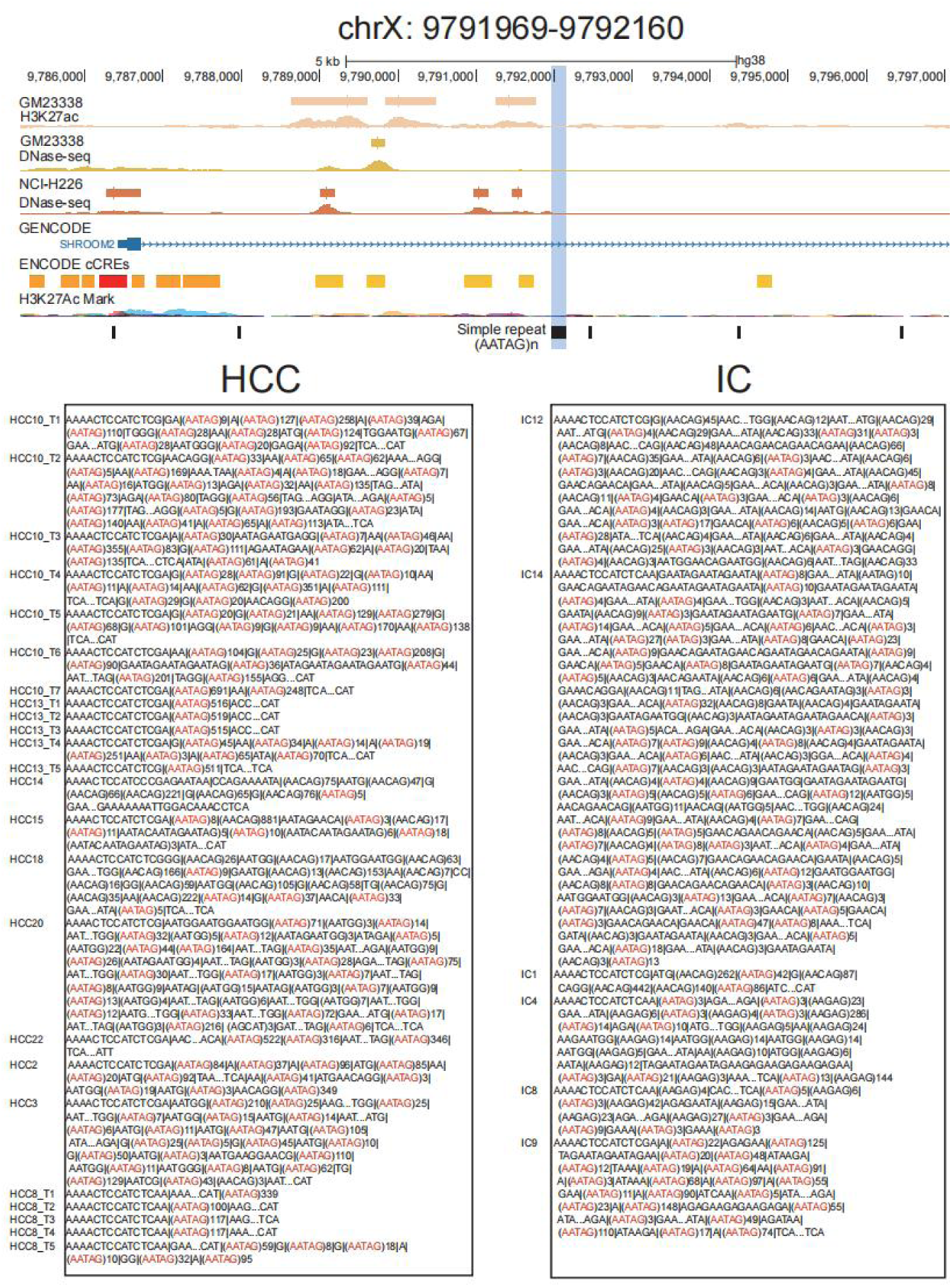
The browser plot of somatic TRIs located in the intron of *SHROOM2*, along with the somatic TRIs sequence displays for HCC and IC. The browser plot illustrates epigenomic signals from ENCODE project, including cell line H3K27ac chip-seq and DNase signal, gene transcript Candidate Cis-Regulatory Elements (cCREs), and H3K27ac marks identified by ENCODE.The somatic TRI (chrX: 9791969-9792160) is highlighted in blue. The bottom panel illustrates detailed somatic sequences of HCC and IC samples harbored the somatic TRI event with the motif AATAG abbreviated and highlighted in red.

**Figure S10.**
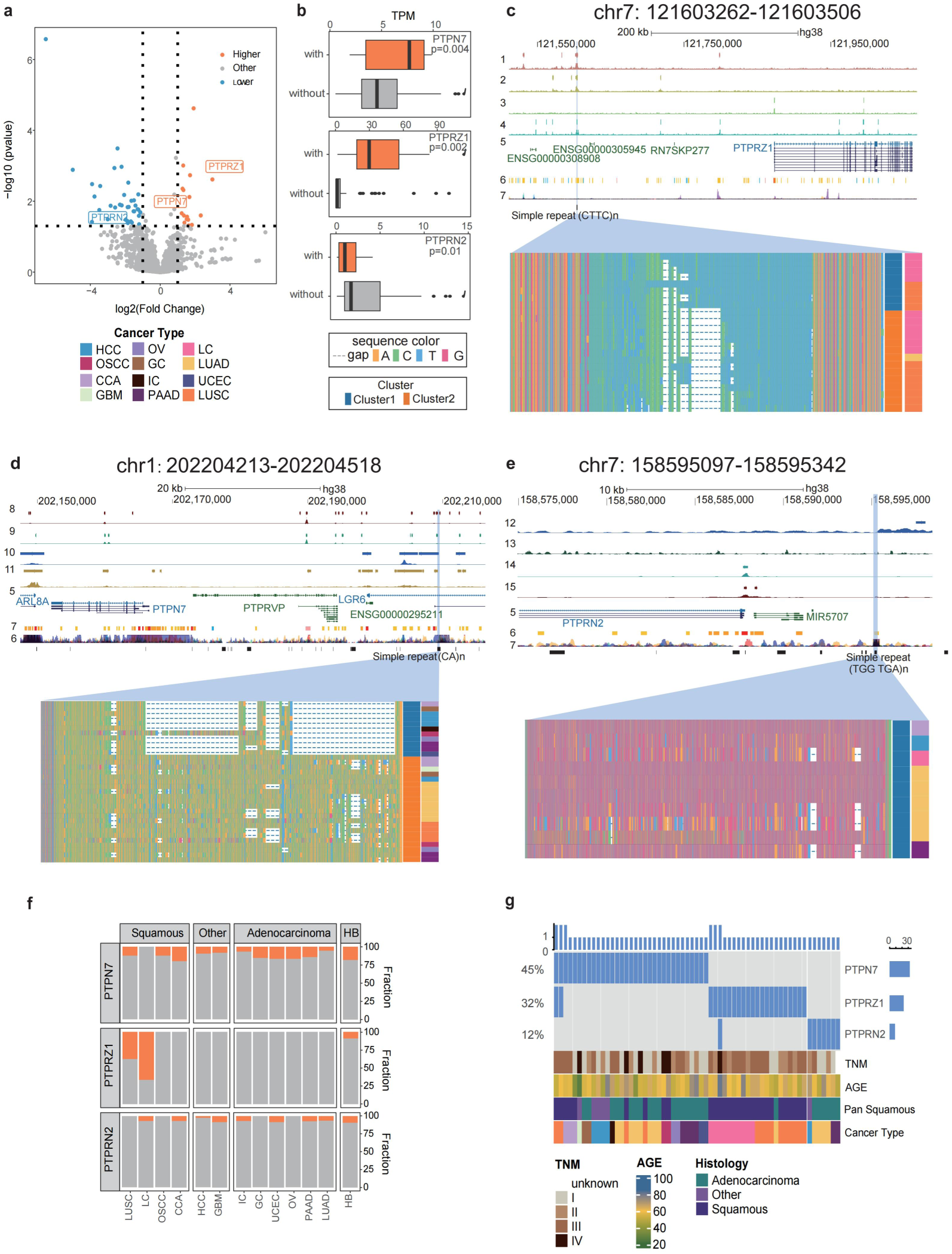
Three somatic TRIs that associated with the expression of PTP family genes in NSCLC. a. Volcano plot showing differentially expressed genes between NSCLC samples with and without high-frequency somatic TRIs. Genes from the PTP family are highlighted with text. Dotted lines parallel to the X-axis indicates where p = 0.05, while two vertical dashed lines represent log2(Fold change) values of −1 and 1, respectively. b. Box plot of the expression levels of differentially expressed PTP family genes with and without somatic TRI. Orange: with TRI; Gray: without TRI. P values were calculated by t-test. c. The browser plot of somatic TRIs (chr7:121603262-121603506) illustrates the following epigenomic signals: (1) H3K27ac in the A673 cell line, (2) H3K27ac in the SK-N-MC cell line, (3) DNase-seq in the A673 cell line, (4) DNase-seq in the SK-N-MC cell line. Additionally,it displays the ENCODE-annotated gene transcript (5), Candidate Cis-Regulatory Elements (cCREs) predicted by ENCODE (6), and H3K27ac marks identified by ENCODE (7). The plot concludes with the CTTC motif of the repeat. An enlarged panel below outlines a detailed sequence overview of various clusters, with vertical labels indicating cluster and cancer type information. d. The browser plot of somatic TRIs (chr1:202204213-202204518) illustrates the following epigenomic signals: (8) DNase-seq in the H7 cell line, (9) DNase-seq in the A549 cell line, (10) H3K27ac in the RWPE1 cell line, (11) H3K27ac in the A549 cell line. Additionally,it displays the ENCODE-annotated gene transcript (5), Candidate Cis-Regulatory Elements (cCREs) predicted by ENCODE (6), and H3K27ac marks identified by ENCODE (7). The plot concludes with the CA motif of the repeat. An enlarged panel below outlines a detailed sequence overview of various clusters, with vertical labels indicating cluster and cancer type information. e. The browser plot of somatic TRIs (chr1:202204213-202204518) illustrates the following epigenomic signals: (12) H3K27ac in the RWPE1 cell line, (13) H3K27ac in the H9 cell line, (14) DNase-seq in the RWPE1 cell line, (15) DNase-seq in the H9 cell line. Additionally,it displays the ENCODE-annotated gene transcript (5), Candidate Cis-Regulatory Elements (cCREs) predicted by ENCODE (6), and H3K27ac marks identified by ENCODE (7). The plot concludes with the TGGTGA motif of the repeat. An enlarged panel below outlines a detailed sequence overview of various clusters, with vertical labels indicating cluster and cancer type information. f. Frequency of somatic TRI occurrence around PTP genes across cancer types. Orange: with TRI; Gray: without TRI. g. Oncoplot of samples associated with the PTP Family somatic TRI windows. The bar charts on the top and right show the counts, while the percentages on the left indicate the proportion among all samples.

**Figure S11.**
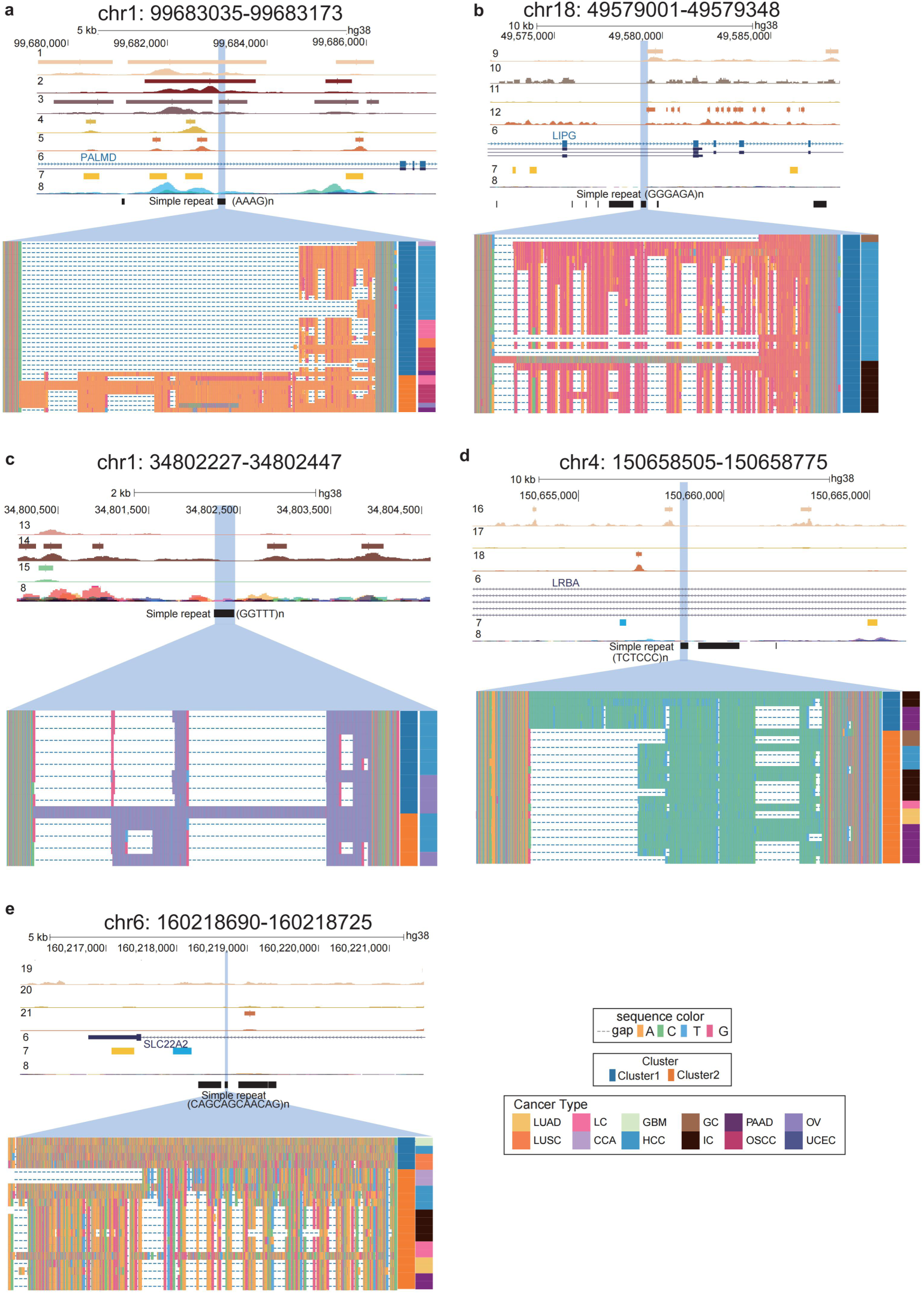
Display of representative somatic TRI events. a. The browser plot of somatic TRIs (chr1:99683035-99683173) with significantly high frequency in HCC, OSCC and LC illustrates the following epigenomic signals: (1) H3K27ac in the HeLa-S3 cell line, (2) H3K27ac in the SJSA1 cell line, (3) H3K27ac in the ACC112 cell line, (4) DNase-seq in the HeLa-S3 cell line, and (5) DNase-seq in the SJSA1 cell line. Additionally, it displays the ENCODE-annotated gene transcript (6), Candidate Cis-Regulatory Elements (cCREs) predicted by ENCODE (7), and H3K27ac marks identified by ENCODE (8). The plot concludes with the AAAG motif of the repeat. An enlarged panel below outlines a detailed sequence overview of various clusters, with vertical labels indicating cluster and cancer type information. b. The browser plot of somatic TRIs (chr18:49579001-49579348) with significantly high frequency in HCC and IC illustrates the following epigenomic signals: (9) H3K27ac in the Caco-2 cell line, (10) H3K27ac in the Karpas-422 cell line, (11) DNase-seq in the Caco-2 cell line, (12) DNase-seq in the Karpas-422 cell line, it displays the ENCODE-annotated gene transcript (6), Candidate Cis-Regulatory Elements (cCREs) predicted by ENCODE (7), and H3K27ac marks identified by ENCODE (8). The plot concludes with the GGGAGA motif of the repeat. An enlarged panel below outlines a detailed sequence overview of various clusters, with vertical labels indicating cluster and cancer type information. c. The browser plot of somatic TRIs (chr1:34802227-34802447) with significantly high frequency in HCC and OV illustrates the following epigenomic signals: (13) H3K27ac in the HFFc6 cell line, (14) DNase-seq in the K562 cell line, (15) DNase-seq in the HFFc6 cell line, and H3K27ac marks identified by ENCODE (8). The plot concludes with the GGTTT motif of the repeat. An enlarged panel below outlines a detailed sequence overview of various clusters, with vertical labels indicating cluster and cancer type information. d. The browser plot of somatic TRIs (chr4:150658505-150658775) with significantly high frequency in IC and PAAD illustrates the following epigenomic signals: (16) H3K27ac in the H9 cell line, (17) DNase-seq in the H9 cell line, (18) DNase-seq in the KBM-7 cell line, it displays the ENCODE-annotated gene transcript (6), Candidate Cis-Regulatory Elements (cCREs) predicted by ENCODE (7), and H3K27ac marks identified by ENCODE (8). The plot concludes with the GGTTT motif of the repeat. An enlarged panel below outlines a detailed sequence overview of various clusters, with vertical labels indicating cluster and cancer type information. e. The browser plot of somatic TRIs (chr6:160218690-160218725) with significantly high frequency in IC and PAAD illustrates the following epigenomic signals: (19) H3K27ac in the WTC11 cell line, (20) DNase-seq in the WTC11 cell line, (21) DNase-seq in the RCC cell line. Additionally, it displays the ENCODE-annotated gene transcript (6), Candidate Cis-Regulatory Elements (cCREs) predicted by ENCODE (7), and H3K27ac marks identified by ENCODE (8). The plot concludes with the CAGCAGCAACAG motif of the repeat. An enlarged panel below outlines a detailed sequence overview of various clusters, with vertical labels indicating cluster and cancer type information.

**Figure S12.**
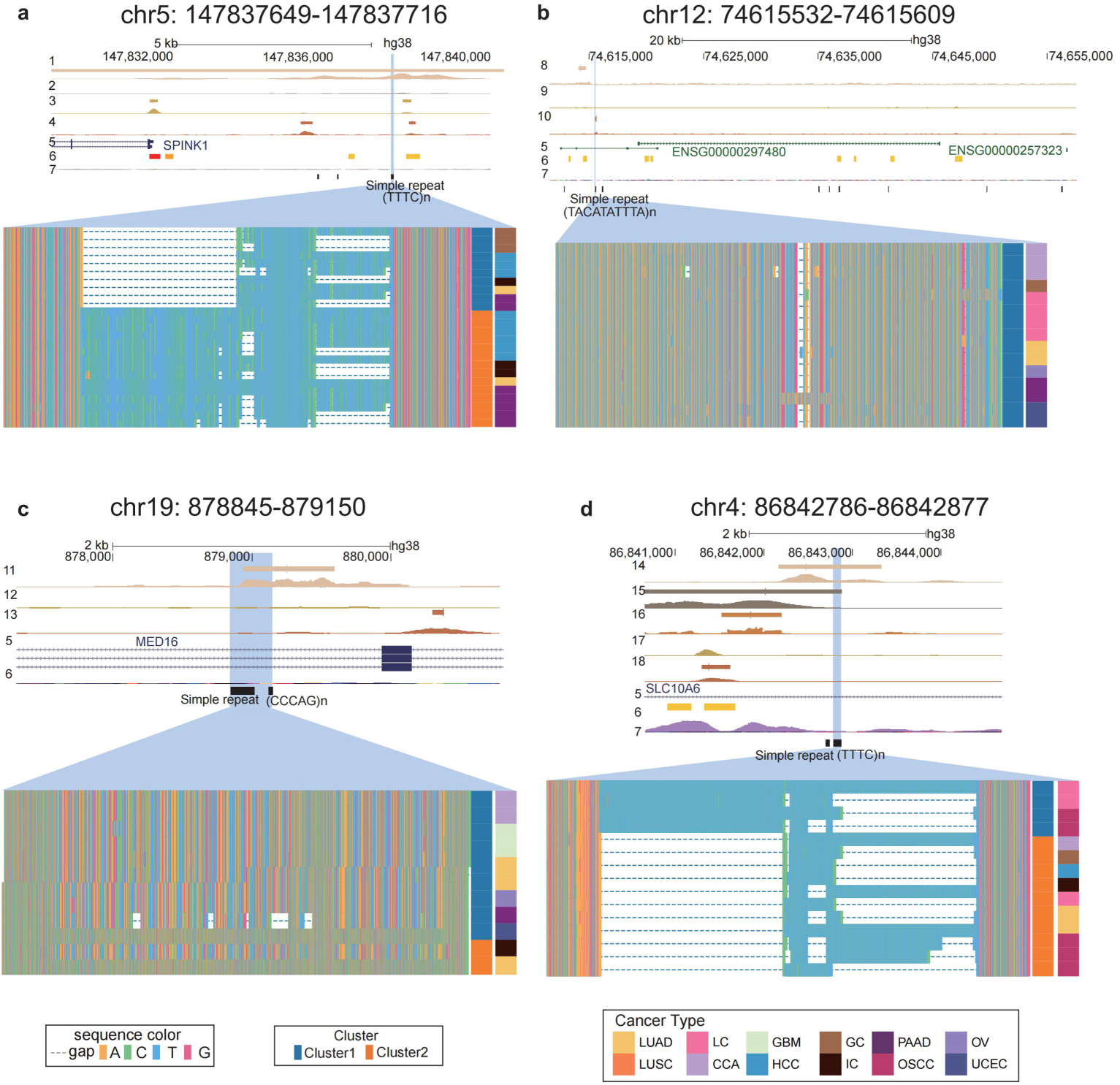
Display of representative somatic TRI events. a. The browser plot of somatic TRIs (chr5:147837649-147837716) with significantly high frequency in IC and PAAD illustrates the following epigenomic signals: (1) H3K27ac in the 22Rv1 cell line, (2) H3K27ac in the A549 cell line, (3) DNase-seq in the Caco-2 cell line, (4) DNase-seq in the A549 cell line. Additionally, it displays the ENCODE-annotated gene transcript (5), Candidate Cis-Regulatory Elements (cCREs) predicted by ENCODE (6), and H3K27ac marks identified by ENCODE (7). The plot concludes with the TTTC motif of the repeat. An enlarged panel below outlines a detailed sequence overview of various clusters, with vertical labels indicating cluster and cancer type information. b. The browser plot of somatic TRIs (chr12:74615532-74615609) illustrates the following epigenomic signals: (8) H3K27ac in the MCF-7 cell line, (9) DNase-seq in the MCF-7 cell line, (10) DNase-seq in the RCC 7860 cell line. Additionally, it displays the ENCODE-annotated gene transcript (5), Candidate Cis-Regulatory Elements (cCREs) predicted by ENCODE (6), and H3K27ac marks identified by ENCODE (7). The plot concludes with the TACATATTTA motif of the repeat. An enlarged panel below outlines a detailed sequence overview of various clusters, with vertical labels indicating cluster and cancer type information. c. The browser plot of somatic TRIs (chr19:878845-879150) illustrates the following epigenomic signals: (11) H3K27ac in the Caco-2 cell line, (12) DNase-seq in the Caco-2 cell line, (13) DNase-seq in the L1-S8R cell line, Additionally, it displays the ENCODE-annotated gene transcript (5), Candidate Cis-Regulatory Elements (cCREs) predicted by ENCODE (6). The plot concludes with the CCCAG motif of the repeat. An enlarged panel below outlines a detailed sequence overview of various clusters, with vertical labels indicating cluster and cancer type information. d. The browser plot of somatic TRIs (chr4:86842786-86842877) illustrates the following epigenomic signals: (14) H3K27ac in the RWPE1 cell line, (15) H3K27ac in the RWPE2 cell line, (16) H3K27ac in the MCF 10A cell line, (17) DNase-seq in the RWPE2 cell line, and (18) DNase-seq in the MCF 10A cell line. Additionally, it displays the ENCODE-annotated gene transcript (5), Candidate Cis-Regulatory Elements (cCREs) predicted by ENCODE (6), and H3K27ac marks identified by ENCODE (7). The plot concludes with the TTTC motif of the repeat. An enlarged panel below outlines a detailed sequence overview of various clusters, with vertical labels indicating cluster and cancer type information.

**Figure S13.**
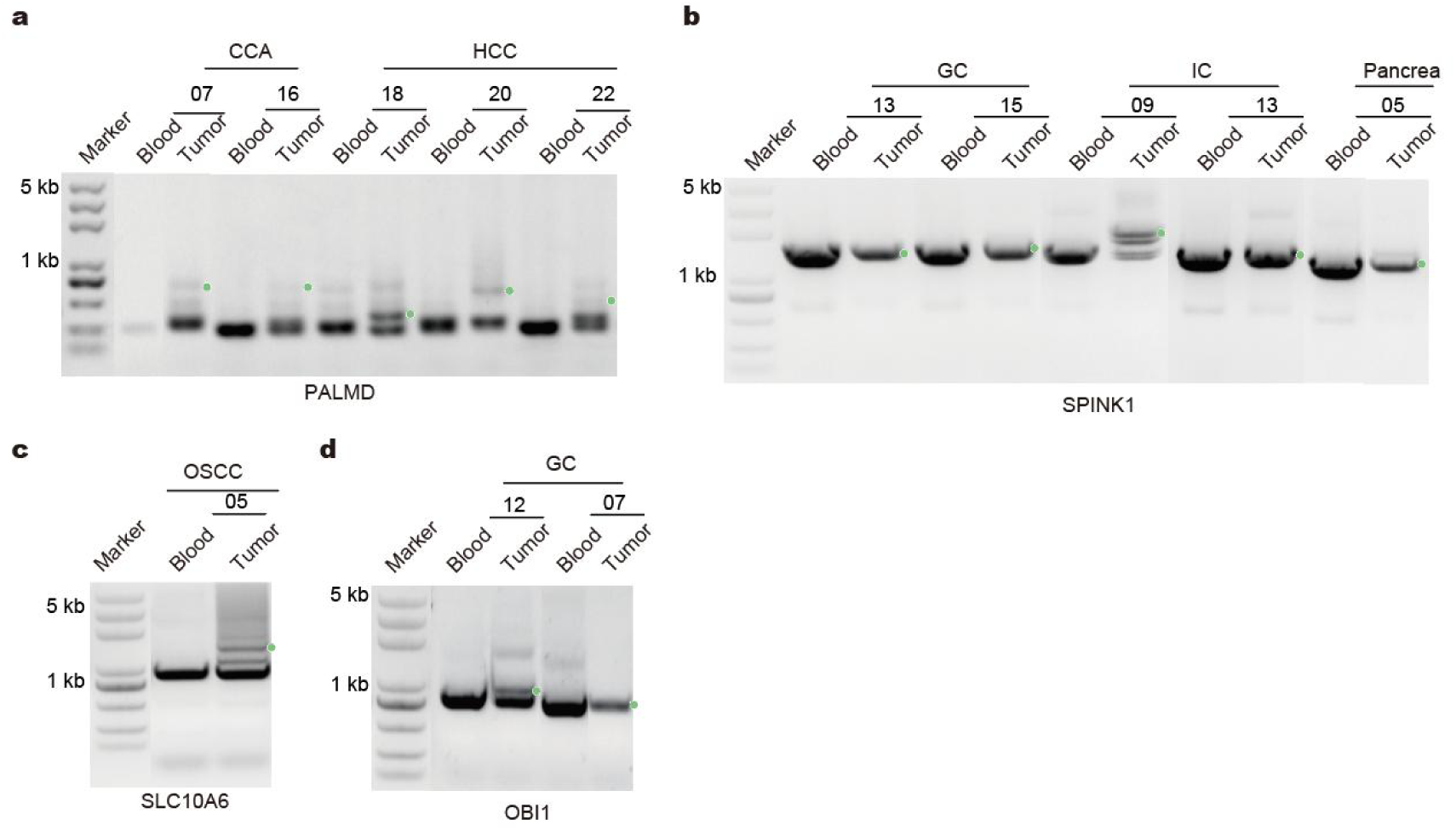
Validation of clinical samples for TRI events. a. Detecting somatic TRI events harbored by *PALMD* through PCR in clinical samples, blood DNA as a control. b. Detecting somatic TRI events harbored by *SPINK1* through PCR in clinical samples, blood DNA as a control. c. Detecting somatic TRI events harbored by *SLC10A6* through PCR in clinical samples, blood DNA as a control. d. Detecting somatic TRI events harbored by *OBI1* through PCR in clinical samples, blood DNA as a control.

**Figure S14.**
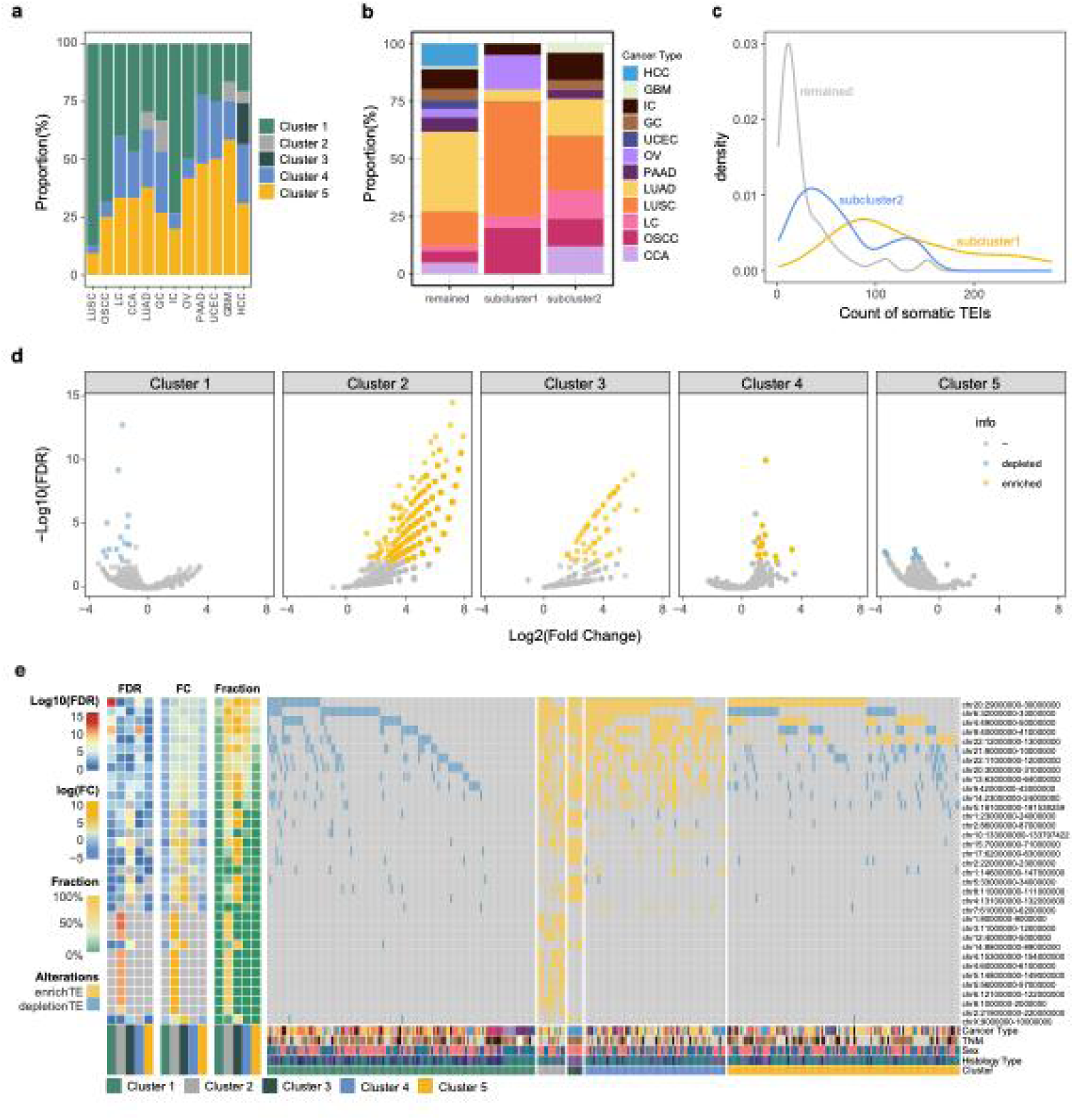
The characteristic of somatic TEI clusters. a. Distribution of the 5 clusters of somatic TEI patterns in 12 cancer types. b. Distribution of the 12 cancer types in subclusters of Cluster 1. ‘other’ presents samples out of subcluster1 and subcluster2. c. Density plot showing the distribution of somatic transposable element insertions (TEIs) in Cluster 1, separated into subcluster1 (yellow), subcluster2 (blue), and the remaining Cluster 1 samples (gray). The x-axis represents the number of somatic TEIs. d. Volcano plot of significant enriched/depleted somatic TEI windows (1Mbp) between samples in a cluster and samples not in the cluster (Fisher’s exact test, p value < 0.05). e. Oncoplot of top 10 enriched/depleted somatic TEI windows of each cluster (FDR < 0.05). The color of the left heatmap represents the significance level (Fisher’s exact test, log10(FDR)) of enrichment or depletion of a specific cluster in a specific 1Mb window. The color of the middle heatmap represents the log2(Fold Change) of enrichment or depletion of a specific cluster in a specific 1Mb window. The color of the middle heatmap represents the log2(Fold Change) of enrichment or depletion of a specific cluster in a specific 1Mb window. The color of the right heatmap represents the fraction of samples harbored somatic TEIs in the specific 1Mb window.

**Figure S15.**
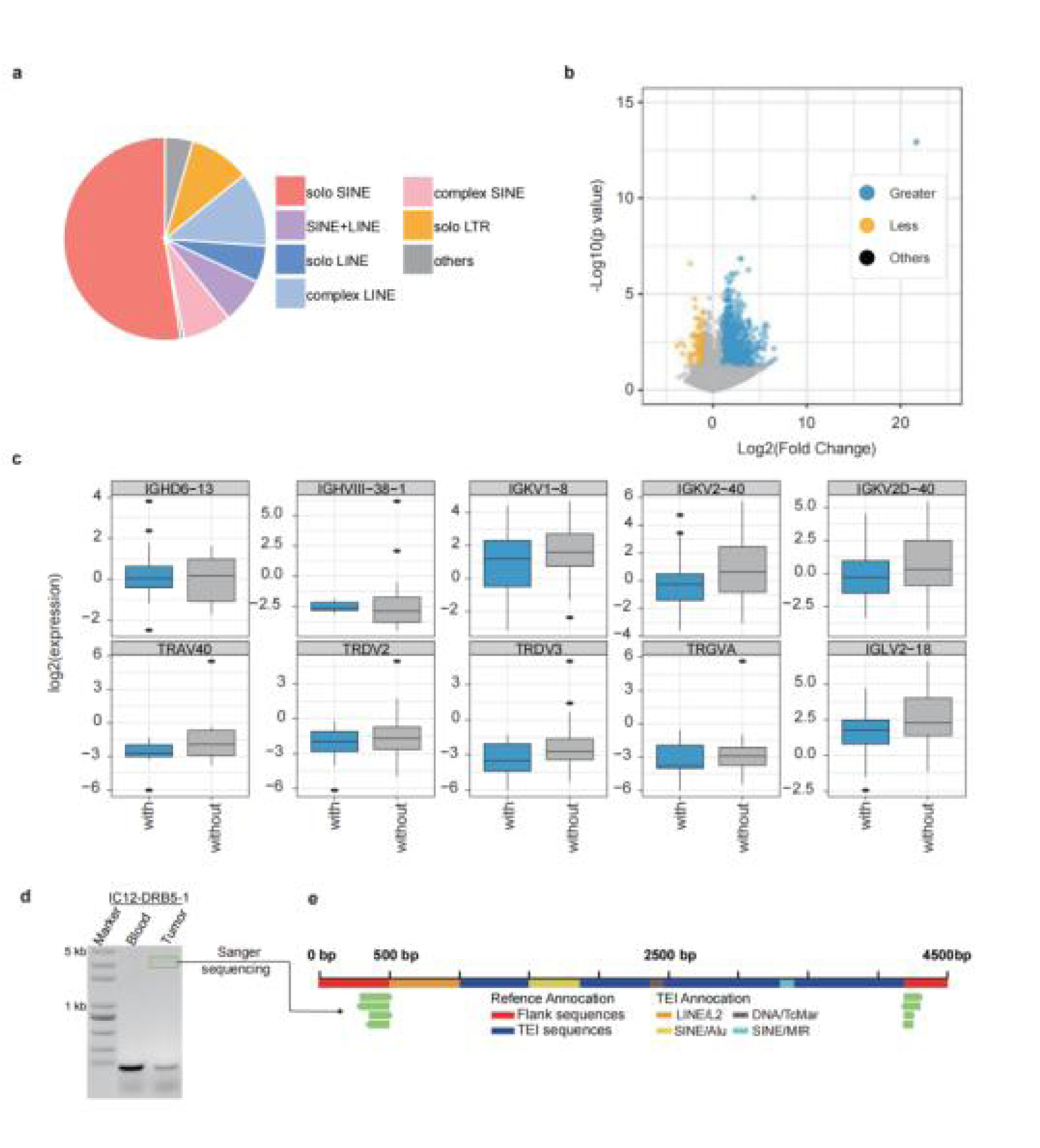
The proportion of different types of TEIs inserted in MHC II gene family, and differentially expressed genes related to these somatic TEIs. a. Pie chart of somatic TEI type distribution. ‘solo SINE’: TEIs contained solo SINE segments. ‘complex SINE’: TEIs contained multi SINE segments and not contained other segments. ‘solo LINE’: TEIs contained solo LINE segments. ‘complex LINE’: TEIs contained multi LINE segments and not contained other segments. ‘SINE+LINE’: TEIs both contained SINE segments and LINE segments. ‘solo LTR’: TEIs consist of solo LTR segments. ‘other’: the rest type of TEI. b. Volcano plot showing the differential gene expression between LUAD with somatic TEIs and LUAD without somatic TEIs. c. Box plot of gene expression (log2) in samples with (blue) and without (gray) somatic TEIs. d. Detection of somatic TEI events via PCR: PCR analysis was performed in IC12, with blood DNA used as a control to distinguish somatic TEI events. e. Sanger sequencing visualization of TEI: The reference genome sequence incorporates both the flanking region (highlighted in red) and the TEI sequence (highlighted in blue). The TEI annotation is displayed above the reference genome for precise localization and identification.

**Figure S16.**
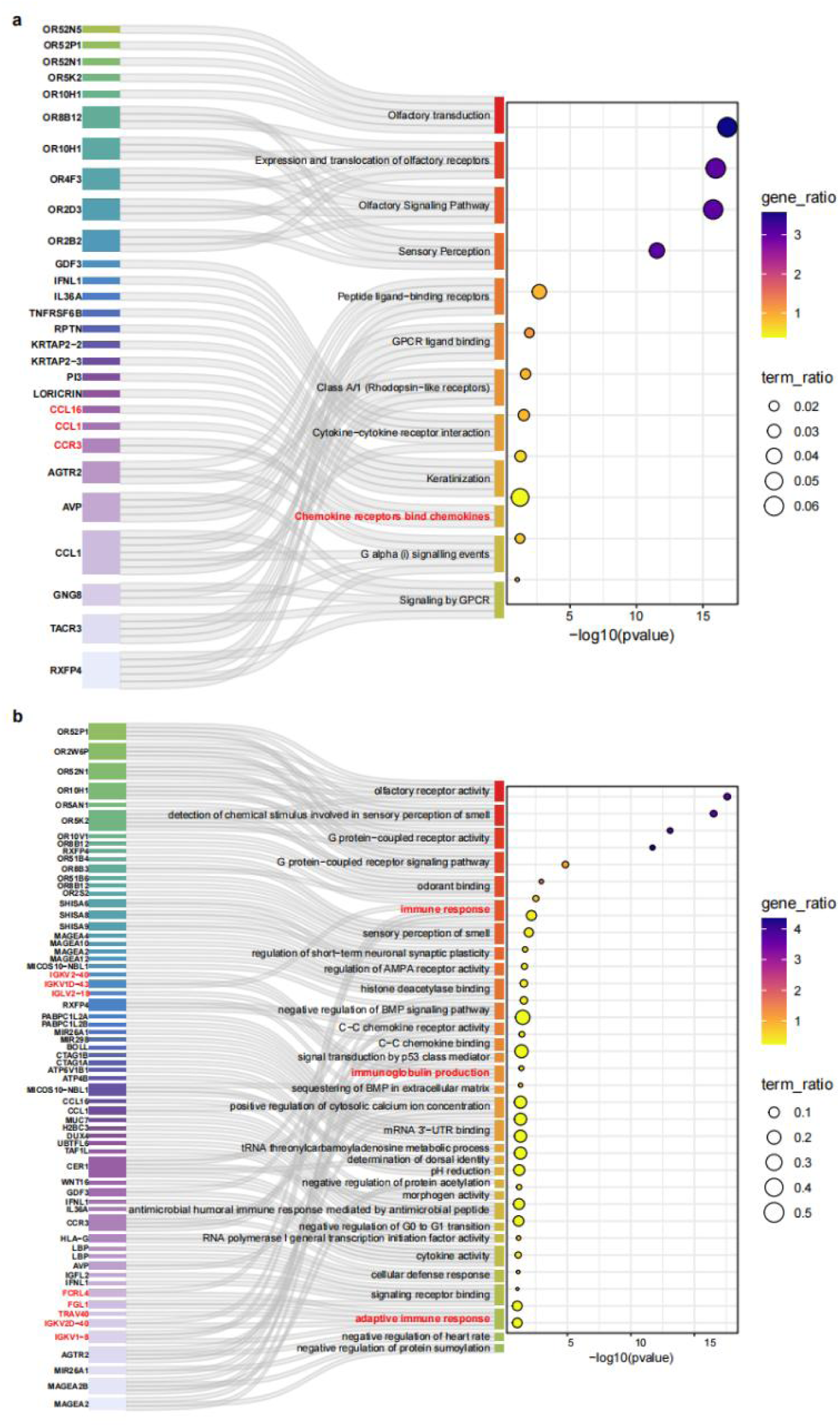
Bubble plot of pathway and GO enrichment. a. Sankey plots of KEGG pathway and corresponding genes are shown on the left, and bubble plots of enrichment results are shown on the right. The x-axis represents the −log10(p-value), y-axis lists the pathways, with the color gradient from purple to yellow representing the gene ratio. The size of the dots corresponds to the term ratio, with larger dots representing a greater magnitude of change. The connecting lines represent relationships or pathways between the genes. b. Sankey plots of GO biological process and corresponding genes are shown on the left, and bubble plots of enrichment results are shown on the right. The x-axis represents the −log10(p-value), y-axis lists the pathways, with the color gradient from purple to yellow representing the gene ratio. The size of the dots corresponds to the term ratio, with larger dots representing a greater magnitude of change. The connecting lines represent relationships or pathways between the genes.

**Figure S17.**
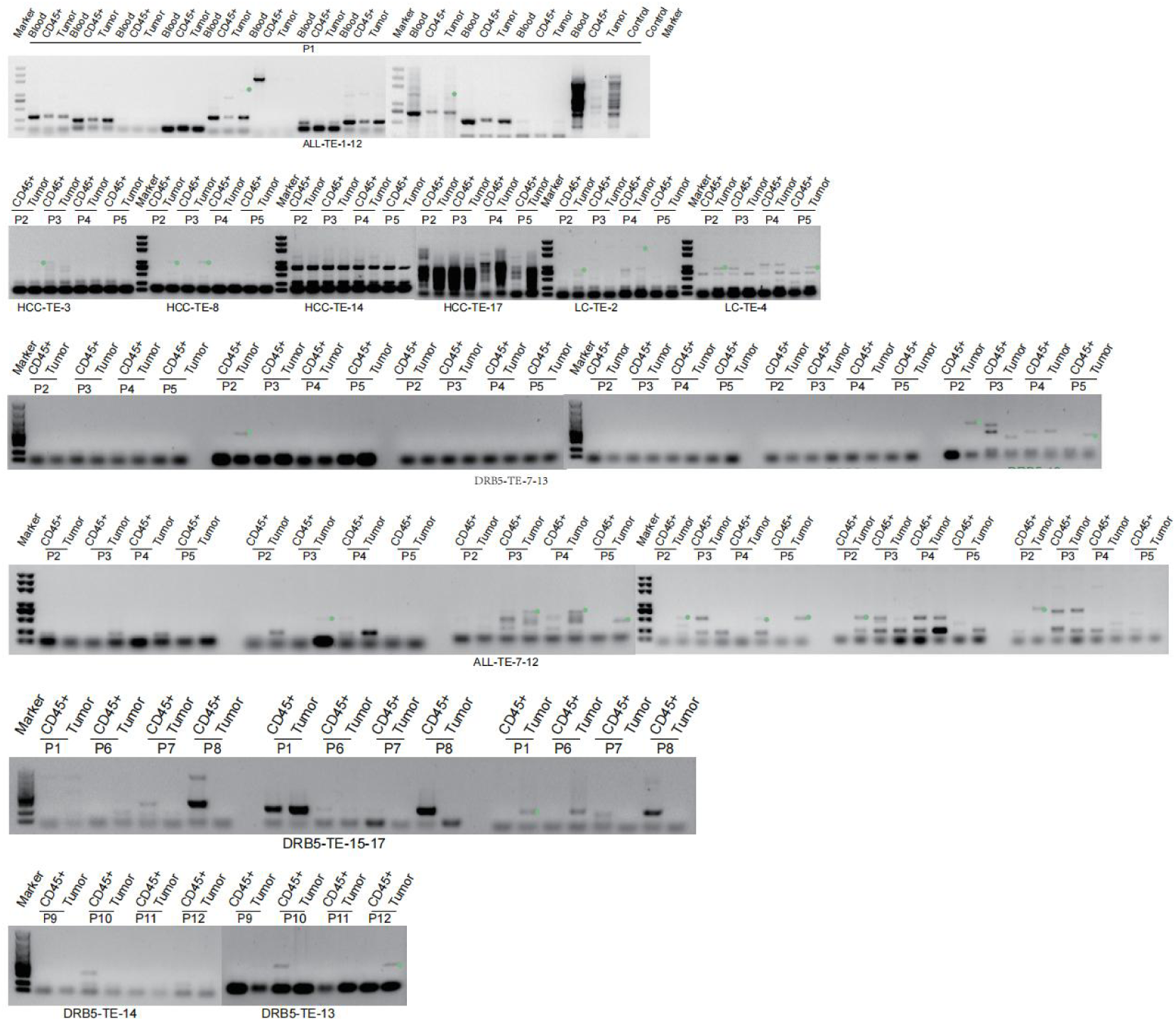
DetectingTEI events through PCR in additional fresh clinical samples after CD45+ cell sorting. The green dots mark the TEI events that were only amplified in the tumor.

**Figure S18.**
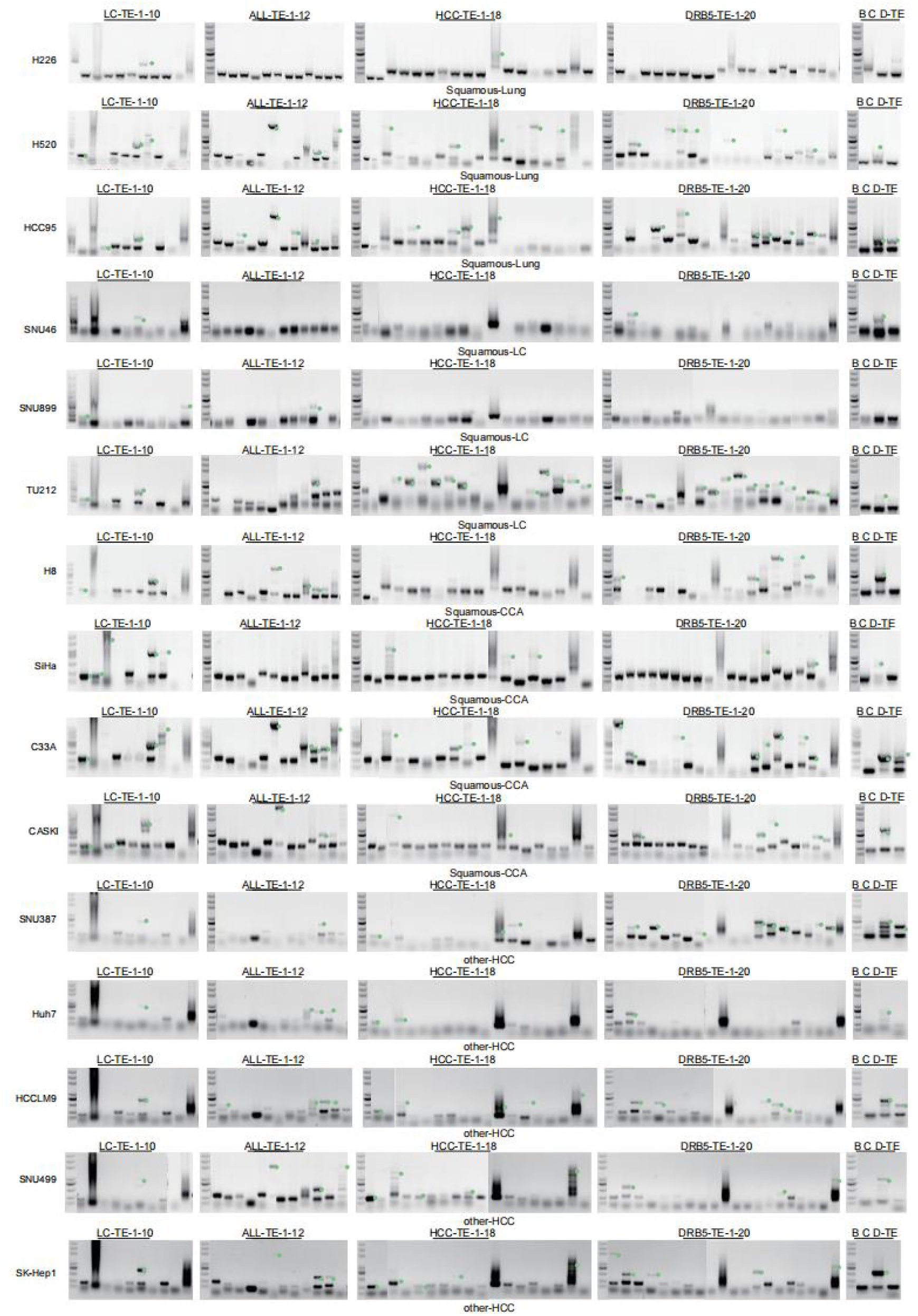
Detecting TEI events through PCR in tumor cell lines. The green dots mark the potential TEI events.

**Figure S19.**
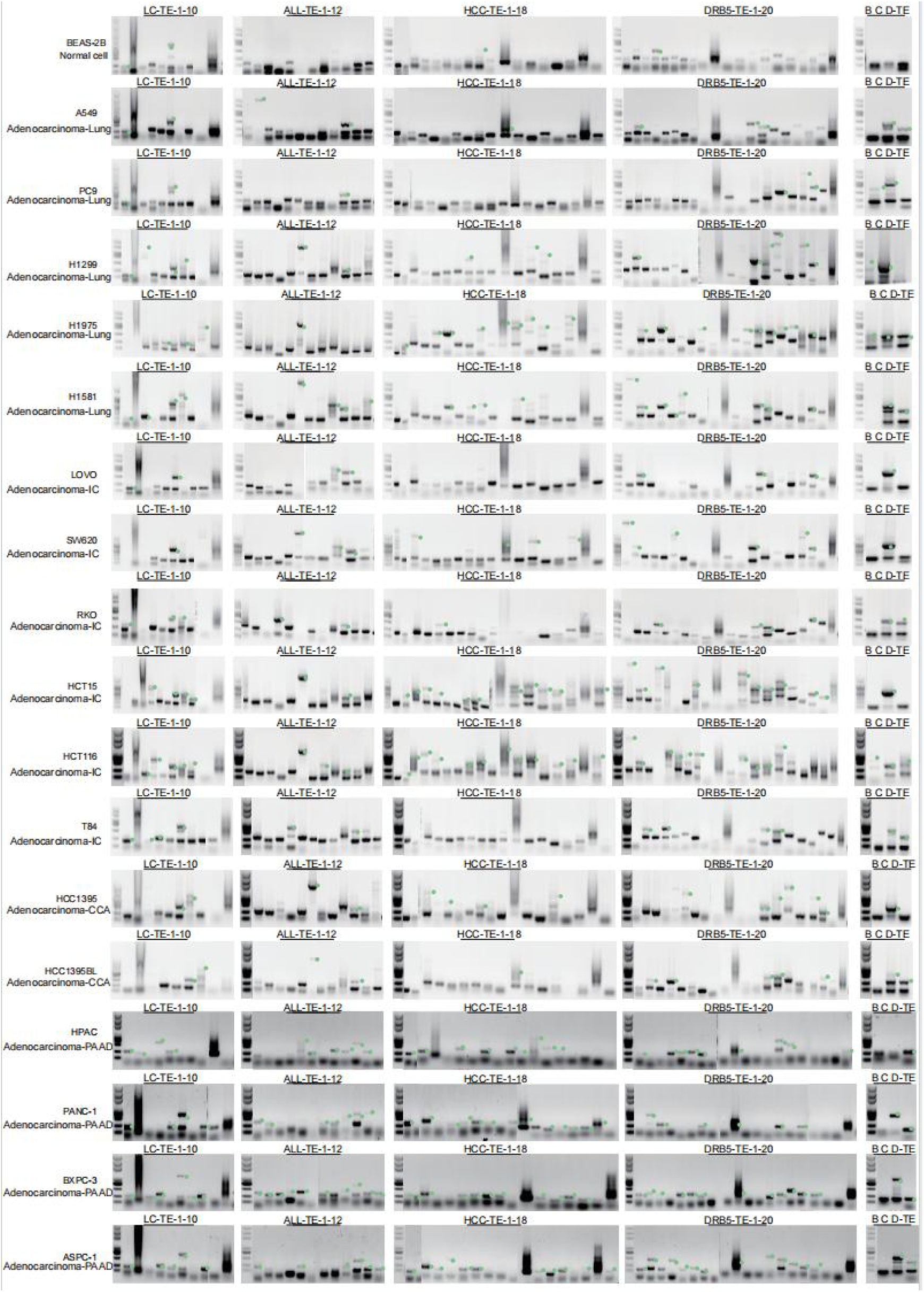
Detecting TEI events through PCR in tumor cell lines. The green dots mark the potential TEI events.

**Figure S20.**
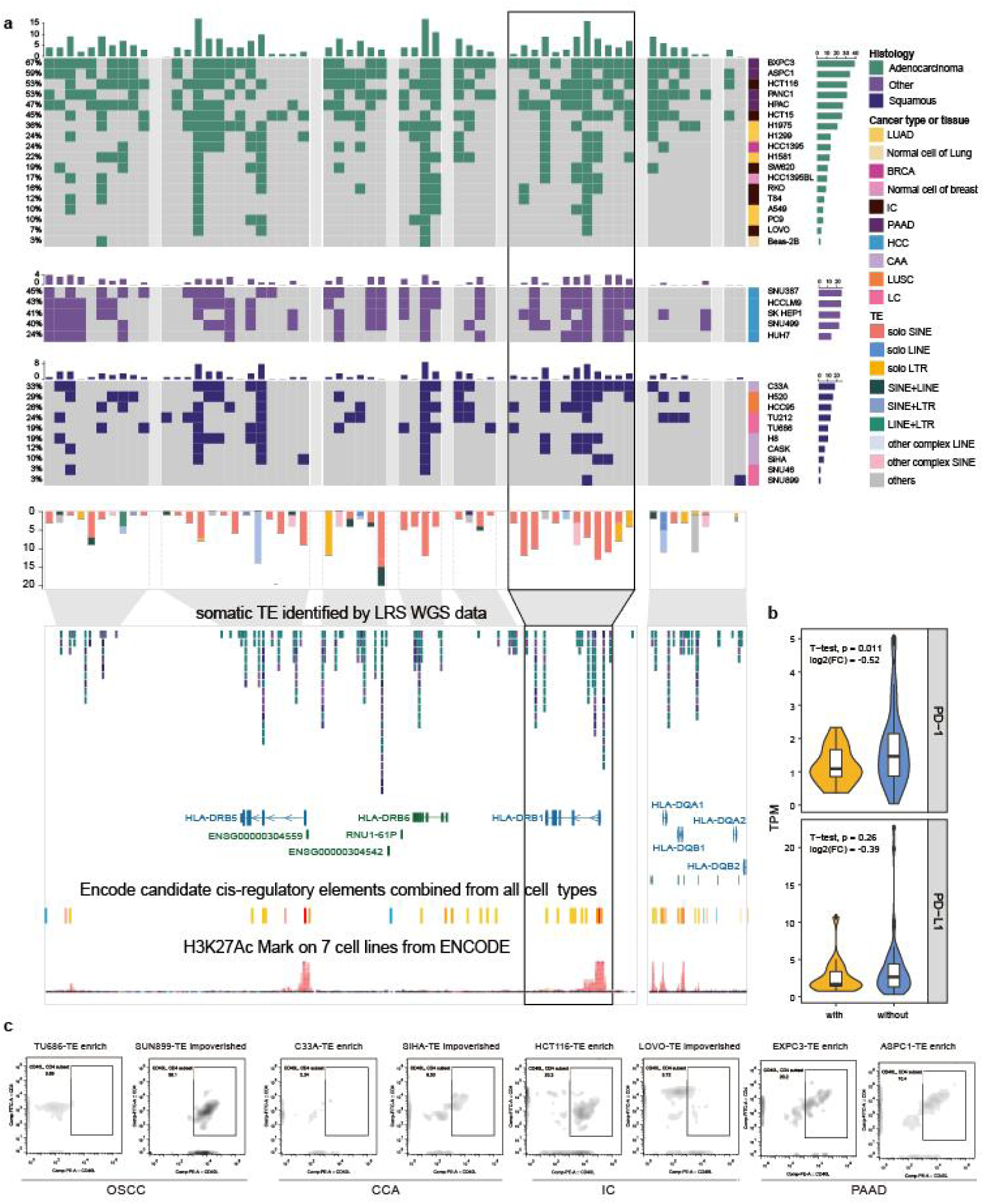
Statistics of TEI events in tumor cell lines and Flow sytometric analysis. a. Statistics of TEI events in tumor cell lines. Each column represents a TEI site, and each row represents a cell line. The first three tracks correspond to cell lines from adenocarcinoma, other types, and squamous, respectively. The fourth track summarizes TEI events from 33 cell lines in the region chr6:32,000,000-33,000,000, with different colors indicating the categories of these TEI events. The lower enlarged panel displays a browser plot showing ENCODE-annotated gene transcripts, cCREs predicted by ENCODE, and epigenomic signals around the somatic TEIs within the MHC II gene family. b. Violin plot showing the expression level of PD-1 and PD-L1 in LUAD samples harboring somatic TEIs within the HLA-DRB1 genebody and its ±5kb flanking regions (‘with’), compare to LUAD samples not harboring somatic TEIs within these regions (‘without’). c. Flow cytometric analyses of CD40L mobilization in CD4+ T cells after 24 hours of in vitro co-culture with tumor cells.

