## Supplementary Note for "Somatic repetitive element insertions Define New Biomarkers in Pan-cancer genome"

**Methylation status of CpG sites inserted by TRIs.**

To explore the role of somatic TRI in gene regulation, we analyzed 152 sets of High-Frequency TRI windows, comparing somatic TRI sequences with their matched germline sequences in terms of CpG counts (Supplementary Figure 1a, Methods). We found significant differences in CpGs counts between somatic and germline sequences in 62 of the somatic TRI windows (Wilcoxon rank test, FDR < 0.05, |log2(Fold Change)|>1, Figure S7d, Table S10). Specifically, 59 somatic TRI windows exhibited a significant increase in CpG counts in the somatic sequences compared to the germline sequences, while the remaining 3 windows showed a decrease (Figure S7d). For the somatic TRI windows with increased CpG counts in somatic sequences, we assessed the methylation status of the newly added CpG sites using Nanopolish (Supplementary Figure 1b, Methods). Notably, 34.17% of the newly added CpG sites had a methylation log-likelihood ratio between -1.5 and 1.5, a range which current algorithms could not reliably determine methylation status (Supplementary Figure 1c and 1d). However, for the remaining 65.83% of newly added CpG sites, we successfully determined their methylation status. Of these, 43.47% were methylated, while 22.36% were unmethylated. These findings aligned with previous reports {Depienne, 2021 #461} and suggested that somatic TRIs contributed to the introduction of new methylation sites by adding CpG sites.

We found 18 out of 62 somatic TRI windows that preferentially introduced methylated CpG sites through insertion events. (Wilcoxon rank test, p-value ≤ 0.05) (Figure S7e, Table S11). Notably, the somatic insertion within the somatic TRI window at chr6:64,295,338-64,295,575, located in the intron of the *EYS* gene, exhibited the highest proportion of newly methylated CpG sites, with a median of 82.14% (59.02% ~ 94.12%). This TRI event resides within a signaling region marked by H3K27ac in the PC-3 cell line, a prostate cancer model, and is adjacent to an ELF-1 binding site (Figure S7f), suggesting a potential role in gene expression regulation. This somatic TRI was detected in 3.34% of pan-cancer patients (10/299), including six LUAD patients. The somatic TRI within this region was categorized into long fragment amplification (5997–6949 bp) and short fragment amplification (1361–3787 bp). However, in the LUAD patients, somatic TRIs of varying lengths were observed, which did not conform to a single category (Figure S7g, Figure S7h). In 80% of these samples, the TRI sequence was dominated by amplification of the repetitive GAA motif, while the remaining 20% showed a combination of 'Expansion & Motif-Change'. A detailed analysis of one sample (Lung120) (Figure S7i) revealed a denser insertion of somatic CpG sites, with 92.86% of the newly inserted CpG sites exhibiting methylation. These findings are consistent with previous reports suggesting that somatic TRI events may introduce novel methylation sites via CpG insertion.

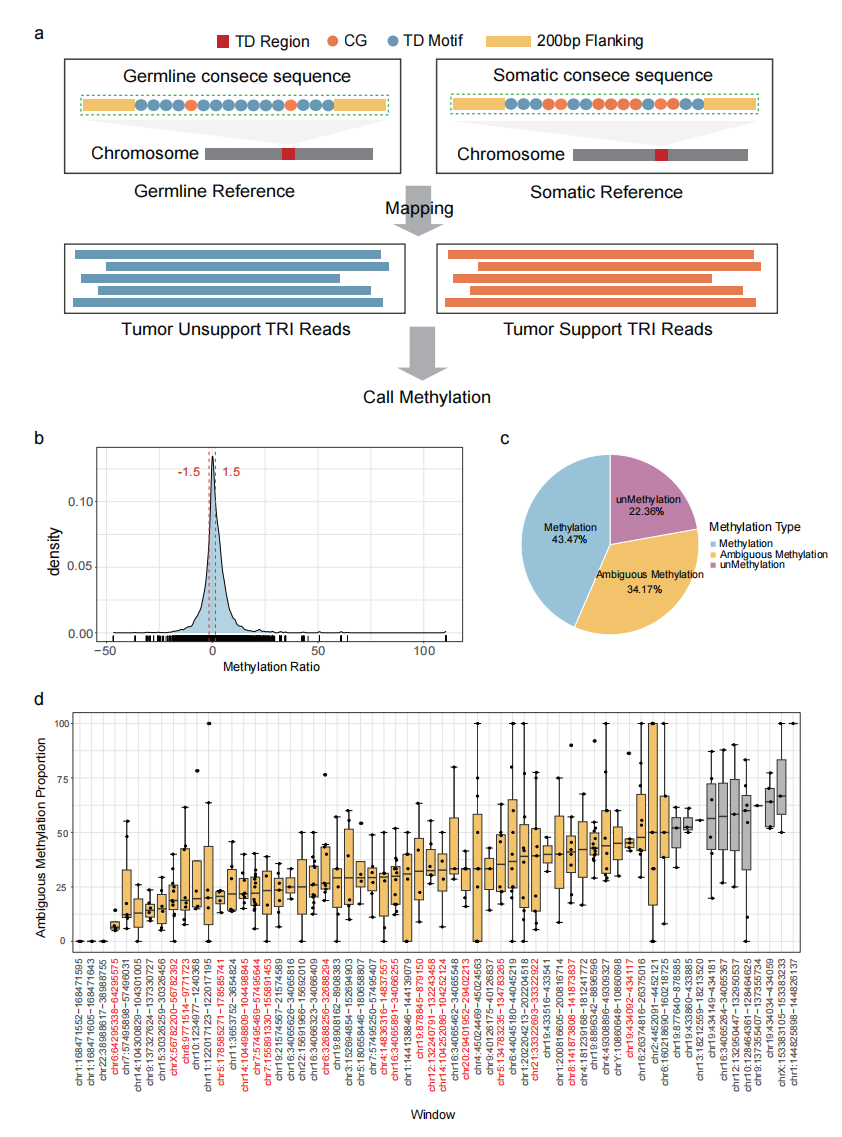

**Supplementary Figure 1. Overview of Methylation in Somatic TRIs with Significant Changes in CpG Counts.**

a. Flowchart depicting the process for methylation calling in somatic TRI sequences.
b. Distribution plot showing the methylation ratio for TRIs with significant changes in CpG counts.
c. Proportion of methylation status for soamtic TRIs with significant CpG changes. Categories include "Methylation" (somatic CpG sites detected as methylated), "Ambiguous Methylation" (unable to determine methylation status), and "Unmethylation" (somatic CpG sites detected as unmethylated).
d. Box plot displaying the proportion of samples with Ambiguous methylation status, ordered by median from smallest to largest. Each dot represents the proportion of samples with Ambiguous CpG methylation status in each TRI window. Gray: Median proportion of ambiguous methylation status greater than 50%; Window marker in red: Methylation levels were significantly elevated (wilcoxon-test p<0.05).

**The sequence features of TEIs.**

We analyzed the known sequence characteristics of somatic TEIs in human genomes. It is well-established that the length of the poly A tail in SINEs and LINEs plays a key role in their functionality and is a critical marker of their retrotranscription activity. Our findings showed that both somatic solo SINEs and somatic solo LINEs were more likely to carry poly A tails compared to their complex counterparts (Chi-square test, p = 3.5e-08 for somatic SINEs, p = 1.5e-05 for somatic LINEs, Supplementary Figure 2a). These findings suggested that the formation of somatic complex TEIs may involve a more disordered process compared to somatic solo TEIs, potentially leading to the loss of poly A tails. However, the functional impact of somatic TEIs that lack poly A tails remains uncertain and requires further investigation.

One well-known feature of somatic LINE insertions is 5’ truncation [citation]. By using LRS, we were able to analyze the integrity of somatic LINE insertions across the entire somatic TEI sequence. Remarkably, we found no fully intact somatic LINE insertions. Instead, 22.7% of somatic LINE showed 5’ truncations, and only 0.2% exhibited 3’ truncations (Supplementary Figure 2b). Additionally, we identified another category (77.1%), which we termed "5' dominant truncation", where the 5' truncation was shorter than the 3' truncation. This unique pattern further emphasized the tendency of somatic LINEs to favor 5’ truncation during insertion.

Although previous studies had indicated that some SINE insertions in the human genome were incomplete, detailed investigations into this phenomenon remained limited [citation]. Our analysis revealed that the most common form of SINE insertion was ‘5' dominant truncation’ (42.5%), followed by 3' truncations (31.5%). Fully intact SINE insertions constituted 14.1%, with 5' truncations accounting for the remaining 12.0% (Supplementary Figure 2c). Notably, we observed a significant difference in the distribution of SINE insertion integrity between high-burden tumor samples and other tumor samples (Supplementary Figure 2d). In high-burden samples, the proportions of full-length SINEs and 5' truncations were significantly higher than in other samples (Wilcoxon test, p=3.7e-06 for full length, p=0.00061 for 5' truncation) (Supplementary Figure 2e). This suggested that SINEs in high-burden samples exhibited higher activity and retained a greater proportion of fully functional insertions.

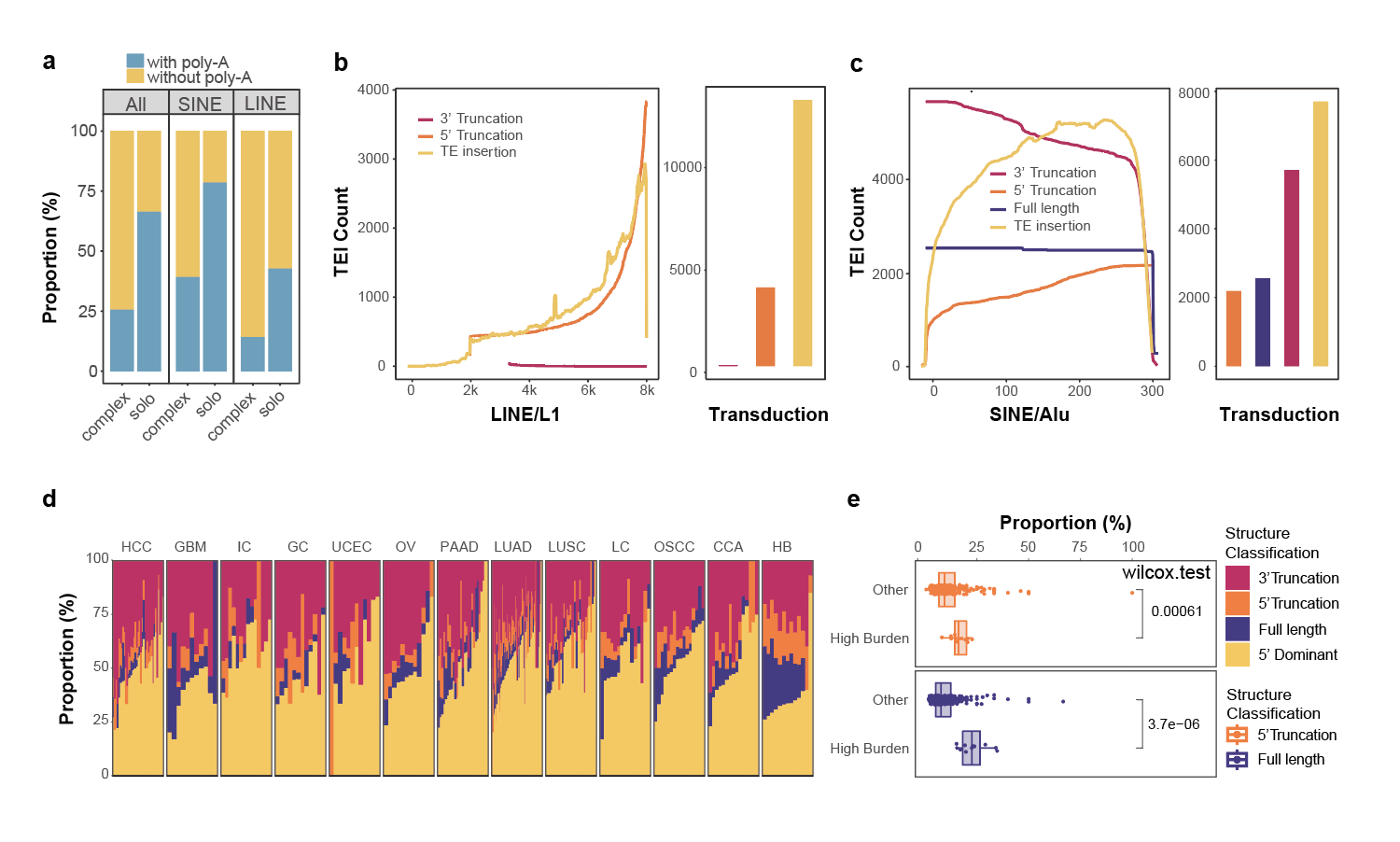

**Supplementary Figure 2. The characteristic of the somatic TEI sequences.**

a. Proportions of somatic TEIs with (blue) and without (yellow) polyA tails. ‘All’: all somatic TEIs, ‘SINE’: somatic TEIs contained SINE segment, ‘LINE’: somatic TEIs contained LINE segments.
b. The lengths of all somatic TEI contains LINE segnments are illustrated as a coverage plot over the schematic representation of a canonical solo-L1 sequence. TEIs are categorized into the following types: Full Length (purple): the TEI sequence aligned completely with the template sequence without gaps at both ends; 5' Truncation (orange): the TEI sequence aligned fully with the 3' end of the template; 3' Truncation (amaranth): the TEI sequence aligned fully with the 5' end of the template; 5’ dominant (yellow): The TEI sequence aligns with the template with gaps at both the 3' and 5' ends. The bar chart on the right shows the number of each type of somatic TEI contains LINE segnments.
c. The lengths of all somatic TEI contains SINE are illustrated as a coverage plot over the schematic representation of a canonical solo-Alu sequence. The bar chart on the right shows the number of each type of TEI.
d. A bar chart illustrating the distribution of different TEI truncation types in various cancer types, with each bar representing a sample and different colors indicating distinct truncation types. Full Length (purple): The TEI sequence aligns completely with the template sequence without gaps at both ends; 5' Truncation (orange): The sequence aligns fully with the 3' end of the template; 3' Truncation (amaranth): The sequence aligns fully with the 5' end of the template; 5’ Dominant (yellow): The TEI sequence aligns with the template strand with gaps at both the 3' and 5' ends.
e. Box plot shows that Full Length and 5' Truncation in high-burden samples are significantly higher than in other cancer types with p-values calculated using the Wilcoxon test.

**Analysis of the Association Between L1 Insertion Rate and Genomic Features**
We conducted a genome-wide analysis to examine the distribution of somatic L1 insertions (somatic TEIs containing L1 segments) in cancer genomes, revealing significant variation in L1 insertions rates (Supplementary Figure 3a). To understand the factors driving this variation, we investigated the relationship between somatic L1 insertion rates and various genomic features. First, we explored whether the presence of L1 endonuclease target site motifs could explain the observed distribution. Using a negative binomial regression model, we assessed the impact of multiple overlapping genomic variables (PMID: 28753428). Our analysis showed that regions with matching motifs had a 2.2-fold higher abundance of L1 events compared to regions without matching motifs (95% confidence interval: 1.6–2.25, Supplementary Figure 3a). We also observed a strong association between L1 events and DNA replication timing. The most recently replicated quarter of the genome exhibited 5.6 times more L1 insertions than the earliest replicated quarter (95% confidence interval: 5.4–5.8, Supplementary Figures 3b and 3c). This finding aligns with a previous study (PMID: 32024998). Furthermore, recent research has shown that L1 retrotransposition is biased towards the S phase of the cell cycle (PMID: 29309036).This aligns with our findings, which suggest that regardless of the underlying mechanisms driving somatic L1 insertions, these events are most prevalent during the later stages of nuclear DNA synthesis.

Next, we investigated the relationship between somatic L1 insertion rates and chromatin accessibility, measured by DNase hypersensitivity, as well as the association with various histone marks. Our analysis revealed that somatic L1 insertions preferentially occur in heterochromatic regions, consistent with previous studies {Tubio, 2014 #130}. Specifically, somatic L1 segments preferentially inserted into closed heterochromatin regions, particularly those marked by K9-trimethylated histone H3 (H3K9me3), showing a 4.57-fold increase from low to high bins (95% confidence interval: 4.23–4.94; Supplementary Figure 3b). In contrast, somatic L1 insertions were reduced in open chromatin regions, with a 6.66-fold decrease in the DNase hypersensitivity low group (95% confidence interval: 5.89–7.54; Supplementary Figure 3b). Additionally, we found a negative correlation between somatic insertion rates and chromatin features associated with active transcription. Somatic L1 insertions were significantly reduced in active promoter regions marked by H3K4me3 (13.72-fold decrease; Supplementary Figure 3b). In highly expressed genes, the somatic L1 insertion rate was also significantly lower, showing a 7.06-fold reduction (95% confidence interval: 6.64–7.5; Supplementary Figure 3b). The reduction was even more pronounced in regions marked by H3K36me3, which are associated with transcriptional activity in the gene body and 3’ end, where somatic L1 insertion rate decreased by 10.54-fold in the highest quartile (95% confidence interval: 8.87–12.58; Supplementary Figure 3b){Barski, 2007 #131}. A similar decrease in L1 insertions was observed in active enhancer regions marked by H3K27ac, with a 10.92-fold reduction (95% confidence interval: 8.57–14.07; Supplementary Figure 3b). These findings were consistent regardless of the mechanisms underlying somatic L1 insertions, or whether the insertions were in solo or complex structures.

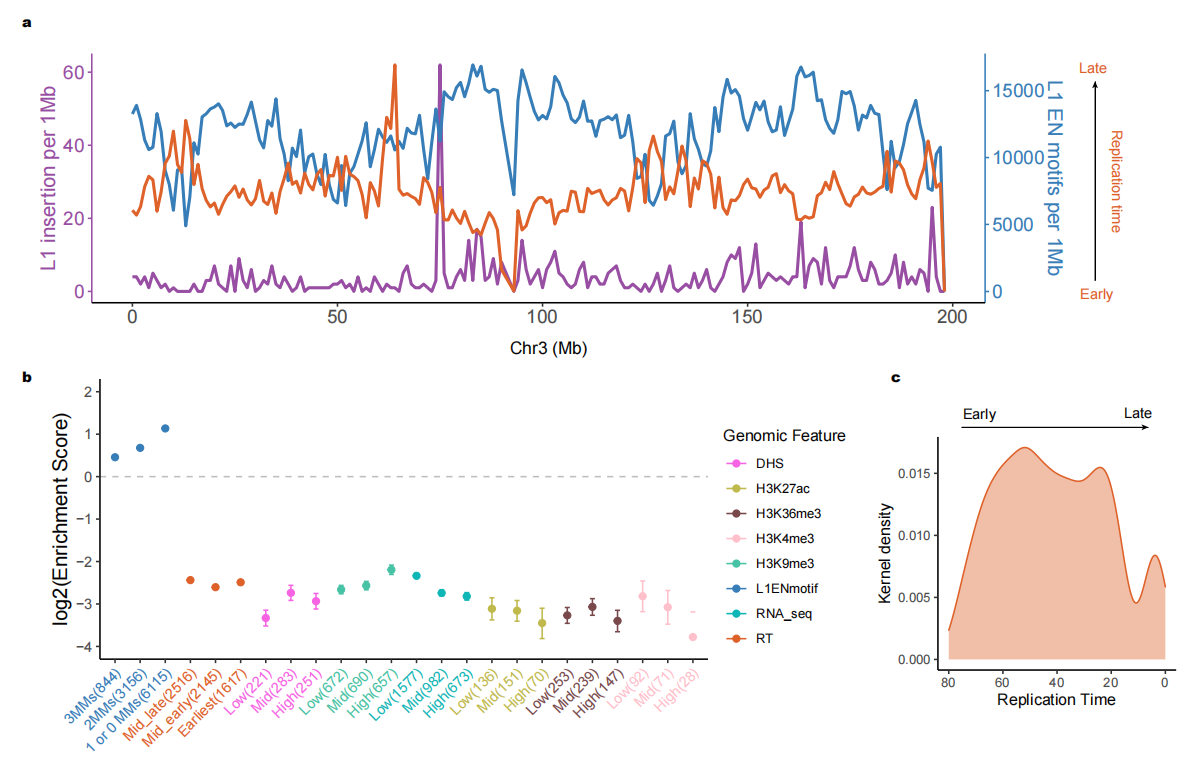

**Supplementary Figure 3. Distribution of L1 somatic insertions across the cancer genome and their relationship with genomic organization features.**

a. The frequency of L1 insertions (purple), the density of L1 endonuclease (EN) motifs (blue), and the DNA replication timing (orange) are plotted, with data aggregated into 1-megabase segments for clarity. Chromosome 3 is shown as an illustrative example.
b. The relationship between L1 insertion frequency and multiple genomic features is analyzed. Enrichment scores (represented as dots) are calculated by comparing L1 insertion rates in bins 1–3 for specific genomic attributes against bin 0, which inherently has a log-transformed enrichment score of zero and is excluded from the plot. Adjustments are made for various covariates. Features such as replication timing, DNase hypersensitivity (DHS), histone marks (H3K9me3, H3K27ac, H3K36me3, H3K4me3), expression level (RNA_seq) are included. Error bars indicate 95% confidence intervals, and the number of observations per bin is shown in parentheses.
c. The density of L1 insertions, estimated using kernel density estimation (KDE), is plotted across the spectrum of replication timing. Replication timing is quantified on a scale ranging from 80 (early replication) to 0 (late replication).
